# Pervasive and diverse collateral sensitivity profiles inform optimal strategies to limit antibiotic resistance

**DOI:** 10.1101/241075

**Authors:** Jeff Maltas, Kevin B. Wood

## Abstract

Evolved resistance to one antibiotic may be associated with “collateral” sensitivity to other drugs. Here we provide an extensive quantitative characterization of collateral effects in *Enterococcus faecalis*, a gram-positive opportunistic pathogen. By combining parallel experimental evolution with high-throughput dose-response measurements, we measure phenotypic profiles of collateral sensitivity and resistance for a total of 900 mutant-drug combinations. We find that collateral effects are pervasive but difficult to predict, as independent populations selected by the same drug can exhibit qualitatively different profiles of collateral sensitivity as well as markedly different fitness costs. Using whole-genome sequencing of evolved populations, we identified mutations in a number of known resistance determinants, including mutations in several genes previously linked with collateral sensitivity in other species. While phenotypic drug sensitivity profiles show significant diversity, they cluster into statistically similar groups characterized by selecting drugs with similar mechanisms. To exploit the statistical structure in these resistance profiles, we develop a simple mathematical model based on a stochastic control process and use it to design optimal drug policies that assign a unique drug to every possible resistance profile. Stochastic simulations reveal that these optimal drug policies outperform intuitive cycling protocols by maintaining long-term sensitivity at the expense of short-term periods of high resistance. The approach reveals a new conceptual strategy for mitigating resistance by balancing short-term inhibition of pathogen growth with infrequent use of drugs intended to steer pathogen populations to a more vulnerable future state. Experiments in laboratory populations confirm that model-inspired sequences of four drugs reduce growth and slow adaptation relative to naive protocols involving the drugs alone, in pairwise cycles, or in four-drug uniform cycles.

## INTRODUCTION

The rapid emergence of drug resistance is an urgent threat to effective treatments for bacterial infections, cancers and many viral infections (1, 2, 3, 4, 5, 6). Unfortunately, the development of novel drugs is a long and arduous process, underscoring the need for alternative approaches to forestall resistance evolution. Recent work has highlighted the promise of evolution-based strategies for optimizing and prolonging the efficacy of established drugs, including optimal dose scheduling (7, 8, 9), antimicrobial stewardship (10,11), drug cycling (12,13,14), consideration of spatial dynamics (15, 16, 17), cooperative dynamics (18, 19, 20, 21), or phenotypic resistance (22, 23, 24), and judicious use of drug combinations (25, 26, 27, 28, 29, 30, 31, 32). In a similar spirit, a number of recent studies have suggested exploiting collateral sensitivity as a means for slowing or even reversing antibiotic resistance (33, 34, 35, 36, 37, 38). Collateral evolution occurs when a population evolves resistance to a target drug while simultaneously exhibiting increased sensitivity or resistance to a different drug. From an evolutionary perspective, collateral effects are reminiscent of the trade-offs inherent when organisms are required to simultaneously adapt to different tasks, an optimization that is often surprisingly simple because it takes place on a lowdimensional phenotypic space (39, 40). If similarly tractable dynamics occur in the evolution of multi-drug resistance, systematic optimization of drug deployment has the promise to mitigate the effects of resistance.

Indeed, recent studies in bacteria have shown that the sequential (41, 42, 43, 44, 45, 38, 46) or simultaneous (47, 48) deployment of antibiotics with mutual collateral sensitivity can sometimes slow the emergence of resistance. Unfortunately, collateral profiles have also been shown to be highly heterogeneous (49, 50) and often not repeatable (51), potentially complicating the design of successful collateral sensitivity cycles. The picture that emerges is enticing, but complex; while collateral effects offer a promising new dimension for improving therapies, the design of drug cycling protocols is an extremely difficult problem that requires optimization at multiple scales, from dynamics within individual hosts to those that occur at the hospital or community scale. Despite many promising recent advances, it is not yet clear how to optimally harness collateral evolutionary effects to design drug policies, even in simplified laboratory scenarios. The problem is challenging for many reasons, including the stochastic nature of evolutionary trajectories and-at an empirical level-the relative paucity of data regarding the prevalence and repeatability of collateral sensitivity profiles in different species.

In this work, we take a step towards answering these questions by investigating how drug sequences might be used to slow resistance in a simplified, single-species bacterial population. We show that even in this idealized scenario, intuitive cycling protocols-for example, sequential application of two drugs exhibiting reciprocal collateral sensitivity-are expected to fail over long time periods, though mathematically optimized policies can maintain long-term drug sensitivity at the price of transient periods of high resistance. As a model system, we focus on *Enterococcus faecalis*, a gram-positive opportunistic bacterial pathogen. *E. faecalis* are found in the gastrointestinal tracts of humans and are implicated in numerous clinical infections, ranging from urinary tract infections to infective endocarditis, where they are responsible for between 5 and 15 percent of cases (52, 53, 54, 55, 56). For our purposes, *E. faecalis* is a convenient model species because it rapidly evolves resistance to antibiotics in the laboratory (57, 58), and fully sequenced reference genomes are available (59).

By combining parallel experimental evolution of *E. faecalis* with high-throughput dose-response measurements, we provide collateral sensitivity and resistance profiles for 60 strains evolved to 15 different antibiotics, yielding a total of 900 mutant-drug combinations. We find that cross resistance and collateral sensitivity are pervasive in drug-resistant mutants, though patterns of collateral effects can vary significantly, even for mutants evolved to the same drug. Notably, however, the sensitivity profiles cluster into groups characterized by selecting drugs from similar drug classes, indicating the existence of large scale statistical structure in the collateral sensitivity profiles. To exploit that structure, we develop a simple mathematical framework based on a Markov Decision Process (MDP) to identify optimal antibiotic policies that minimize resistance. These policies yield drug sequences that can be tuned to optimize either short-term or long-term evolutionary outcomes, and they codify the trade-offs between instantaneous drug efficacy and delayed evolutionary consequences. While clearly too simple to capture evolution in realistic clinical scenarios, the model points to new conceptual strategies for mitigating resistance by balancing short-term growth inhibition with infrequent use of drugs intended to steer pathogen populations to a more vulnerable future state.

## RESULTS

### Collateral effects are pervasive and heterogeneous

To investigate collateral drug effects in *E. faecalis*, we exposed four independent populations of strain V583 to increasing concentrations of a single drug over 8 days (approximately 60 generations) using serial passage laboratory evolution (Figure 1A, Methods). We repeated this laboratory evolution for a total of 15 antibiotics spanning a wide range of classes and mechanisms of action (Table 2). Many, but not all, of these drugs are clinically relevant for the treatment of enterococcal infections. As a control, we also evolved 4 independent populations of the ancestral V583 strain to media (BHI) alone. After approximately 60 generations, we isolated a single colony (hereafter termed a “mutant”) from each population and measured its response to all 15 drugs using replicate dose-response experiments (Figure 1B). To quantify resistance, we estimated the half maximal inhibitory concentration (IC_50_) for each mutant-drug combination using non linear least squares fitting to a Hill-like dose response function (Methods; see Figure S1 for examples). A mutant strain was defined to be collaterally sensitive if its IC_50_ had decreased by more than 3*σ_WT_* relative to the ancestral strain (*σ_WT_* is defined as the uncertainty-standard error across replicates-of the IC_50_ measured in the ancestral strain). Similarly, an increase in IC_50_ by more than 3*σ_WT_* relative to the ancestral strain corresponds to cross-resistance. As a measure of cross resistance / sensitivity, we then calculate *C* ≡ log_2_ (IC_50,*Mut*_/IC_50, *WT*_), the (log-scaled) fold change in IC_50_ of each mutant relative to wild-type (WT); values of *C* > 0 indicate cross resistance, while values of *C* < 0 indicate collateral sensitivity (Figure 1C). For each mutant, we refer to the set of C values (one for each testing drug) as its collateral sensitivity profile 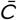.

**FIG 1.**
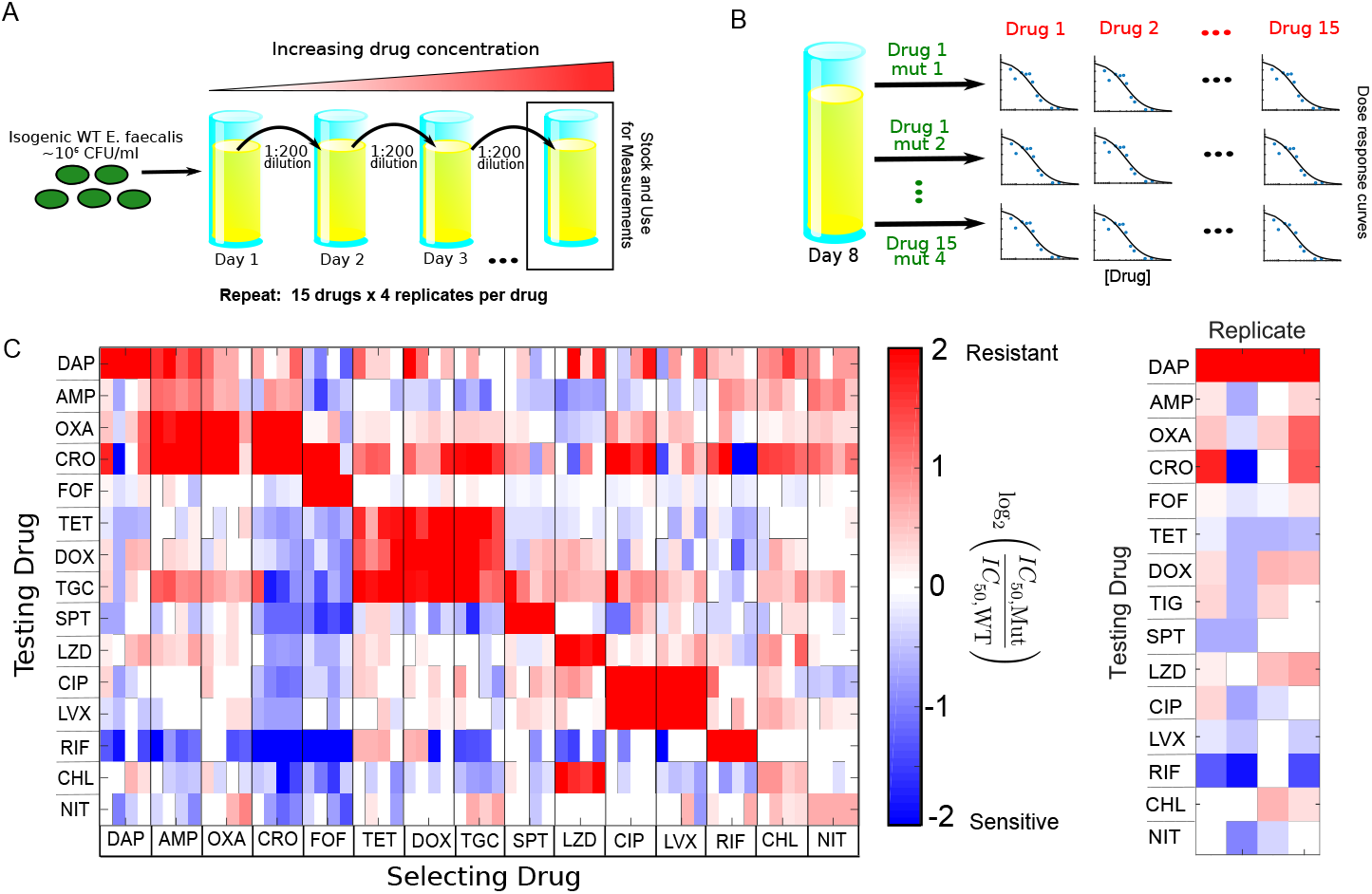
Collateral effects are pervasive and vary across parallel evolution experiments in *E. faecalis.* A. *E. faecalis* strain V583 was exposed to increasing concentrations of a single antibiotic over an 8-day serial passage experiment with daily 200-fold dilutions (≈60 generations total; see Methods). The evolution was performed in quadruplicate for each drug and repeated for a total of 15 drugs (Table 2). After 8 days, a single mutant was isolated from each population. B. The half maximal inhibitory concentration (IC_50_) for each of 15 drugs was estimated for all 60 mutants by nonlinear fitting of a dose response curve (relative OD) to a Hill-like function (Methods). C. Main panel: resistance (red) or sensitivity (blue) of each evolved mutant (horizontal axis; 15 drugs x 4 mutants per drug) to each drug (vertical axis) is quantified by the log_2_-transformed relative increase in the IC_50_ of the testing drug relative to that of wild-type (V583) cells. While the color scale ranges from a 4x decrease to a 4x increase in IC_50_, it should be noted that both resistance to the selecting drug (diagonal blocks) and collateral effects can be significantly higher. Each column of the heat map represents a collateral sensitivity profile for one mutant. Right panel: enlarged first column from main panel. Mutants isolated from replicate populations evolved to daptomcyin exhibit diverse sensitivity prof les. While all mutants are resistant to the selecting drug (daptomycin), mutants may exhibit either sensitivity or resistance to other drugs. For example, the first and last replicates exhibit cross resistance to ceftriaxone (CRO), while replicate 2 exhibits collateral sensitivity and replicate 3 shows little effect.

**TABLE 1.**
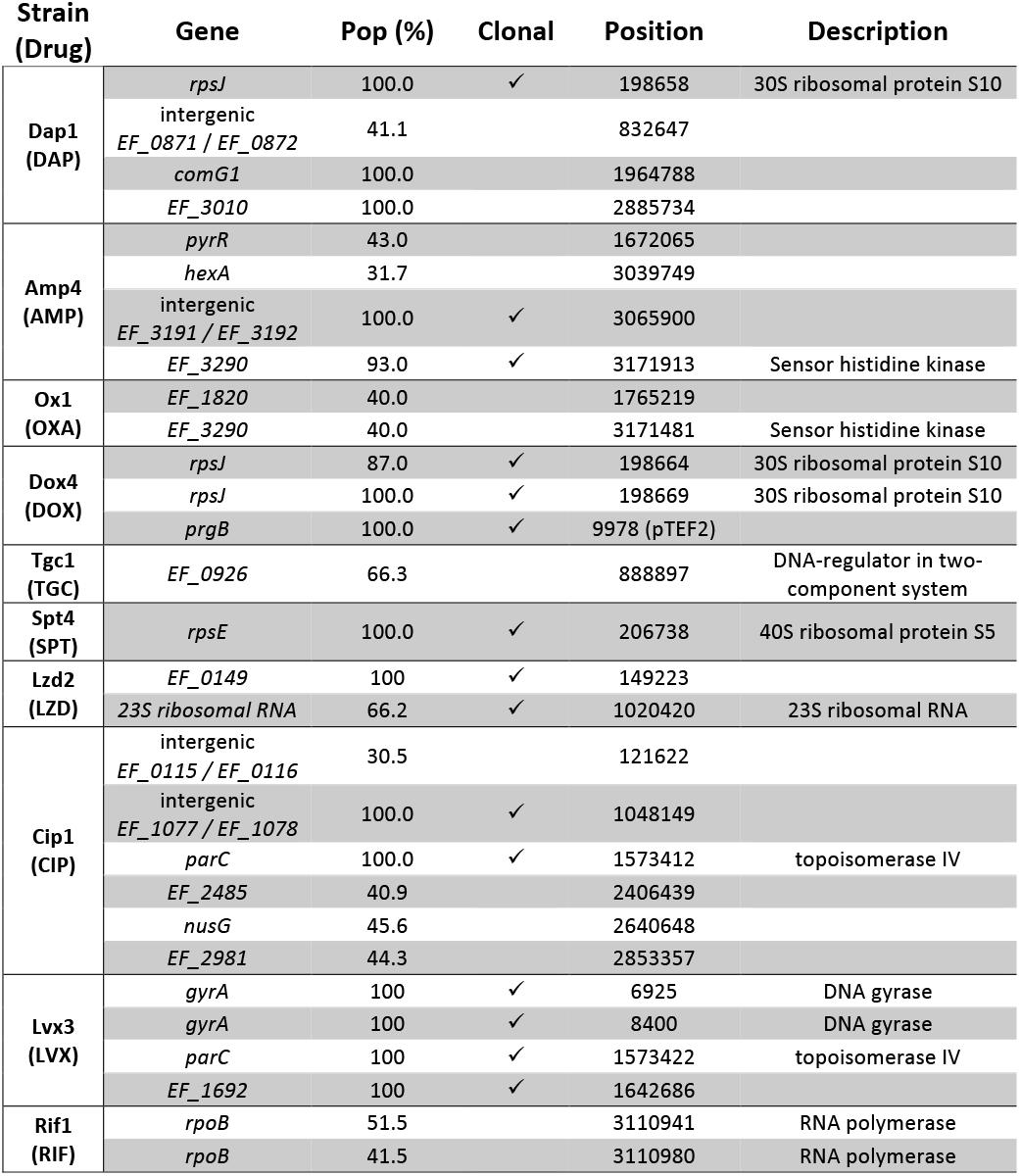
Mutations identified in selected populations. Check marks indicate the same mutation was also identified in clonal isolates from the same population.

**TABLE 2.**
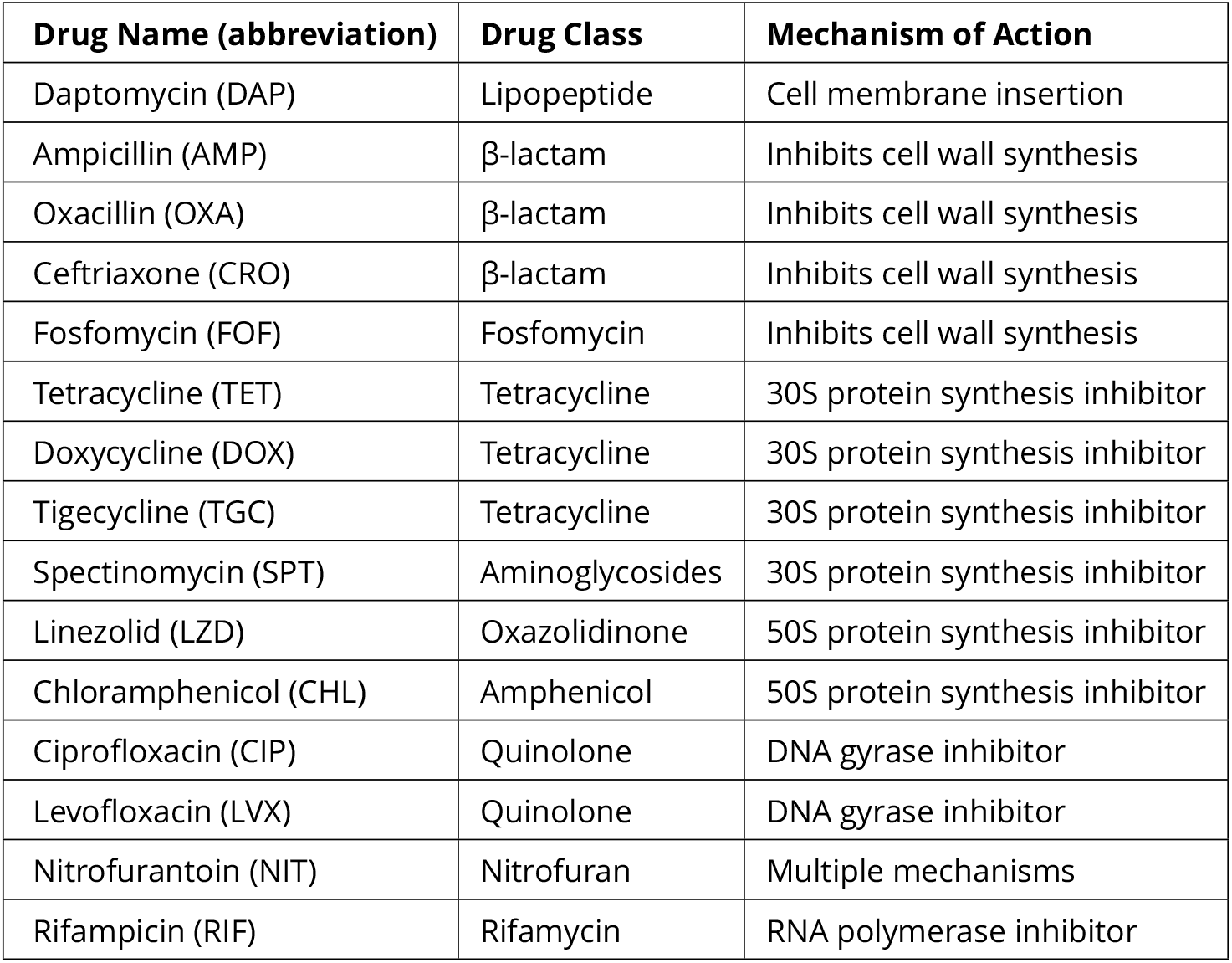
Table of antibiotics used in this study and their targets.

| Drug Name (abbreviation) | Drug Class | Mechanism of Action |
| --- | --- | --- |
| Daptomycin (DAP) | Lipopeptide | Cell membrane insertion |
| Ampicillin (AMP) | β-lactam | Inhibits cell wall synthesis |
| Oxacillin (OXA) | β-lactam | Inhibits cell wall synthesis |
| Ceftriaxone (CRO) | β-lactam | Inhibits cell wall synthesis |
| Fosfomycin (FOF) | Fosfomycin | Inhibits cell wall synthesis |
| Tetracycline (TET) | Tetracycline | 30S protein synthesis inhibitor |
| Doxycycline (DOX) | Tetracycline | 30S protein synthesis inhibitor |
| Tigecycline (TGC) | Tetracycline | 30S protein synthesis inhibitor |
| Spectinomycin (SPT) | Aminoglycosides | 30S protein synthesis inhibitor |
| Linezolid (LZD) | Oxazolidinone | 50S protein synthesis inhibitor |
| Chloramphenicol (CHL) | Amphenicol | 50S protein synthesis inhibitor |
| Ciprofloxacin (CIP) | Quinolone | DNA gyrase inhibitor |
| Levofloxacin (LVX) | Quinolone | DNA gyrase inhibitor |
| Nitrofurantoin (NIT) | Nitrofuran | Multiple mechanisms |
| Rifampicin (RIF) | Rifamycin | RNA polymerase inhibitor |

Our results indicate that collateral effects-including sensitivity-are pervasive, with approximately 73 percent (612/840) of all (collateral) drug-mutant combinations exhibiting a statistically significant change in IC_50_. By contrast, none of the four V583 strains propagated in BHI alone showed any collateral effects. The isolates exhibit collateral sensitivity to a median of 4 drugs, with only 3 of the 60 mutants (5 percent) exhibiting no collateral sensitivity at all; on the other hand, mutants selected by ceftriaxone (CRO) and fosfomycin (FOF) exhibit particularly widespread collateral sensitivity. Cross resistance is similarly prevalent, with only 2 strains failing to exhibit cross resistance to at least one drug. Somewhat surprisingly, 56 of 60 mutants exhibit cross resistance to at least one drug from a different class (e.g. all mutants evolved to ciprofloxacin (CIP), a DNA synthesis inhibitor, show increased resistance CRO, an inhibitor of cell wall synthesis). The collateral effects can also be quite large; we measured 8 instances of collateral sensitivity where the IC_50_ decreases by 16 fold or more. We observe a strong, repeatable collateral sensitivity to rifampicin (RIF) when mutants were selected by inhibitors of cell wall synthesis, an effect that-to our knowledge-has not been reported elsewhere. More typically, however, collateral effects are smaller than the direct effects to the selecting drug, with 46 percent (384/840) exhibiting more than a factor 2 change in IC_50_ and only 7 percent (61/840) exhibiting more than a factor 4 change.

### Isolates exhibit variability in fitness costs and collateral prof les

To investigate the potential impact of resistance mutations on fitness, we estimated the specific growth rate and the lag time in drug-free media for isolates selected from each of the 60 populations (4 populations per selecting drug). The growth costs exhibit significant variability, even across different populations selected by the same drug (Figure 2, similar to results in other species (49). In some isolates-such as those selected by oxacillin (OXA) or nitrofurantoin (NIT)-growth rate and lag times are indistinguishable from those of the ancestral strains. On the other hand, isolates selected by CRO and FOF-selecting conditions that frequently result in collateral sensitivity-show dramatically reduced growth and an increased lag time, suggesting that the selected resistance determinants are associated with strong pleiotropic effects even in drug-free media.

**FIG 2.**
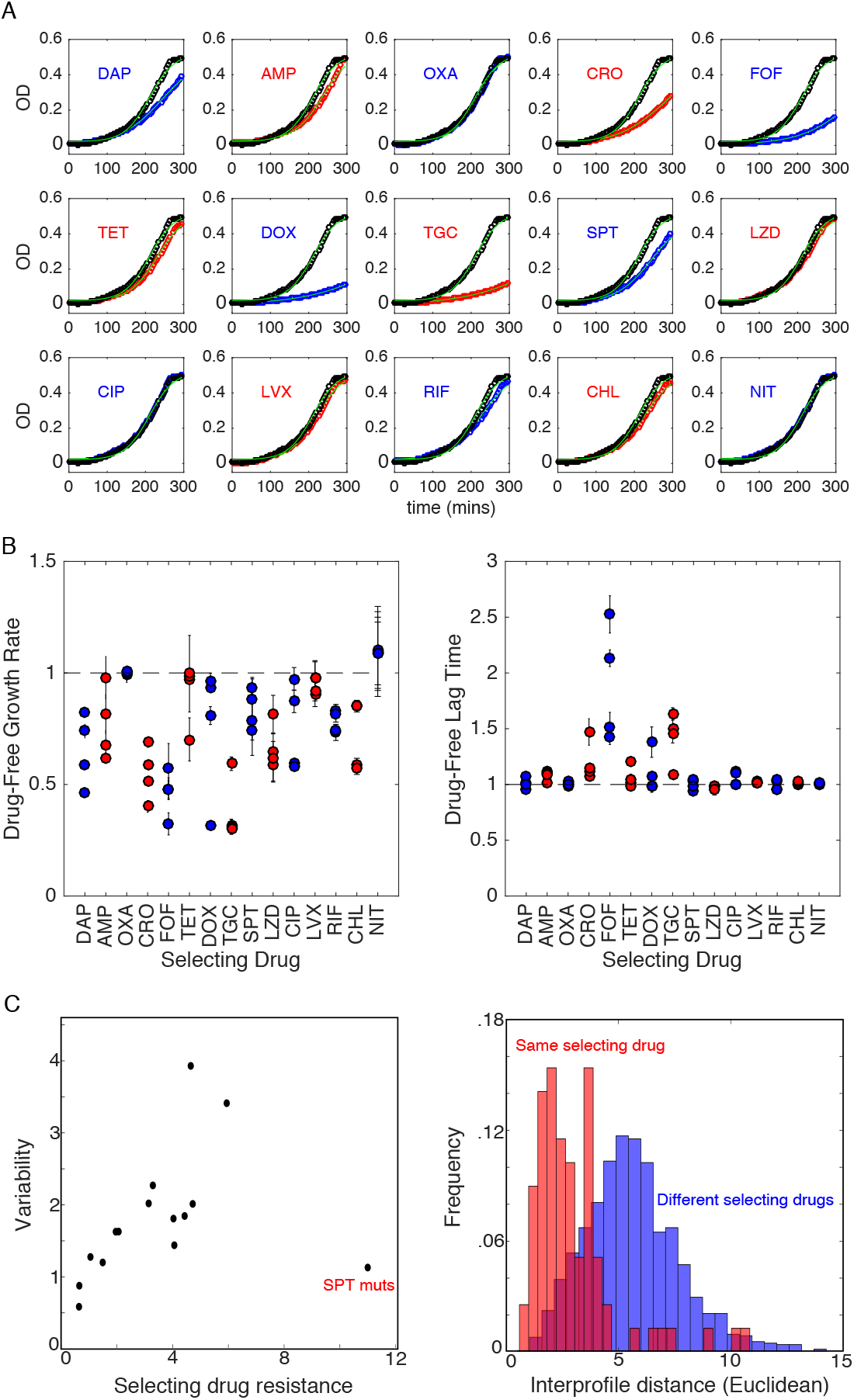
Growth costs and lag times for isolates selected by different antibiotics A. Example optical density (OD) time series for single isolates selected by each of the 15 drugs. Blue or red circles correspond to the isolate, black circles to ancestral strains. Light green lines show fits to logistic growth function (60) given by *g*(*t*) = *g*_0_ + *K* (1 + exp(4*μ*(*λ* − *t*)/*K* + 2))^-1^, where *μ* is the maximum specific growth rate, *λ* is the lag time, and *K* is the carryingcapacity. To reduce the number of free parameters, we fix *K* = 0.5 to match that of the ancestral strain. B. Maximum specifc growth rate (*μ*, left) and lag time (*λ*, right) in drug-free media for isolates from each of the four populations selected by each drug. All values are normalized by the values measured in the ancestral strain. Errorbars are standard errors of the mean estimated from 3 technical replicates for each isolate. C. Left panel: variability in replicates for all 15 drugs vs the (log2-scaled) fold increase in IC50 to the selecting drug. Variability is defined as 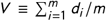, where *m* = 4 is the number of replicates and *d_i_* is the Euclidean distance between mutant *i* and the centroid formed by all vectors corresponding to a given selecting drug (Figure S2). Right panel: histogram of Euclidean distances between collateral profiles in pairs of isolates selected by the same (red) or different (blue) drugs. To emphasize collateral, rather than direct, effects, the component(s) of each collateral profile corresponding to the selecting drug(s)were removed prior to calculating variability (panel C) and pairwise Euclidean distances (D).

Our results indicate that collateral profiles can vary significantly even when mutants are evolved in parallel to the same drug (Figure 1C). For example, all 4 mutants selected by daptomycin (DAP) exhibit high-level resistance to the selecting drug, but replicates 1 and 4 exhibit collateral resistance to CRO, while replicate 2 exhibits collateral sensitivity and replicate 3 shows little effect (Figure 1C, right panel). To quantify the variation between replicates selected by the same drug, we considered the collateral profile of each mutant (i.e. a column of the collateral sensitivity matrix) as a vector in 15dimensional drug resistance space. Then, for each set of replicates, we defined the variability 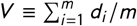, where *m* = 4 is the number of replicates and *d_i_* is the Euclidean distance between mutant *i* and the centroid formed by all vectors corresponding to a given selecting drug (Figure S2). Variability differs for different selecting drugs, with DAP and RIF showing the largest variability and NIT the smallest (Figure S2). We find that the variability is significantly correlated with average resistance to the selecting drug, even when one removes contributions to variability from the selecting drug itself (Figure 2C, left), indicating that collateral (rather than direct) effects underlie the correlation. Such a correlation might be expected if, for example, resistance arises from an accumulation of stochastic events following a Poisson-like distribution, where the mean is proportional to the variance. We do note, however, that selection by spectinomcyin (SPT) represents a notable exception to this trend. These results suggest that the repeatability of collateral effects is sensitive to the drug used for selection. As a result, certain drugs may be more appropriate for establishing robust antibiotic cycling profiles.

To further quantify the variability within and between isolates selected by different drug, we calculated the pairwise Euclidian distance between collateral profiles of isolates selected in the same drug and pairs of isolates selected in different drugs (Figure 2C, right). We see the distributions do have some overlap; that is, pairs of isolates selected by the same drug are sometimes more distinct from one another, by this metric, than pairs selected by different drugs. However, the distribution for different selecting drugs (blue) is shifted significantly to the right, indicating that isolates selected by the same drug are more similar to one another (on average) than to isolates selected by different drugs.

### Cross resistance to daptomycin appears frequently under selection by different drugs

Daptomycin is a lipopeptide antibiotic sometimes used as a last line of defense against gram-positive bacterial infections, including vancomycin resistant enterococci (VRE). While DAP resistance was initially believed to be rare (61), it has become increasingly documented in clinical settings (62). Recent work in a related enterococcal species has shown that cross resistance to DAP can arise from serial exposure to chlorhexidine, a common antiseptic (63), but less is known about DAP cross resistance following exposure to other antimicrobial agents. Surprisingly, our results indicate that DAP resistance is common when populations are selected by other antibiotics, with 64 percent of all evolved lineages displaying DAP cross resistance and only 11 percent displaying collateral sensitivity (Figure 3A).

**FIG 3.**
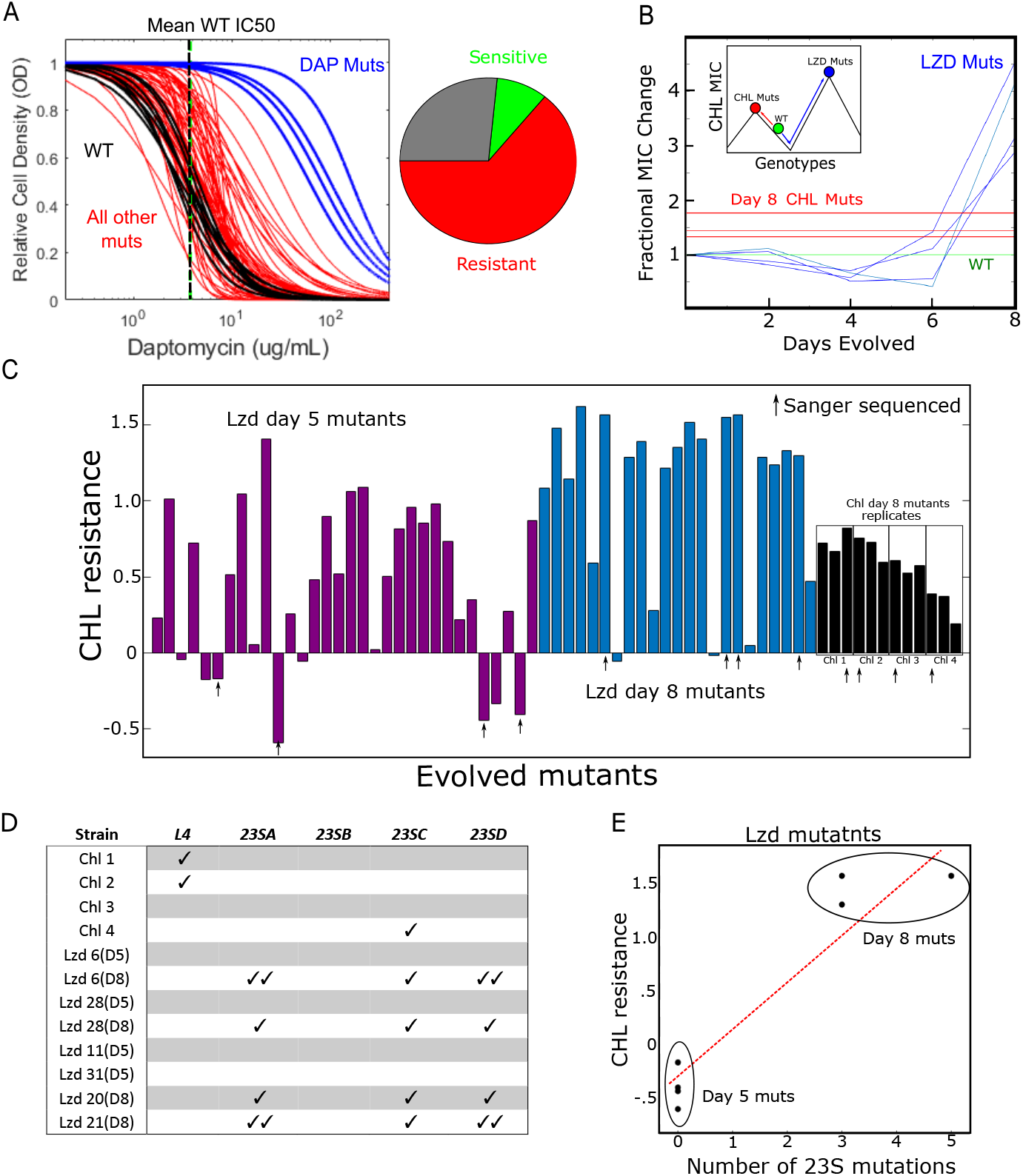
Collateral effects can lead to frequent or high-level resistance to non-selecting drugs. A. Estimated dose response curves (fit to Hill-like function) for all mutants tested against daptomycin. Strains evolved to daptomycin (blue) and all other drugs (red) frequently exhibit increased resistance to daptomycin relative to wild-type (black, individual replicates; dotted black line, mean IC_50_). Right inset: Approximately 64 percent of all drug-evolved mutants exhibit increased daptomycin resistance, while only 11 percent exhibit collateral sensitivity. B. Fractional change in chloramphenicol (CHL) IC_50_ for mutants evolved to linezolid (blue). The width of the green line represents the confidence interval (± 3 standard errors of the mean measured over 8 replicate measurements) for the (normalized) choramphenicol IC _0_ in wild-type cells. For comparison, the red lines represent the final (day 8) CHL resistance achieved in populations evolved directly to CHL. Inset: Schematic depicting two hypothetical paths to different CHL resistance maximums. The green circle represents the sensitive wild-type. Evolution can occur to CHL directly (red line) or to CHL collaterally through LZD resistance (blue line). The LZD evolution depicts early collateral sensitivity before ultimately achieving a higher total resistance. C. CHL resistance (log_2_-scaled change in IC_50_ relative to ancestor) for LZD-selected isolates at day 5 (purple) and day 8 (blue), and for individual colony isolates (4) for each of the four CHL-selected populations (black). Arrows indicate 12 isolates chosen for Sanger sequencing. D. Mutations observed in four different genes associated with LZD-resistance in each of the 12 selected isolates from panel C. E. CHL resistance and number of 23S mutations in LZD isolates on days 5 and 8.

### Selection by LZD leads to higher CHL resistance than direct selection by CHL

Surprisingly, we found that isolates selected by linezolid (LZD) developed higher resistance to chloramphenicol (CHL) than isolates selected directly by CHL (Figure 3B). The isolates from LZD and from CHL exhibit similar growth and lag-time distributions in drug-free media (Figure 2), suggesting that this effect is not driven by fitness costs alone. To investigate further, we isolated LZD-selected mutants at days 2, 4, 6 and 8 of the laboratory evolution and measured the resistance of each to CHL. We find that early-stage (days 4-6) mutants exhibit low level CHL sensitivity just prior to a dramatic increase in cross resistance around day 8. These findings suggest LZD selection drives the population across a CHL fitness valley, ultimately leading to levels of resistance that exceed those observed by direct CHL selection (Figure 3B, inset).

To further investigate the repeatability of this phenomenon, we exposed 32 additional populations to increasing LZD concentrations in parallel over 8 days. Using the four initial LZD mutants as a guide, we measured the CHL susceptibility of isolates from each population at day 5 (Figure 3C, purple) and day 8 (Figure 3C, blue). In addition, to account for potential heterogeneity in the original populations, we measured CHL susceptibility in three different (single colony) isolates from each of the original four populations selected in CHL (Figure 3C, black). On day 5, almost a third (10 of 32) of the LZD-selected strains exhibited CHL resistance greater than that of any day 8 CHL-selected strains, while 25 percent (8 of 32) were more CHL-sensitive than even the ancestral strains. By contrast, on day 8 the vast majority of isolates (17 of 23) were highly CHL-resistant, with only a few strains (2 of 23) exhibiting small levels collateral sensitivity.

To identify genes that may be responsible for collateral CHL resistance, we PCR-amplified and (Sanger) sequenced seven genes (*rpsJ, L3, L4*, which are genes for ribosomal proteins, and four genes for 23S rRNA, *23SA, 23SB, 23SC, 23SD*) previously associated with LZD resistance (64) in a subset of 12 isolates. We selected the most CHL-resistant isolate from each CHL population, two pairs of day 5/day 8 LZD-selected isolates that exhibited collateral sensitivity on day 5 and cross resistance on day 8, two LZD-selected isolates with high-level collateral sensitivity to CHL on day 5, and two LZD isolates with large cross resistance on day 8 (Figure 3C; specific isolates marked by black arrows). We did not observe mutations in *rpsJ, L3* or *23SB* in any strain. In addition, the four LZD-selected isolates showed no mutations in any of the sequenced genes on day 5 (Figure 3D). By contrast, all 4 of the LZD-selected strains contained at least 3 mutations in the 23S rRNA genes on day 8. Two of the CHL-selected isolates had mutations in *L4* and one had a single mutation in the 23SC gene.

We observe a strong correlation between the level of CHL resistance and the total number of 23S rRNA mutations, similar to the dosing behavior previously observed for LZD (64). This correlation suggests that the 23S mutations found in LZD-selected (and CHL-resistant) isolates from day 8-but missing in the CHL-sensitive isolates from day 5-may be responsible for the later-stage, high-level cross resistance to CHL. Elucidating the precise evolutionary dynamics underlying differential selection for these mutations in LZD and CHL remains an open question, though the early (day 5) CHL-sensitivity observed in LZD-selected isolates suggests it may be necessary to cross a fitness valley in CHL resistance in order to eventually achieve higher CHL resistance.

### Whole-genome sequencing reveals known resistance determinants and mutations in genes previously linked with collateral sensitivity

To investigate the genetic changes in drug-selected populations, we sequenced population samples from one evolved population per drug. In addition, we isolated and sequenced a single clone from each population. As controls, we sequenced two different isolates from the ancestral V583 stock as well as both single isolates and a population sample propagated in drug-free media. We then used breseq (65), an established computational pipeline capable of mutant identification in both clonal and population samples (Methods). To minimize potential artifacts from sample preparation or analysis, we excluded from further analysis four populations where variants identified by clonal and population samples did not overlap. In addition, we limit our focus to those mutations estimated to occur with frequency greater than 30 percent in the population samples.

This analysis revealed a total of 29 mutations in the 11 populations (Table 1; note that the population selected in NIT contained no identifiable mutations). The control strain propagated in BHI contained no mutations relative to the ancestral strains. For 9 of the 11 selecting drugs, we identified mutations that likely confer resistance to the selecting drug. For example, we observed mutations in drug targets associated with protein synthesis inhibitors (*rpsJ* (66), *rpsE* (*67*)), fluoroquinolones (*parC, gyrA* (68)), and RNA synthesis inhibitors (*rpoB* (68)). We also identified mutations in a sensor histidine kinases (69, 70) (EF_3290) in populations selected by cell-wall inhibitors and mutations in 23S rRNA genes in the LZD-selected population (64, 71). Surprisingly, the DAP-selected population did not contain mutations in genes previously identified with DAP resistance (57, 58), though we observe a mutation in *rpsJ* in both the clonal and population sequences. While previous experiments have shown rpsJ not to confer DAP resistance in one genetic background of *E. faecalis* strain S613 (66), it may underlie the observed cross resistance to other antibiotics. Finally, we observe no mutations in either the clonal or population sequencing for the Nit1 population, despite repeated experiments confirming increased resistance to NIT. Because the resistance is relatively low-level (IC_50_ increases by approximately 50 percent relative to ancestor), it is possible the observed resistance corresponds to transient phenotypic resistance, similar to the post-antibiotic effect or the cellular hysteresis observed when drugs are rapidly cycled (46). Finally, the TGC-selected population contains a mutation in EF_0926, the response regulator in a two-component signaling system (TCS) with the sensor kinase EF_0927. While this specific system has not been implicated in tigecycline (TGC) resistance, similar TCS have been linked to TGC resistance in *A. baumannii* (72).

Several of the mutations we identified occur in genes previously linked with collateral effects in other species. For example, mutations in the topisomerase gene *gyrA* have been posited to induce collateral sensitivities via global transcriptional changes induced by modulated DNA supercoiling (35, 73, 74). Similarly, mutations in ribosomal genes, such as *rpsE*, have been linked with multi-drug resistance modulated by large-scale changes in the transcriptome (75).

### Sensitivity profiles cluster into groups based on known classes of selecting drug

Our results indicate that there is significant heterogeneity in collateral sensitivity profiles, even when parallel populations are selected with the same antibiotic. While the genetic networks underlying these phenotypic responses are complex and, in many cases, poorly understood, one might expect that selection by chemically or mechanistically similar drugs would lead to profiles with shared statistical properties. For example, previous work showed (in a different context) that pairwise interactions between simultaneously applied antibiotics can be used to cluster drugs into groups that interact monochromatically with one another; these groups correspond to known drug classes (76), highlighting statistical structure in drug interaction networks that appear, on the surface, to be extremely heterogeneous. Recent work in bacteria has also shown that phenotypic profiles of mutants selected by drugs from the same class tend to cluster together in *P. aeruginosa* (38) and *E. coli* (77).

Similarly, we asked whether collateral sensitivity profiles in *E.faeaclis* can be used to cluster resistant mutants into statistically similar classes. We first performed hierarchical clustering (Methods) on collateral profiles of 52 different mutants (Figure 4, x-axis; note that we excluded mutants selected by CHL and NIT, which did not achieve resistance of at least 2x to the selecting drug). Despite the heterogeneity in collateral profiles, they cluster into groups characterized-exclusively-by selecting drugs from the same drug classes before grouping mutants from any two different drug classes. For example, inhibitors of cell wall synthesis (AMP, CRO, FOF, OXA) cluster into one group (noted by A in Figure 3), while tetracycline-like drugs (TET, DOX, TGC) cluster into another (noted by B). This approach also separates spectinomycin (SPT, aminoglycoside antibiotic class) from the tetracycline class of antibiotics (TET, DOX, TGC) despite the fact that they both target the 30S subunit of the ribosome, suggesting that it may help identify drugs with similar mechanisms but statistically distinct collateral profiles.

**FIG 4.**
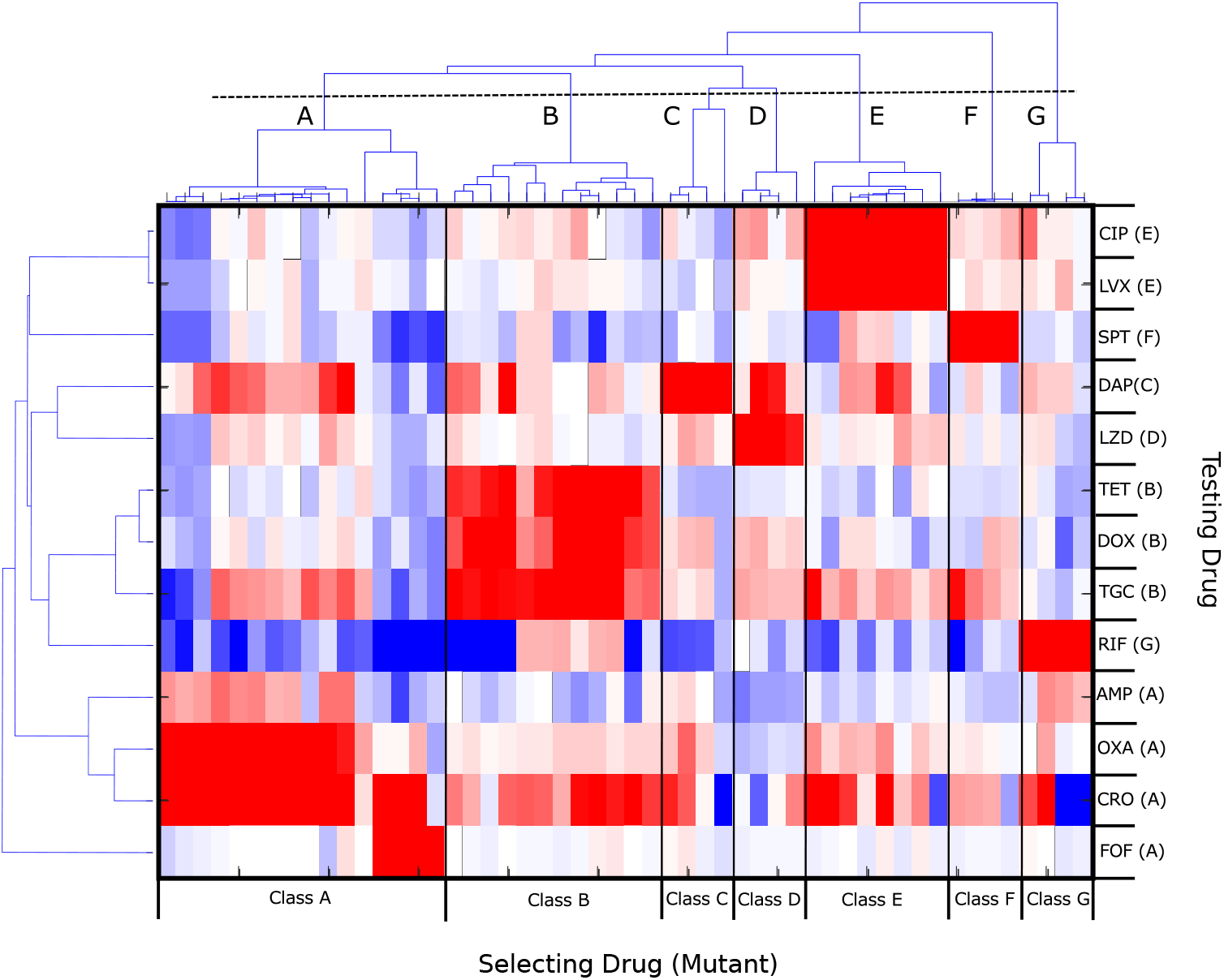
Hierarchical clustering of collateral sensitivity profiles partitions mutants into groups selected by known drug classes. Heatmap with ordering of rows (testing drug) and columns (4 replicate experiments with the same selecting drug) determined via hierarchical clustering. Colormap and scale are identical to those used in Figure 1. Collateral profiles (columns) for mutants selected by drugs from known drug classes (here labeled A-G) cluster together; if clusters are defined using the dashed line (top), there are 7 distinct clusters, each corresponding to a particular drug class: A. Cell-wall synthesis inhibitors (AMP, OXA, CRO, FOF), B. Tetracyclines (TET, DOX, TGC), C. Lipopeptides (DAP), D. Oxazolidinones (LZD), E. Fluoroquinolones (CIP, LVX), F. Aminocyclitols (SPT), and G. Antimycobacterials (RIF). When clustering the testing drugs (rows), drugs from the same class are frequently but not always clustered together. For example, cell-wall drugs such as AMP, OXA, and CRO form a distinct cluster that does not include FOF (bottom 4 rows).

We then performed a similar clustering analysis of the collateral responses across the 14 different testing drugs (Figure 4, y-axis), which again leads to groupings the correspond to known drug classes. One drug, FOF, provides an interesting exception. Mutants selected for FOF resistance cluster with those of other cell-wall synthesis inhibitors (Class A, columns). However, the behavior of FOF as a testing drug (last row) is noticeably distinct from that of other cell-wall synthesis inhibitors (the 3 rows directly above FOF). Taken together, the clustering analysis reveals clear statistical patterns that connect known mechanisms of antibiotics to their behavior as both selecting and testing agents.

### A Markov decision process (MDP) model predicts optimal drug policies to constrain resistance

Our results indicate that collateral sensitivity is pervasive, and while collateral sensitivity profiles are highly heterogeneous, clustering suggests the existence of statistical structure in the data. Nevertheless, because of the stochastic nature of the sensitivity profiles, it is not clear whether this information can be leveraged to design drug sequences that constrain evolution. It is important to note that our goal, at this stage, is not to design specific drug sequences that might be transferred directly to the clinic, but instead to evaluate-in a simple setting-the feasibility of slowing resistance in even the most optimized cases. Given that resistance to the selecting drugs is often larger in magnitude than collateral (off-diagonal) effects, it is not clear a priori that a feasible strategy exists that prevents the inevitable march to high-level resistance, even in a highly idealized setting.

To address this problem, we develop a simple mathematical model based on a Markov decision process (MDP) to predict optimal drug policies. MDP’s are widely used in applied mathematics and finance and have a well-developed theoretical basis (78, 79, 80). In a MDP, a system transitions stochastically between discrete states. At each time step, we must make a decision (called an “action”), and for each state-action combination there is an associated instantaneous “reward” (or cost). The action influences not only the instantaneous reward, but also which state will occur next. The goal of the MDP is to develop a policy-a set of actions corresponding to each state-that will optimize some objective function (e.g. maximize some cumulative reward) over a given time period.

For our system, the state *s_t_* at time step *t* = 0,1,2,… is defined by the resistance profile of the population, a vector that describes the resistance level to each available drug. At each time step, an action *a_t_* is chosen that determines the drug to be applied. The system-which is assumed to be Markovian-then transitions with probability *P_a_*(*s*_*t*+1_ |*s_t_, a_t_*) to a new state *s*_*t_i_*+1_, and the transition probabilities are estimated from evolutionary experiments (or any available data). The instantaneous reward function *R_a_* (*s*) is chosen to be the (negative of the) resistance to the currently applied drug; intuitively, it provides a measure of how well the current drug inhibits the current population. The optimal policy *π*\*(*s*) is formally a mapping from each state to the optimal action; intuitively, it tells which drug should be applied for a given resistance profile. The policy is chosen to maximize a cumulative reward function 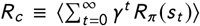, where brackets indicate an expectation value conditioned on the initial state *s*_0_ and the choice of policy *π*. The parameter *γ* (0 ≤ *γ* < 1) is a discount factor that determines the timescale for the optimization; *γ* ≈ 1 leads to a solution that performs optimally on long timescales, while *yγ* ≈ 0 leads to solutions that maximize near-term success.

To apply the MDP framework to collateral sensitivity profiles, we must infer from our data a set of (stochastic) rules for transitioning between states (i.e. we must estimate *P_a_*(*s*_*t*+1_ |*s_t_, a_t_*)). While many choices are possible-and different rules may be useful to describing different evolutionary scenarios-here we consider a simple model where the resistance to each drug is increased/decreased additively according to the collateral effects measured for the selecting drug in question. Specifically, the state *s*_*t*+1_ following application of a drug at time *t* is given by 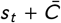, where 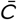 is one of the four collateral profiles (see Figure 1) measured following selection by that drug. Because resistance/sensitivity is measured using log-scaled ratios of IC_50_’s, these additive changes in the resistance profile correspond to multiplicative changes in the relative IC_50_ for each drug. For instance, if one selection step increases the IC_50_ by a factor of 3, then two consecutive selection steps would increase IC_50_ by a factor of 9. This model assumes that selection by a given drug always produces changes in the resistance profile with the same statistical properties. For example, selection by DAP increases the resistance to DAP (with probability 1) while simultaneously either increasing resistance to AMP (with probability 1/4), decreasing resistance to AMP (with probability 1/4), or leaving resistance to AMP unchanged (probability 1/2). Repeated application of the same drug will steadily increase the population’s resistance to that drug, but the process could potentially sensitize the population to other drugs. This model implicitly assumes sufficiently strong selection that, at each step, the state of the system is fully described by a single “effective” resistance profile (rather than, for example, an ensemble of profiles that would be required to model clonal interference). While we focus here on this particular model, we stress that this MDP framework can be easily adopted to other scenarios by modifying *P_a_*(*s*_*t*+1_ |*s_t_, a_t_*).

For numerical efficiency, we discretized both the state space (i.e. the resistance to each drug is restricted to a finite number of levels) as well as the measured collateral profiles (exposure to a drug leads to an increase/decrease of 0,1, or 2 resistance levels; Figure 5A, Figure S3). In practice, this means that resistance will eventually saturate at a finite value if a single drug is applied repeatedly. In addition, we restrict our calculations to a representative subset of six drugs (DAP, AMP, FOF, TGC, LZD, RIF). The set includes inhibitors of cell wall, protein, or RNA synthesis, and five of the six drugs (excluding RIF) are clinically relevant for enterococcus infections. We note, however, that the results are qualitatively similar for different discretization schemes (Figure S4) and for different drug choices (Figures S5–S7).

**FIG 5.**
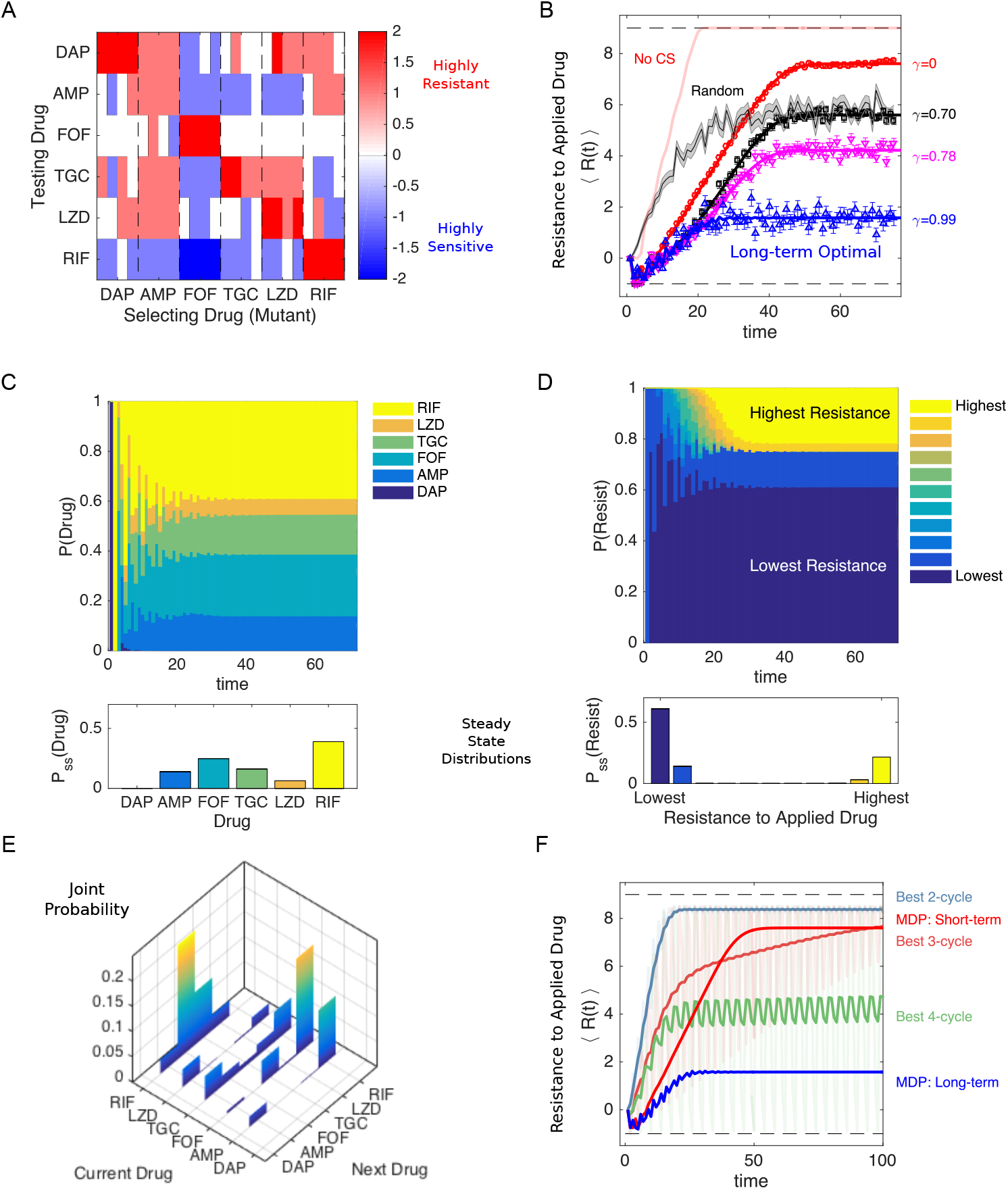
Simulated optimal drug sequences constrain resistance on long timescales and outperform simple collateral sensitivity cycles A. Discretized collateral sensitivity or resistance *C_d_* ∈ {-2,-1,0,1,2} for a selection of six drugs. For each selecting drug, the heat map shows the level of cross resistance or sensitivity (*C_d_*) to each testing drug (the subscript d indicates the profiles are discretized) for *n_r_* = 4 independently evolved populations. See Figure 1 for original (non-discretized) data. B. Average level of resistance (〈*R*(*t*)〉) to the applied drug for policies with *γ* = 0 (red), *γ* = 0.7 (black), *γ* = 0.78 (magenta), and *γ* = 0.99 (blue). Resistance to each drug is characterized by 11 discrete levels arbitrary labeled with integer values from −1 (least resistant) to 9 (most resistant). At time 0, the population starts in the second lowest resistance level (0) for all drugs. Symbols (circles, triangles, squares) are the mean of 10^3^ independent simulations of the MDP, with error bars ± SEM. Solid lines are numerical calculations using exact Markov chain calculations (see Methods). Light red line, long-term optimal policy (*γ* = 0.99) calculated using the data in A but with collateral sensitivity values set to 0. Black shaded line, randomly cycled drugs (± SEM). C. The time-dependent probability P(Drug) of choosing each of the six drugs when the optimal policy (*γ* = 0.99) is used. Inset, steady state distribution P_*s*_*s*(Drug). D. The probability P(Resist) of the population exhibiting a particular level of resistance to the applied drug when the optimal policy (*γ* = 0.99) is used. Inset, steady state distribution P_*s*_*s*(Drug). E. Steady state joint probability distribution P(current drug, next drug) for consecutive time steps when the optimal policy (*γ* = 0.99) is used. F. Average level of resistance (〈*R*(*t*)〉) to the applied drug for collateral sensitivity cycles of 2 (dark green, LZD-RIF), 3 (pink, AMP-RIF-LZD), or 4 (dark green, AMP-RIF-TGC-LZD) drugs are compared with MDP policies with *γ* = 0 (short-term, red) and *γ* = 0.99 (long-term, blue). For visualizing the results of the collateral sensitivity cycles, which give rise to periodic behavior with large amplitude, the curves show a moving time average (window size 10 steps), but the original curves are shown transparently in the background.

### Drug policies can be tuned to minimize resistance on different timescales

The optimal policy *π*\*(*s*) is a high-dimensional mapping that is difficult to directly visualize. For intuition on the policy, we calculated the frequency with which each drug is prescribed as a function of resistance to each of the six individual drugs (Figures S8, S9; top panels). Not surprisingly, we found that when resistance to a particular drug is very low, that drug is often chosen as optimal. In addition, the specific frequency distributions vary significantly depending on *γ*, which sets the timescale of the optimization. For example, the long-term optimal policy (*γ* = 0.99) yields a frequency distribution that is approximately independent of the level of resistance to FOF (Figure S8, upper right panel). By contrast, the frequency distribution for a short-term policy (*γ* = 0.1) changes with FOF resistance; at low levels of resistance, FOF is frequently applied as the optimal drug, but it is essentially never applied once FOF resistance reaches a certain threshold (Figure S9, upper right panel). Both the short- and long-term optimal policies lead to aperiodic drug sequences, but the resulting resistance levels vary significantly (Figures S8, S9, bottom panels). These differences reflect a key distinction in the policies: short-term policies depend sensitively on the current resistance level and maximize efficacy (minimize resistance) at early times, while long-term policies may tolerate short-term performance failure in exchange for success on longer timescales.

### Optimal policies outperform random cycling and rely on collateral sensitivity

To compare the outcomes of different policies, we simulated the MDP and calculated the expected resistance level to the applied drug over time, ⌨*R*(*t*)〉), from 1000 independent realizations (Figure 5B). All MDP policies perform significantly better than random drug cycling for the first 10-20 time steps and even lead to an initial decrease in resistance. The long-term policy (*γ* = 0.99, blue) is able to maintain low-level resistance indefinitely, while the short-term policy (*γ* = 0) eventually gives rise to high-level (almost saturating) resistance. Notably, if we repeat this calculation on an identical data set but with all collateral sensitivities set to 0, the level of resistance rapidly increases to its saturating value (Figure 5B, light red line), indicating that collateral sensitivity is critical to the success of these policies. We note that the timescales used here are not necessarily reflective of a clinical situation, and instead our goal is to understand the performance of the optimization over a wide range of timescales.

### Optimal policies highlight new strategy for minimizing resistance

To understand the optimal policy dynamics, we calculated the time-dependent probability distributions P(Drug)-the probability of applying a particular drug-and P(Resist)-the probability of observing a given level of resistance to the applied drug-for the MDP following the long-term policy (*γ* = 0.99, Figure 5C-D). We also calculated the (steady state) joint probability distribution characterizing the prescribed drugs at consecutive time steps (Figure 5E). The distributions reveal highly non-uniform behavior; after an initial transient period, RIF is applied most often, followed by FOF, while DAP is essentially never prescribed. Certain patterns also emerge between consecutively applied drugs; for example, FOF is frequently followed by RIF. Somewhat surprisingly, the distribution of resistance levels is highly bimodal, with the lowest possible resistance level occurring most often, followed by the highest possible level, then the second lowest level, and then the second highest level (Figure 5D). The policy achieves a low average level of resistance not by consistently maintaining some intermediate level of resistance to the applied drug, but instead by switching between highly-effective drugs and highly-ineffective drugs, with the latter occurring much less frequently. In words, rare periods of high resistance are the price of frequent periods of very low resistance. These qualitative trends occur for other drug choices (Figures S5–S7) and are relatively insensitive to the number of discretization levels chosen (Figure S4). The results suggest a new conceptual strategy for minimizing resistance: interspersing frequent steps of instantly effective drugs (low resistance)-which provide short-term inhibition of pathogen growth-with rare steps of relatively ineffective drugs (high resistance), which provide little short-term inhibition but shepherd the population to a more vulnerable future state.

### Optimal policies maintain lower long-term resistance than collateral sensitivity cycles

The resurgent interest in collateral sensitivity was sparked, in part, by innovative recent work that demonstrated the successful application of collateral sensitivity cycles, where each drug in a sequence promotes evolved sensitivity to the next drug (41). To compare the performance of the MDP to that expected from collateral sensitivity cycles, we identified all collateral sensitivity cycles for the six drug network and calculated 〈*R*(*t*)〉 for 100 time steps of each cycle. We then determined the “best” cycle of a given length-defined as the cycle with the lowest mean value of 〈*R*(*t*)〉 over the last ten time steps-and compared the performance of those cycles to the short- and long-term MDP policies (Figure 5F). The MDP long-term optimal solution (*γ* = 0.99) maintains resistance at a lower average value than for all of the collateral sensitivity cycles. For MDP policies with shorter time horizons (e.g. the instant gratification cycle, *γ* = 0), however, the collateral sensitivity cycles of 3 and 4 drugs (as well as the long-term MDP solution) lead to lower resistance at intermediate or longer time scales, reflecting the inherent trade-offs between instantaneous drug efficacy and long-term sustainability. One advantage of the MDP optimization is that it allows for explicit tuning of the policy (via γ) to achieve maximal efficacy over the desired time horizon.

### Optimized drug sequences improve growth inhibition and reduce adaptation rates in lab evolution experiments

The MDP-based optimal policies perform well in stochastic simulations and highlight new strategies for potentially slowing resistance. However, the model contains a number of assumptions that lead to an oversimplified picture of the true evolutionary dynamics. As a result, it is not clear whether optimized drug sequences from this model will be effective in real, evolving pathogen populations.

To test the performance of MDP-based drug cycles, we designed a lab evolution experiment comparing inhibitory effects of different drug cycling protocols over 20 days. For experimental feasibility, we restrict our focus to a subset of four drugs (FOF, RIF, AMP, TGC) and reduced the length of each evolutionary time step from 8 days-as in the original collateral sensitivity experiment (Figure 1)- to 2 days. First, we experimentally measured the collateral sensitivity matrix for the four drug set following 2 days of lab evolution in eight replicate populations per drug (Figure 6A). We then calculated the optimal policy for two different values of *γ* (*γ* = 0.9, *γ* = 0.78), both corresponding to timescales commensurate with the planned experiment. In both cases, the steady state distribution of drug application P(Drug) calls for frequent use of TGC and relatively rare use of FOF, though the specific distribution depends on the particular choice of *γ* (Figure 6B, top panel; see also Figure S10).

**FIG 6.**
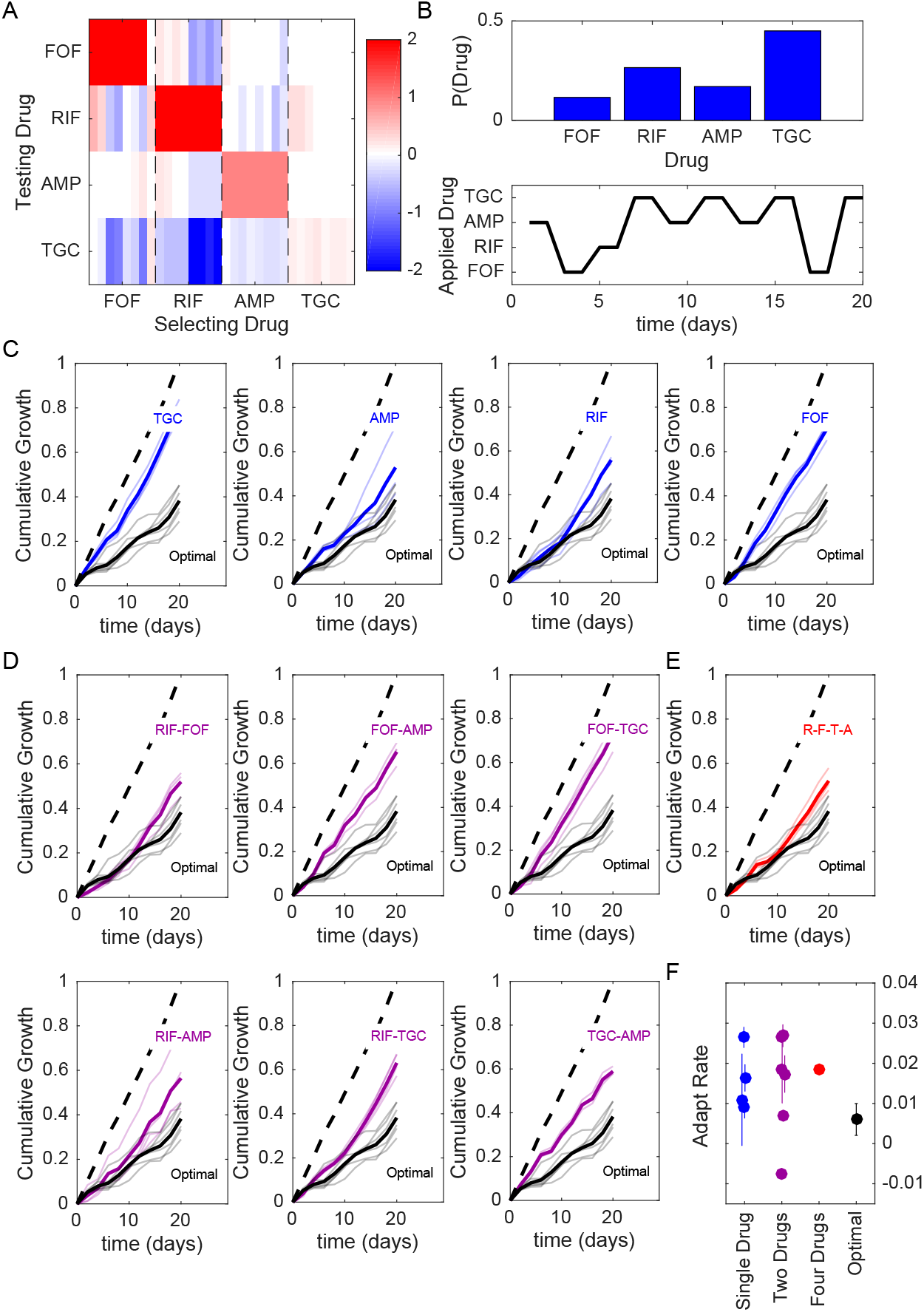
Optimized drug sequences reduce cumulative growth and adaptation rates in lab evolution experiments. A. Resistance (red) or sensitivity (blue) of each evolved mutant (horizontal axis; 4 drugs x 8 mutant per drug) to each drug (vertical axis) following 2 days of selection is quantified by the log_2_-transformed relative increase in the IC_50_ of the testing drug relative to that of wild-type (V583) cells. B. Top: distribution of applied drug at time step 20 (approximate steady state) calculated using an optimal policy with *γ* = 0.9. Bottom: sequence of applied drug from one particular realization of the stochastic process with the optimal policy (*γ* = 0.9). C-E. Cumulative population growth over time for populations exposed to single drug sequences (C, blue), two-drug sequences (D,magenta), a four drug sequence (E, red), or the optimal sequence from panel B (black curves, all panels). Transparent lines represent individual replicate experiments and each thicker dark line corresponds to a mean over replicates. Dashed line, drug-free control (normalized to a growth of 1 at the end of the experiment). F. Adaptation rate for single drug (blue), two-drug (magenta), four drug (red), and optimal sequences (black). Error bars are standard errors across replicates. Adaptation rate is defined as the slope of the best fit linear regression describing time series of daily growth (see Figure S14).

An exact application of the optimal policy requires measuring the full sensitivity profile at each step and using that profile, in accordance with the policy, to choose the next drug in the sequence. However, simulations suggest that choosing a predetermined drug cycle-that is, a cycle drawn from a particular realization of the stochastic process-is expected to perform near-optimally on the timescale of the experiment (Figures S11, S12). For experimental convenience, we choose a single MDP-derived cycle for each value of *γ*. For example, for *γ* = 0.9 the sequence involves ten 2-day time steps, with drugs applied in the following order: AMP-FOF-RIF-TGC-AMP-TGC-AMP-TGC-FOF-TGC (Figure 6B, bottom; see Figure S10 for *γ* = 0.78 example). To evaluate the efficacy of the MDP-derived cycle, we exposed a total of 60 replicate populations to one of 13 different drug cycle protocols (including the two MDP-derived cycles) over a 20-day serial passage lab evolution experiment. Every 40 hours, we measured the optical density of each population and then diluted each into fresh media containing the prescribed drug (added after a brief drug-free outgrowth phase, see Methods). Drug concentrations were chosen to be just above the MIC for ancestral populations-with MIC determined by complete absence of growth in ancestral strains after 24 hours under identical conditions-and the same concentration was applied at every time step calling for the associated drug. As a measure of drug efficacy, we defined the cumulative growth of a population at time t as the sum of the optical density measurements up to and including time *t*. Note that because drug-resistant populations often reach a steady-state carrying capacity-in our case, about OD=0.6-considerably faster than the 40-hour time window, cumulative growth is a conservative measure that underestimates differences in population size that would occur in exponentially-growing populations (for example, in a chemostat).

In addition to the two MDP-derived drug protocols, we also tested protocols calling for repeated application of each drug alone (Figure 6C), each of the six possible 2-drug cycles (Figure 6D), a four drug cycle consisting of repeated application of RIF-FOF-TGC-AMP ((Figure 6E), and a drug-free control (dashed lines, Figure 6C-E). In all cases, cumulative growth was normalized to the value of the drug-free control at the end of the 20-day experiment. To compare results from the model with experiment, we mapped each of the discrete resistance levels to an OD value, with the highest level of resistance corresponding to drug-free growth (OD≈ 0.6 each day) and the lowest resistance level corresponding to no growth (OD=0); see Figures S11, S12. We found experimentally that cycles involving sequential application of drugs with (on average) mutual collateral sensitivity-for example, cycles of RIF-FOF or RIF-AMP (see (Figure 6A)-are among the best-performing two-drug cycles, as predicted by previous studies (41). However, the MDP-derived protocols led to a significant reduction in cumulative growth, outperforming every other protocol, often by significant margins (Figure 6; Figure S10). In addition to cumulative growth, we characterized each trajectory by calculating the adaptation rate, which is defined as the average rate of increase of instantaneous growth over time (i.e. the slope of the best-fit regression line for instantaneous growth vs time over days 2-20, Figures S13–S14). Adaptation rate, which is essentially an estimate of the average convexity of the cumulative growth curves, provides no information on the magnitude of the growth at each step, but instead measures how rapidly that growth is increasing over time (starting with the first measurement after day 2). In addition to reducing cumulative growth, the MDP-derived sequences led to lower rates of adaptation than nearly every other protocol (Figure 6F; Figure S10F). A notable exception is the TGC-AMP cycle, which exhibits a (small) negative adaptation rate, reflecting that fact that growth at day 2 has already achieved relatively high levels-roughly 60 percent of drug-free growth-suggesting that adaptation largely occurs in that first period but is nearly absent after that.

## DISCUSSION

Our work provides an extensive quantitative study of phenotypic and genetic collateral drug effects in *E.faecalis.* We have shown that cross resistance and collateral sensitivity are widespread but heterogeneous, with patterns of collateral effects often varying even between mutants evolved to the same drug. Our results contain a number of surprising, drug-specific observations; for example, we observed a strong, repeatable collateral sensitivity to RIF when mutants were selected by inhibitors of cell wall synthesis. Additionally, cross-resistance to DAP is particularly common when cells are selected by other frequently used antibiotics. Because the FDA/CLSI breakpoint for DAP resistance is not dramatically different than the MIC distributions found in clinical isolates prior to DAP use (81), one may speculate that even small collateral effects could have potentially harmful consequences for clinical treatments involving DAP. In addition, we found that selection by one drug, LZD, led to higher overall resistance to CHL than direct selection by CHL. While choramphenicol is rarely used clinically, the result illustrates that 1) collateral effects can be highly dynamic, and 2) indirect selection may drive a population across a fitness valley to an otherwise inaccessible fitness peak.

Our findings also point to global trends in collateral sensitivity profiles. For example, we found that the repeatability of collateral effects is sensitive to the drug used for selection, meaning that some drugs may be better than others for establishing robust antibiotic cycling profiles. On the other hand, despite the apparent unpredictability of collateral effects at the level of individual mutants, the sensitivity profiles for mutants selected by drugs from known classes tend to cluster into statistically similar groups. As proof-of-principle, we show how these profiles can be incorporated into a simple mathematical framework that optimizes drug protocols while accounting for effects of both stochasticity and different time horizons. Within this framework, drug policies can be tuned to optimize either short-term or long-term evolutionary outcomes. The ability to systematically tune these timescales may eventually be useful in designing drug protocols that interpolate between short-term, patient-centric outcomes and long-term, hospital-level optimization.

Our results complement recent studies on collateral sensitivity and also raise a number of new questions for future work. Much of the previous work on collateral networks in bacteria has focused on gram-negative bacteria and highlighted the role of aminoglycosides in collateral sensitivity (41, 36). Many gram-positive bacteria, including enterococci, are intrinsically resistant to aminoglycosides (82), and we therefore included only one (SPT) in our study. In that case, however, we did observe collateral sensitivity to cell wall inhibitors (AMP and FOF) in SPT-selected populations, consistent with findings in other species (41, 36), though it is not clear from our results whether aminoglycoside resistance would be associated with more widespread collateral sensitivity in *E. faecalis.* Recent work demonstrates that collateral profiles may be largely conserved across a wide range of *E. coli* isolates (77), offering hope that large scale analysis of clinical isolates may soon identify similar patterns in enterococci.

Multiple studies have shown that collateral profiles are heterogeneous (49, 50), and optimization will therefore require incorporation of stochastic effects such as likelihood scores (51). These likelihood scores could potentially inform transition probabilities in our MDP approach, leading to specific predictions for optimal drug sequences based on known fitness landscapes. While we have quantified the variability in evolved populations in several ways (e.g. variability scores, interprofile distance, population sequencing), we cannot definitely comment on the source of that variability; it could arise, for example, from different fixation events in independent populations or, alternatively, from clonal interference and random sampling in isolating individual clones. Indeed, population sequencing does suggest some measure of heterogeneity, even when we limit our analysis to mutations occurring at greater than 30 percent. In any event, our results point to a rich collection of possible collateral profiles, meaning that successful approaches for limiting resistance will likely require incorporation of variability and heterogeneity.

Several previous studies have indicated that cycles involving mutually collaterally sensitive drugs may be chosen to minimize the evolution of resistance (41, 42). In the context of our MDP model, these cycles fall somewhere between the short-time-horizon optimization and the long-term optimal strategy, and in some cases, the collateral sensitivity cycling can lead to considerable slowing of resistance. However, our results indicate that the MDP optimizations on longer time-horizons lead to systematically lower resistance, a consequence of intermixing (locally) sub-optimal steps where the drug is instantaneously less effective but shepherds the population to a more vulnerable evolutionary state. We also find experimentally that mutual collateral sensitivity cycles with two drugs do generally outperform most other two-drug and single-drug protocols-as predicted by previous studies-but they generally underperform the MDP-based sequences.

It is important to keep in mind several limitations of our work. Designing effective drug protocols for clinical use is an extremely challenging and multi-scale problem. Our approach was not to develop a detailed, clinically accurate model, but instead to focus on a simpler question: optimizing drug cycles in single-species host-free populations. Even in this idealized scenario, which corresponds most closely to in vitro lab experiments, slowing resistance is a difficult and poorly understood problem (despite much recent progress). Our results are promising because they show systematic optimization is indeed possible given the measured collateral sensitivity profiles.

We have chosen to focus on a simple evolutionary scenario where collateral effects accumulate over time based on the history of drug exposure. By using a simple model that can be analyzed in detail, our goal was to identify new conceptual strategies-and with them, experimentally testable predictions-for exploiting correlations in phenotypic resistance profiles. While we’ve focused on an extremely simple model, the MDP framework can be readily extended to account for different evolutionary scenarios and to incorporate more complex clinically-inspired considerations. For example, it would be straightforward to include fitness costs associated with different resistance profiles; in turn, the model might be extended to allow for drug-free periods (“drug holidays”), which potentially exploit these fitness costs to minimize resistance (50). In addition, the current model inherently assumes that the dominant collateral effects are independent of the genetic background. In fact, collateral sensitivity profiles in cancer have been previously shown to be time-dependent (83, 50), epistasis certainly occurs (49, 84), and population heterogeneity could limit efficacy of this strategy under some conditions (85). Unfortunately, the frequency and relative impact of these confounding effects are difficult to gauge. However, the relative success of the MDP-inspired sequences in lab evolution experiments underscores the potential of the approach. In particular, our findings offer hope that strategies combining frequent use of highly effective drugs with rare periods of “evolutionary steering” by less effective drugs may be promising even when the detailed assumptions of the model do not strictly hold.

Our future work will focus on experimentally characterizing dynamic properties of collateral effects and expanding the MDP approach to account for time-varying sensitivity profiles and epistasis. It may also be interesting to investigate collateral effects in microbial biofilms, where antibiotics can have counterintuitive effects even on evolutionarily short timescales (86). On longer timescales, elegant experimental approaches to biofilm evolution have revealed that spatial structure can give rise to rich evolutionary dynamics (87, 88) and potentially, but not necessarily, divergent results for biofilm and planktonic populations (89)

Finally, our results raise questions about the potential molecular and genetic mechanisms underlying the observed collateral effects. The phenotypic clustering analysis presented here may point to shared mechanistic explanations for sensitivity profiles selected by similar drugs, and the full genome sequencing identifies candidate genes associated with increased resistance. However, fully elucidating the detailed genetic underpinnings of collateral sensitivity remains an ongoing challenge for future work. At the same time, because the MDP framework depends only on phenotypic measurements, it may allow for systematic optimization of drug cycling policies even when molecular mechanisms are not fully known.

## MATERIALS AND METHODS

### Strains, antibiotics and media

All resistant lineages were derived from *E. faecalis* V583, a fully sequenced vancomycin-resistant clinical isolate (90). The 15 antibiotics used are listed in Table 2. Each antibiotic was prepared from powder stock and stored at −20°C with the exception of ampicillin, which was stored at −80°C. Evolution and IC50 measurements were conducted in BHI medium alone with the exception of DAP, which requires an addition of 50 mM calcium for antimicrobial activity.

### Laboratory Evolution Experiments

Evolution experiments to each antibiotic were performed in quadruplicate. Evolutions were performed using 1 mL BHI medium in 96-well plates with maximum volume 2 mL. Each day, populations were grown in at least three different antibiotic concentrations spanning both sub- and super-MIC doses. After 16-20 hours of incubation at 37°C, the well with the highest drug concentration that contained visual growth was propagated into 2 higher concentrations (typically a factor 2x and 4x increase in drug concentration) and 1 lower concentration to maintain a living mutant lineage (always half the concentration that most recently produced growth). A 1/200 dilution was used to inoculate the next day’s evolution plate, and the process was repeated for a total of 8 days of selection. On the final day of evolution all strains were stocked in 30 percent glycerol. Strains were then plated on a pure BHI plate and a single colony was selected for IC_50_ determination. In the case of LZD mutants, days 2, 4, and 6 were also stocked for further testing.

### Measuring Drug Resistance and Sensitivity

Experiments to estimate IC_50_ were performed in replicate in 96-well plates by exposing mutants to a drug gradient consisting of 6-14 points-one per well—typically in a linear dilution series prepared in BHI medium with a total volume of 205 uL (200 uL of BHI, 5 uL of 1.5OD cells) per well. After 20 hours of growth the optical density at 600 nm (OD600) was measured using an Enspire Multimodal Plate Reader (Perkin Elmer) with an automated 20-plate stacker assembly. This process was repeated for all 60 mutants as well as the wild-type, which was measured in replicates of 8.

The optical density (OD600) measurements for each drug concentration were normalized by the OD600 in the absence of drug. To quantify drug resistance, the resulting dose response curve was fit to a Hill-like function *f* (*x*) = (1 + (*x/K*)^*h*^)^-1^ using nonlinear least squares fitting, where *K* is the half-maximal inhibitory concentration (IC_50_) and h is a Hill coefficient describing the steepness of the dose-response relationship. A mutant strain was defined to be collaterally sensitive if its IC_50_ had decreased by more than 3*σ_WT_* relative to the ancestral strain (*σ_WT_* is defined as the uncertainty-standard error across replicates-of the IC_50_ measured in the ancestral strain). Similarly, an increase in IC_50_ by more than 3*σ_WT_* relative to the ancestral strain corresponds to cross-resistance.

### Hierarchical clustering

Hierarchical clustering was performed in Matlab using, as input, the collateral profiles 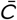 for each mutant. The distance between each pair of mutants was calculated using a correlation metric (Matlab function pdist with parameter ‘correlation’), and the linkage criteria was chosen to be the mean average linkage clustering.

### Markov decision process (MDP) model

The MDP model consists of a finite set of states (S), a finite set of actions (A), a conditional probability (*P_a_*(*s*’|*s, a*)) describing (action-dependent) Markovian transitions between these states, and an instantaneous reward function (*R_a_*(*s*)) associated with each state and action combination. The state of the system *s* ∈ *S* is an-dimensional vector, with the number of drugs and each component *s^i^* ∈ {*r_min_, r_min_* + 1,…, *r_max_*} indicating the level of resistance to drug *i*. The action *a* ∈ *A* ≡ {1,2,…, *n_d_*} is the choice of drug at the current step, and we take the reward function *R_a_* (s) to be the (negative ofthe) resistance level to the currently applied drug (i.e. the *a*-th component of *s*). The goal of the MDP is to identify a policy *π*(*s*), which is a mapping from *S* to *A* that specifies an optimal action for each state. The policy is chosen to maximize a cumulative reward function 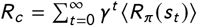, where *t* is the time step, *s_t_* is the state of the system at time *t, R_π_* (*s_t_*) is a random variable describing the instantaneous reward assuming that the actions are chosen according to policy *π*, and brackets indicate an expectation value. The parameter *α* (0 ≤ *α* < 1) is a discount factor that determines the relative importance of instantaneous vs long-term optimization. In words, we seek an optimal policy-which associates the resistance profile of a given population to an optimal drug choice-that minimizes the cumulative expected resistance to the applied drug.

The MDP problem was solved using value iteration, a standard dynamic programming algorithm for MDP models. Briefly, the optimization was performed by first computing the optimal value function *V*(*s*), which associates to each state *s* the expected reward obtained by following a particular policy and starting in that state. Following the well-established value iteration algorithm (80, 78, 79), we iterate according to *V*_*i*+1_ (*s*) = max_{*a*}_ (*R_a_*(*s*) + *γ* Σ_*s*‣_, *P*(*s*′|*s, a*)*V*(*s*′)). Given the optimal value function, the optimal policy is then given by the action that minimizes the optimal value function at the next time step.

Once the optimal policy *π* = *π*\* is found, the system is reduced to a simple Markov chain with transition matrix *T_π*_* = *P_π*(s)_*(*s*′|*s, π*\*(*s*)), where the subscript *π*\* means that the decision in each state is determined by the policy *π\** (i.e. that *a* = *π*\*(*s*) for a system in state *s*). Explicitly, the Markov chain dynamics are given by *P*_*t*+1_(*s*) = *T_π*_ P_t_*(*s*), with *P_t_*(*s*) the probability to be in state *s* at time step *t*. All quantities of interest-including P(Drug), P(Resist) (see Figure 5), and 〈*R*(*t*)〉-can be calculated directly from *P_t_*(*s*). For example, 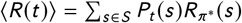, with *R_π*_*(*s*) the instantaneous reward for a system in state *s* under optimal policy *π*\*.

### Experiments to evaluate different drug sequence protocols

Experiments to evaluate different drug sequence protocols were performed in replicate in 96-well plates by exposing populations to antibiotic concentrations just above the wild-type MIC value, determined by an absence of measurable growth after 24 hours. Seed populations were grown overnight from single colonies and then diluted 1:200 into fresh BHI plates with appropriate antibiotic concentration according to each prescribed policy. Populations were left to grow inside a plate reader for 40 hours, while OD readings were taken every 20 minutes for at least the first 6 hours. To estimate daily growth, we took a final OD reading for each population after 40 hours. The populations were then diluted 1:200 into fresh BHI media, following a brief 2-hour outgrowth phase, populations were then diluted immediately into pre-prepared plates containing the appropriate drug concentrations. The purpose of the outgrowth phase is to minimize drug-drug interactions and post-antibiotic effects that may occur if the population were to be diluted into the next drug-plate immediately. To avoid contamination, each plate was covered during growth phase. In addition, each experimental plate contained 36 control wells with BHI alone — no cells. If any of these wells displayed visible growth, the plate was considered to be contaminated and discarded; the experiment was then started again from the previous night’s stock. During the 20 day experiment, only one such restart was required. Strains were stocked at −80C in 15 percent glycerol at the end of each 40 hour growth.

### Whole-genome sequencing

To identify any genomic changes that contributed to the measured collateral phenotypes identified, we sequenced 15 independently evolved drug mutants along with two V583 ancestors as well as a control V583 strain propagated in BHI for the 8 days. Each of the 15 drug-selected mutants and BHI-control were subjected to both clonal and population sequencing. Populations were streaked from a frozen stock, grown up in BHI, triple washed in PBS and DNA was isolated using a Quick-DNA Fungal/Bacterial Kit (Zymo Reserach). The clonal samples were sequenced in two batches via the University of Michigan sequencing core while the population samples were sequenced via the Microbial Genome Sequencing Center (MiGS) at University of Pittsburgh.

The resulting genomic data was analyzed using the high-throughput computational pipeline breseq, with default settings. Average read coverage depth was about 50 on batch 1, 300 on batch 2 and 200 on the population sequencing batch. Briefly, genomes were trimmed and subsequently aligned to E. faecalis strain V583 (Accession numbers: AE016830 – AE016833) via Bowtie 2. A sequence read was discarded if less than 90 percent of the length of the read did not match the reference genome or a predicted candidate junction. At each position a Bayesian posterior probability is calculated and the log10 ratio of that probability versus the probability of another base (A, T, C, G, gap) is calculated. Sufficiently high consensus scores are marked as read alignment evidence (in our case a consensus score of 10). Any mutation that occurred in either of the 2 control V583 strains was filtered from the results.

## Supporting information

SI_MaltasWood_Aug2019

## ACKNOWLEDGMENTS

This work is supported, in part, by the National Science Foundation (NSF No. 1553028 to KW) and the National Institutes of Health (NIH No. 1R35GM124875-01 to KW). The format for this preprint is adapted from the American Society for Microbiology (ASM) template available on Overleaf.com.

## SUPPLEMENTAL INFORMATION

**FIG S1.**
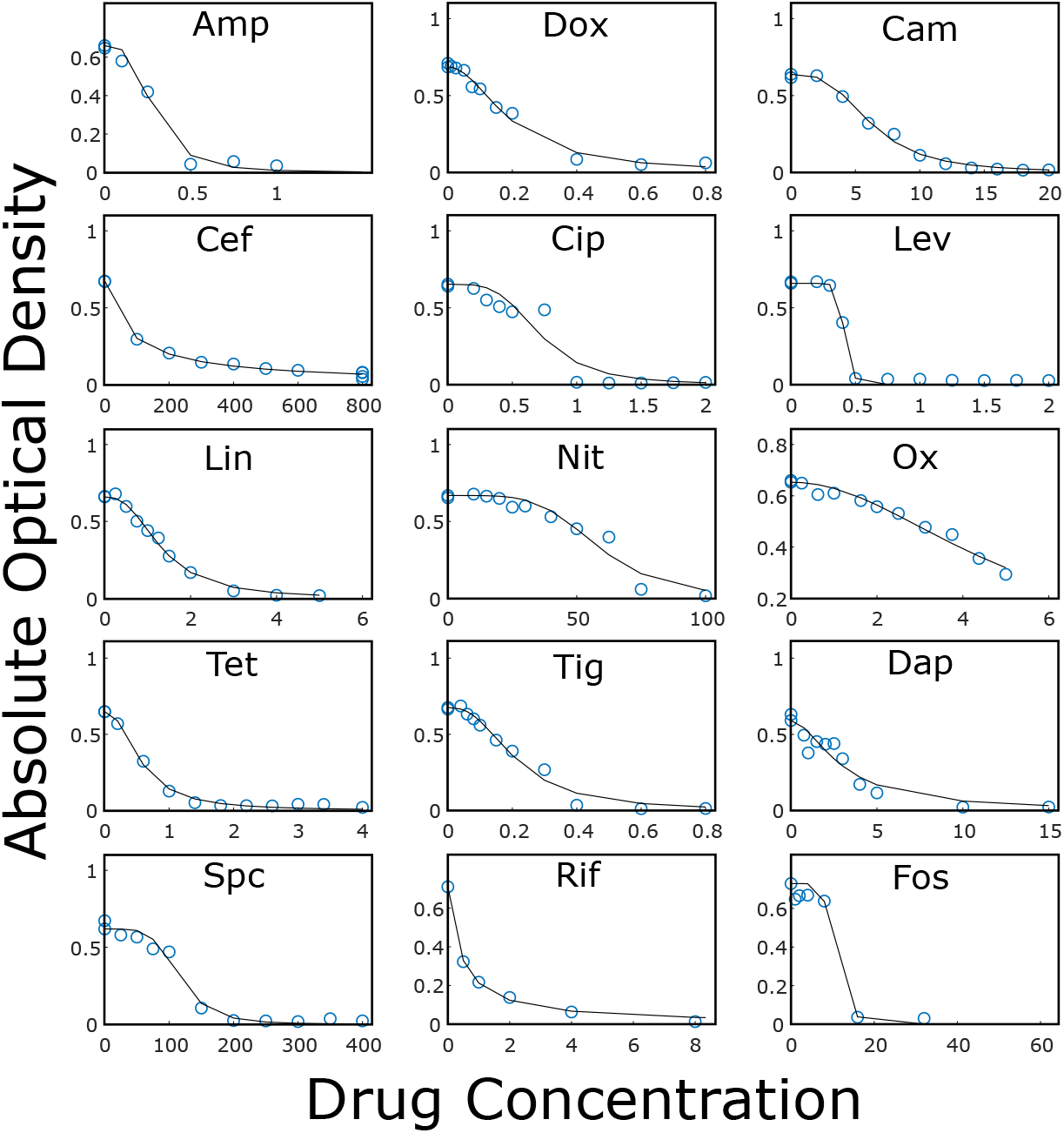
Example dose response curves for each drug Optical density (OD) of V583 cultures after 12 hours of incubation at various drug concentrations (blue circles). All drug concentrations are measured in *μ*g/mL. Lines: fit of normalized dose response curve to Hill-like function *f*(*x*) = (1 + (*x/K*)^*h*^)^-1^, with *K* the IC_50_ and *h* a Hill coefficient.

**FIG S2.**
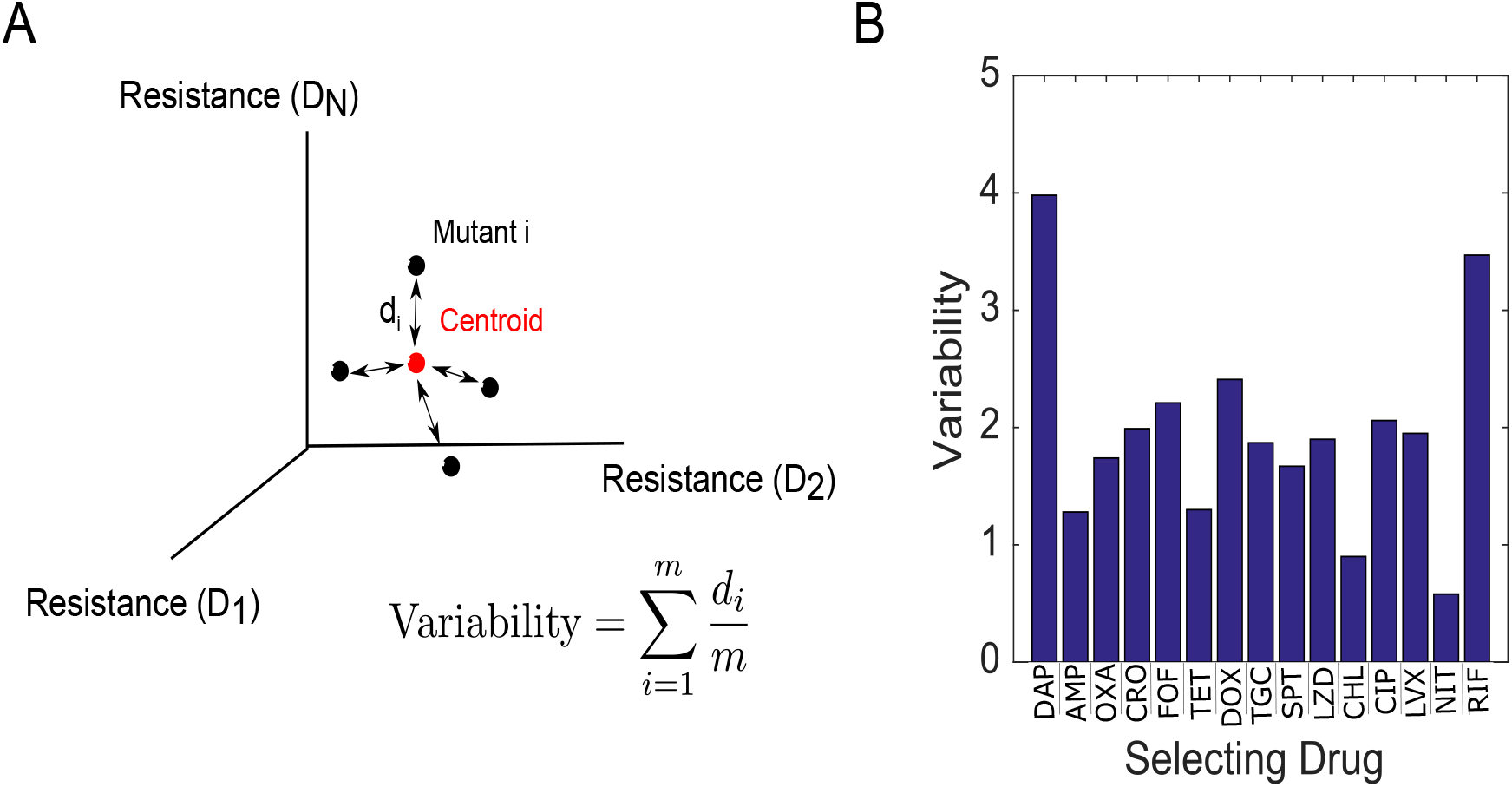
Variation within replicate populations. A. Variability in collateral profiles between mutants selected by the same drug is defined by first representing each mutant’s collateral profile as a vector 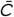 in 15-dimensional drug space. Dimension *i* represents the log2-scaled fold increase in IC_50_ (relative to wild-type) for drug *i*. The variability for a set of mutants evolved to the same drug is then given by the average Euclidean distance *d_i_* for a mutant from the centroid. B. Variability in replicates (defined in panel A) for all 15 drugs used for selection.

**TABLE 3.**
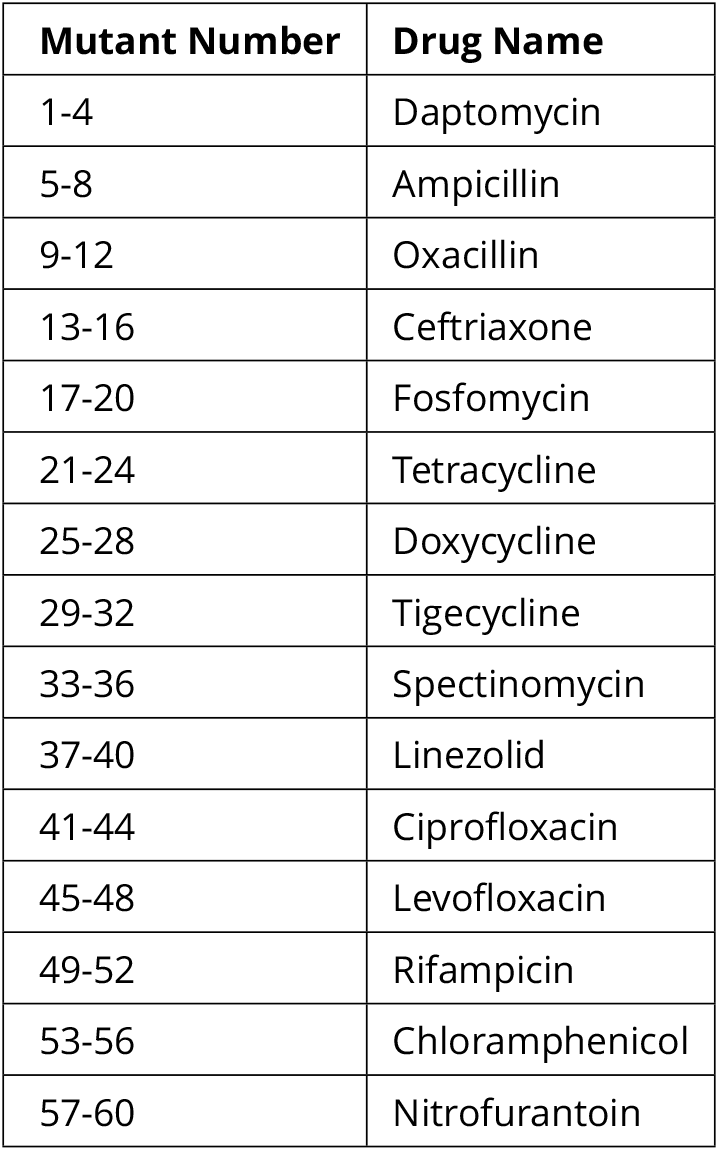
Mutant Number Table For Dendrograms

**FIG S3.**
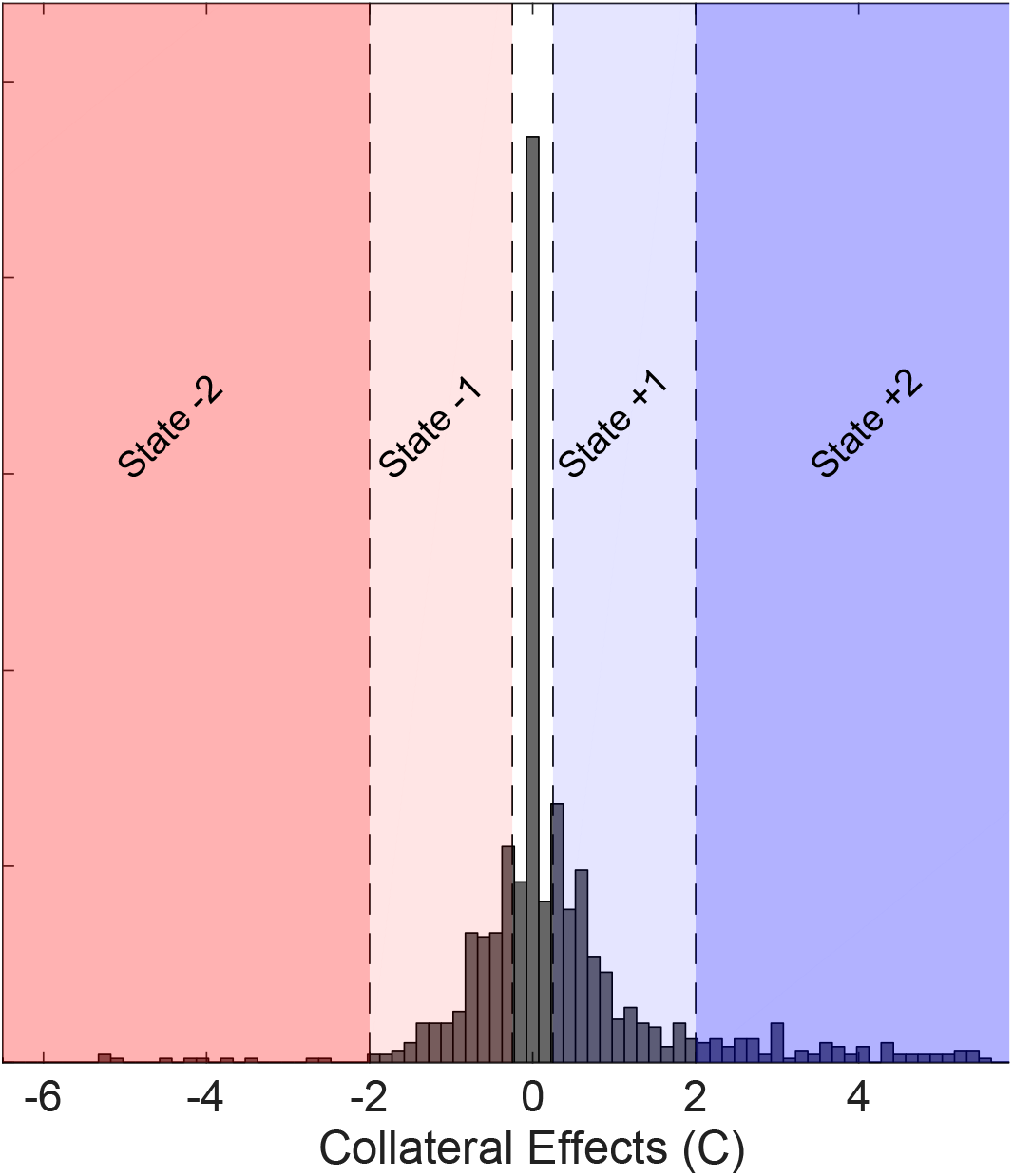
Discretization of collateral effects Histogram of collateral effects (*C* > 0 resistance, *C* < 0 sensitivity). Shaded regions indicate the five levels of discretization chosen for the MDP model (*C* < −2, red; −2 ≤ *C* < −0.25, light red; −0.25 ≤ *C* ≤ 0.25, white; 0.25 < *C* ≤ 2, light blue; *C* > 2, dark blue). The discretized values range from −2 (reducing resistance by two levels) to +2 (increasing resistance by two levels).

**FIG S4.**
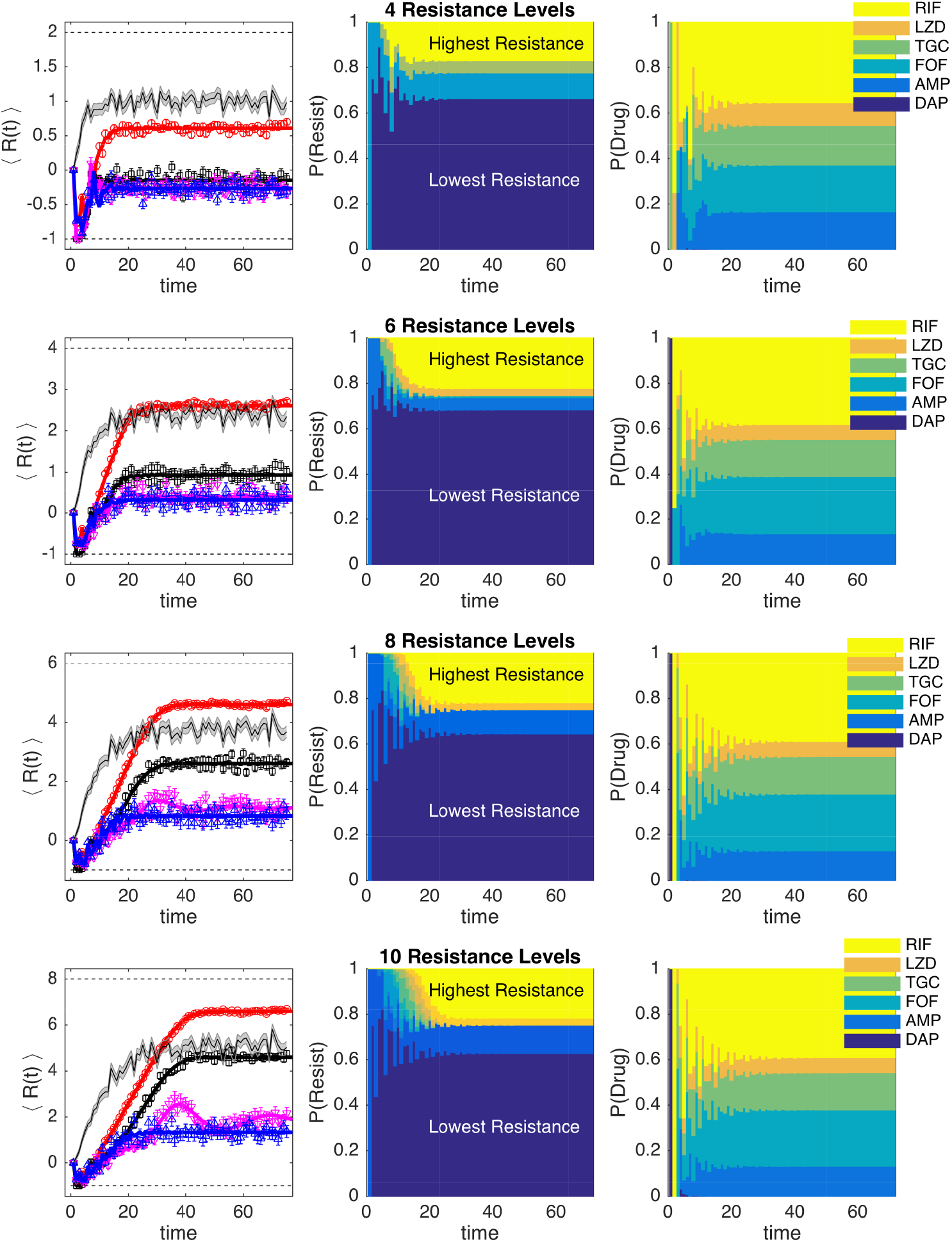
MDP models with different numbers of states show similar qualitative behavior In all panels, the MDP is solved for a selection of six drugs: daptomycin (DAP), ampicillin (AMP), fosfomycin (FOF), tigecycline (TGC), linezolid (LZD), and rifampicin (RIF). Left column: Average level of resistance (〈*R*(*t*)〉) to the applied drug for policies with *γ* = 0 (red), *γ* = 0.7 (black), *γ* = 0.9 (magenta), and *γ* = 0.99 (blue). Resistance to each drug is characterized by 4 (top row), 6, 8, or 10 (bottom row) discrete levels. At time 0, the population starts in the second lowest resistance level (0) for all drugs. Symbols (circles, triangles, squares) are the mean of 10^3^ independent simulations of the MDP, with error bars ± SEM. Solid lines are numerical calculations using exact Markov chain calculations (see Methods). Black shaded line, randomly cycled drugs. Middle column: The probability P(Resist) of the population exhibiting a particular level of resistance to the applied drug when the optimal policy (*γ* = 0.99) is used. Right column: The time-dependent probability P(Drug) of choosing each of the six drugs when the optimal policy (*γ* = 0.99) is used.

**FIG S5.**
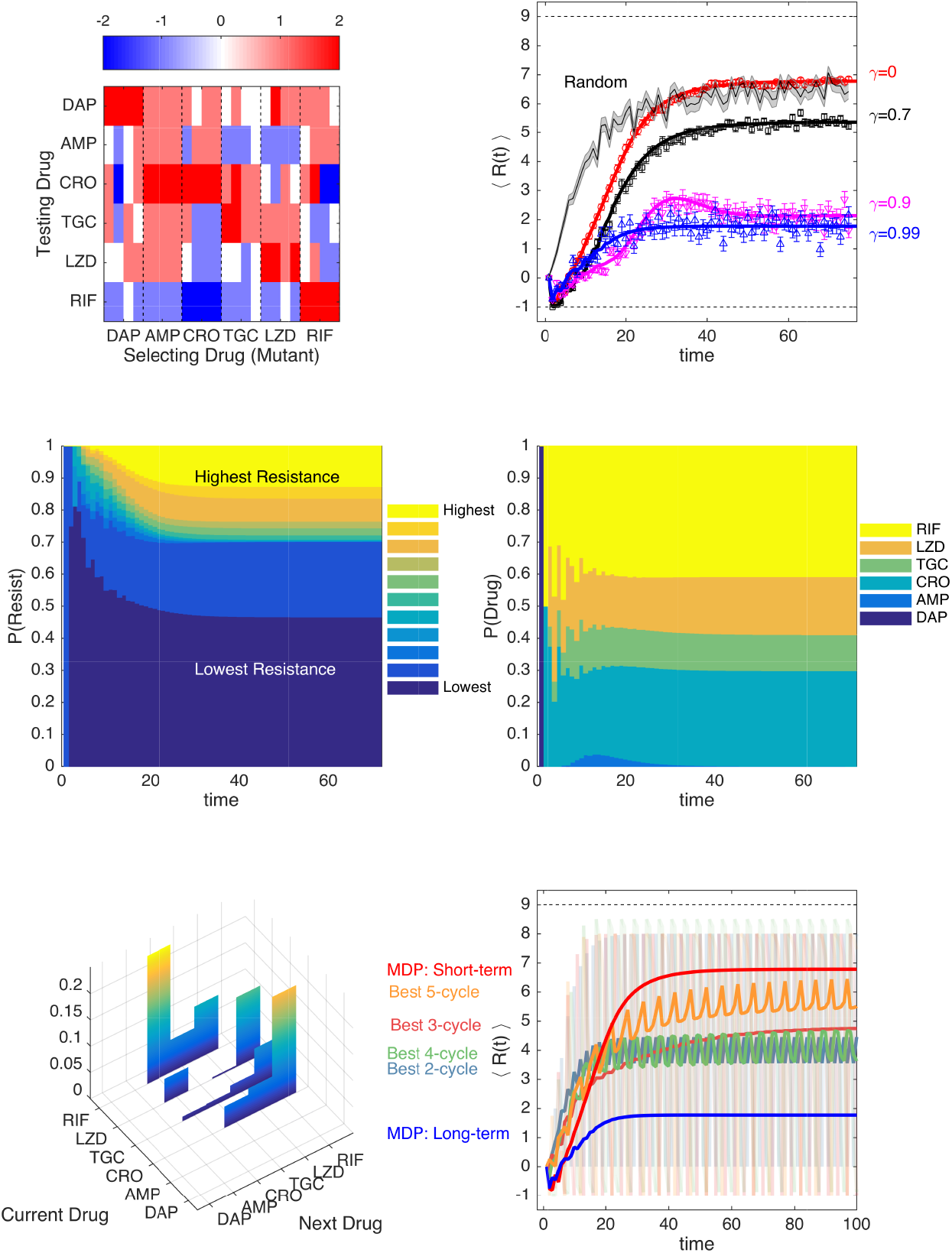
Optimal drug sequences constrain resistance on long timescales and outperform simple collateral sensitivitycycles A. Average ofdiscretized collateral sensitivity or resistance *C_d_* ∈ {−2, −1,0,1,2} for a selection of six drugs: daptomycin (DAP), ampicillin (AMP), ceftriaxone (CRO), tigecycline (TGC), linezolid (LZD), and ri-fampicin (RIF). For each selecting drug, the heat map shows the average value of *C_d_* from *n_r_* =4 independently evolved populations. See Fig 1 for original (non-discretized) data. B. Average level of resistance (〈*R*(*t*)〉) to the applied drug for policies with *γ* = 0 (red),*γ* = 0.7 (black), *γ* = 0.9 (magenta), and *γ* = 0.99 (blue). Resistance to each drug is characterized by11 discrete levels ranging from −1 (least resistant) to 9 (most resistant). At time 0, the population starts in the second lowest resistance level (0) for all drugs. Symbols (circles, triangles, squares) are the mean of 10^3^ independent simulations of the MDP, with error bars ± SEM. Solid lines are numerical calculations using exact Markov chain calculations (see Methods). Black shaded line, randomly cycled drugs. C. The probability P(Resist) of the population exhibitinga particular level of resistance to the applied drug when the optimal policy (*γ* = 0.99) is used. D. The time-dependent probability P(Drug) of choosing each of the six drugs when the optimal policy (*γ* = 0.99) is used. E. Steady state joint probability distribution P(current drug, next drug) for consecutive time steps when the optimal policy (*γ* = 0.99) is used. F. Average level of resistance (〈*R*(*t*)〉) to the applied drug for collateral sensitivity cycles of2 (dark green, CRO-RIF), 3 (pink, RIF-CRO-TGC), 4 (light green, TGC-LZD-AMP-RIF), and 5 (orange, AMP-RIF-CRO-TGC-LZD) drugs are compared with MDP policies with *γ* = 0 (short-term, red) and *γ* = 0.99 (long-term, blue). For visualizing the results of the collateral sensitivity cycles, which give rise to periodic behavior with large amplitude, the curves show a moving time average (window size 10 steps), but the smoothed curves are shown transparently in the background.

**FIG S6.**
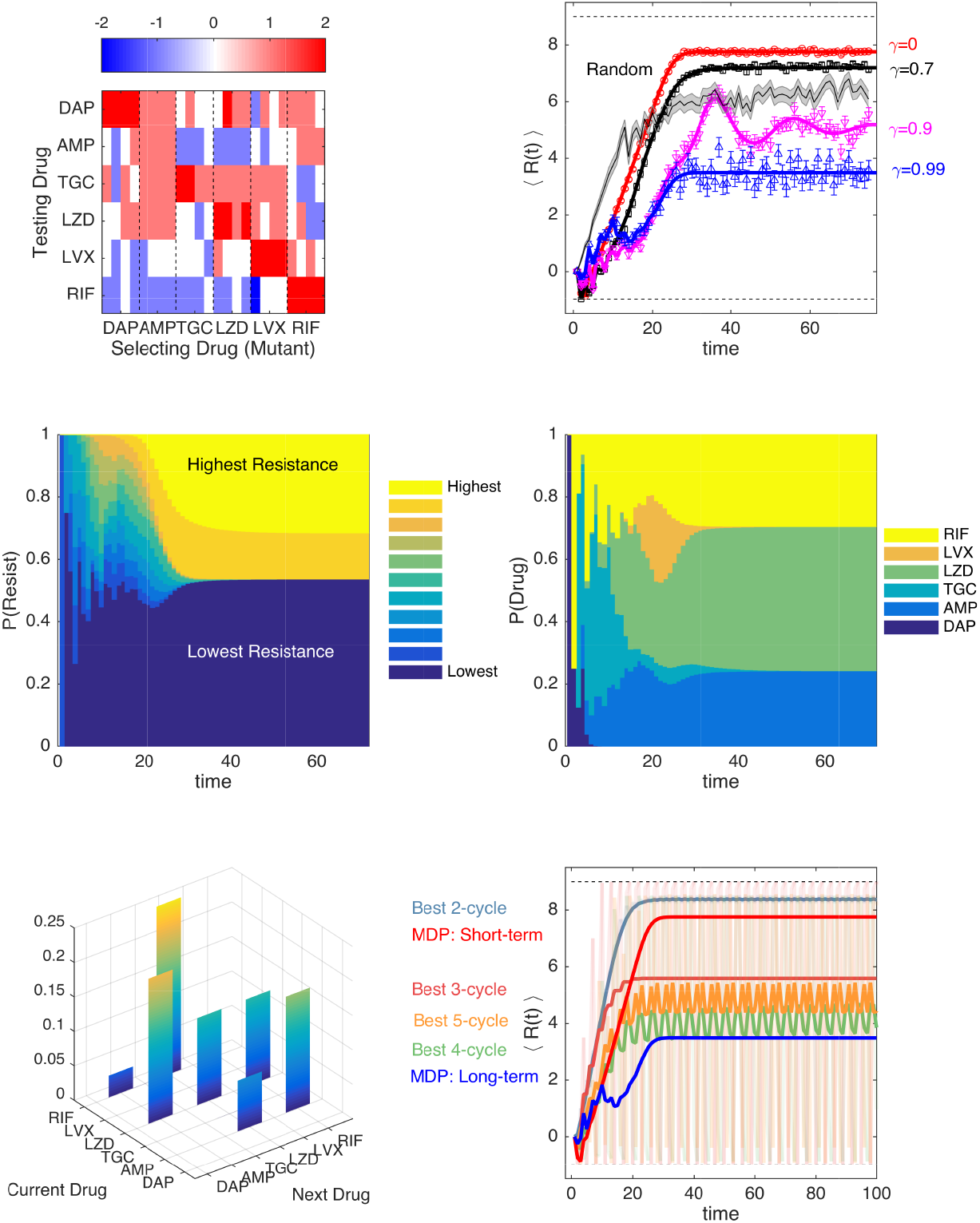
Optimal drug sequences constrain resistance on long timescales and outperform simple collateral sensitivitycycles A.Average ofdiscretized collateral sensitivity or resistance *C_d_* ∈ {−2, −1,0,1,2} for a selection of six drugs: daptomycin (DAP), ampicillin (AMP), tigecycline (TGC), linezolid (LZD), levofloxacin (LVX), and ri-fampicin (RIF). For each selecting drug, the heat map shows the average value of *C_d_* from *n_r_* =4 independently evolved populations. See Fig 1 for original (non-discretized) data. B. Average level of resistance (〈*R*(*t*)〉) to the applied drug for policies with *γ* = 0 (red), *γ* = 0.7 (black), *γ* = 0.9 (magenta), and *γ* = 0.99 (blue). Resistance to each drug is characterized by 11 discrete levels ranging from −1 (least resistant) to 9 (most resistant). At time 0, the population starts in the second lowest resistance level (0) for all drugs. Symbols (circles, triangles, squares) are the mean of 10^3^ independent simulations of the MDP, with error bars ± SEM. Solid lines are numerical calculations using exact Markov chain calculations (see Methods). Black shaded line, randomly cycled drugs. C. The probability P(Resist) of the population exhibiting a particular level of resistance to the applied drug when the optimal policy (*γ* = 0.99) is used. D. The time-dependent probability P(Drug) of choosing each of the six drugs when the optimal policy (*γ* = 0.99) is used. E. Steady state joint probability distribution P(current drug, next drug) for consecutive time steps when the optimal policy (*γ* = 0.99) is used. F. Average level of resistance (〈*R*(*t*)〉) to the applied drug for collateral sensitivity cycles of 2(dark green, TGC-RIF), 3 (pink, LZD-AMP-LVX), 4 (light green, RIF-TGC-LZD-AMP), and 5 (orange, AMP-LVX-RIF-TGC-LZD) drugs are compared with MDP policies with *γ* = 0 (short-term, red) and *γ* = 0.99 (long-term, blue). For visualizing the results of the collateral sensitivity cycles, which give rise to periodic behavior with large amplitude, the curves show a moving time average (window size 10 steps), but the smoothed curves are shown transparently in the background.

**FIG S7.**
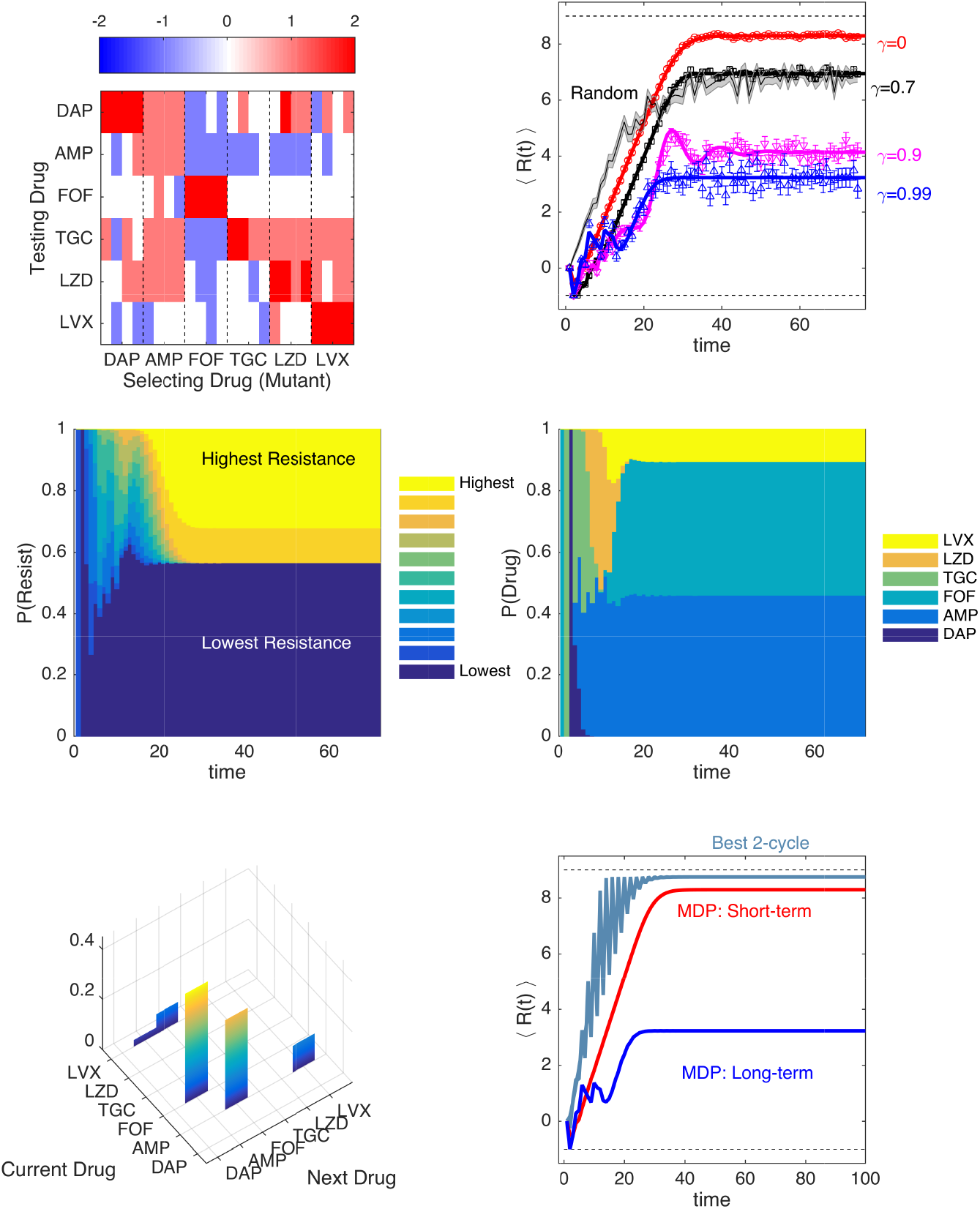
Optimal drug sequences constrain resistance on long timescales and outperform simple collateral sensitivity cycles A. Average of discretized collateral sensitivity or resistance *C_d_* ∈ {−2, −1,0,1,2} for a selection of six drugs: daptomycin (DAP), ampicillin (AMP), tigecycline(TGC), linezolid (LZD), levofloxacin (LVX), and rifampicin (RIF). For each selecting drug, the heat map shows the average value of *C_d_* from *n_r_* =4 independently evolved populations. See Fig 1 for original (non-discretized) data. B. Average level of resistance (〈*R*(*t*)〉) to the applied drug for policies with *γ* = 0 (red), *γ* = 0.7 (black), *γ* = 0.9 (magenta), and *γ* = 0.99 (blue). Resistance to each drug is characterized by 11 discrete levels ranging from −1 (least resistant) to 9 (most resistant). At time 0, the population starts in the second lowest resistance level (0) for all drugs. Symbols (circles, triangles, squares) are the mean of 10^3^ independent simulations of the MDP, with error bars ± SEM. Solid lines are numerical calculations using exact Markov chain calculations (see Methods). Black shaded line, randomly cycled drugs. C. The probability P(Resist) of the population exhibiting a particular level of resistance to the applied drug when the optimal policy (*γ* = 0.99) is used. D. The time-dependent probability P(Drug) of choosing each of the six drugs when the optimal policy (*γ* = 0.99) is used. E. Steady state joint probability distribution P(current drug, next drug) for consecutive time steps when the optimal policy (*γ* = 0.99) is used. F. Average level of resistance (〈*R*(*t*)〈) to the applied drug forcollateral sensitivity cycles of 2(darkgreen, AMP-LVX) drugs are compared with MDP policies with *γ* = 0 (short-term, red) and *γ* = 0.99 (long-term, blue). For visualizing the results of the collateral sensitivity cycles, which give rise to periodic behavior with large amplitude, the curves show a moving time average (window size 10 steps), butthe smoothed curves are shown transparently in the background.

**FIG S8.**
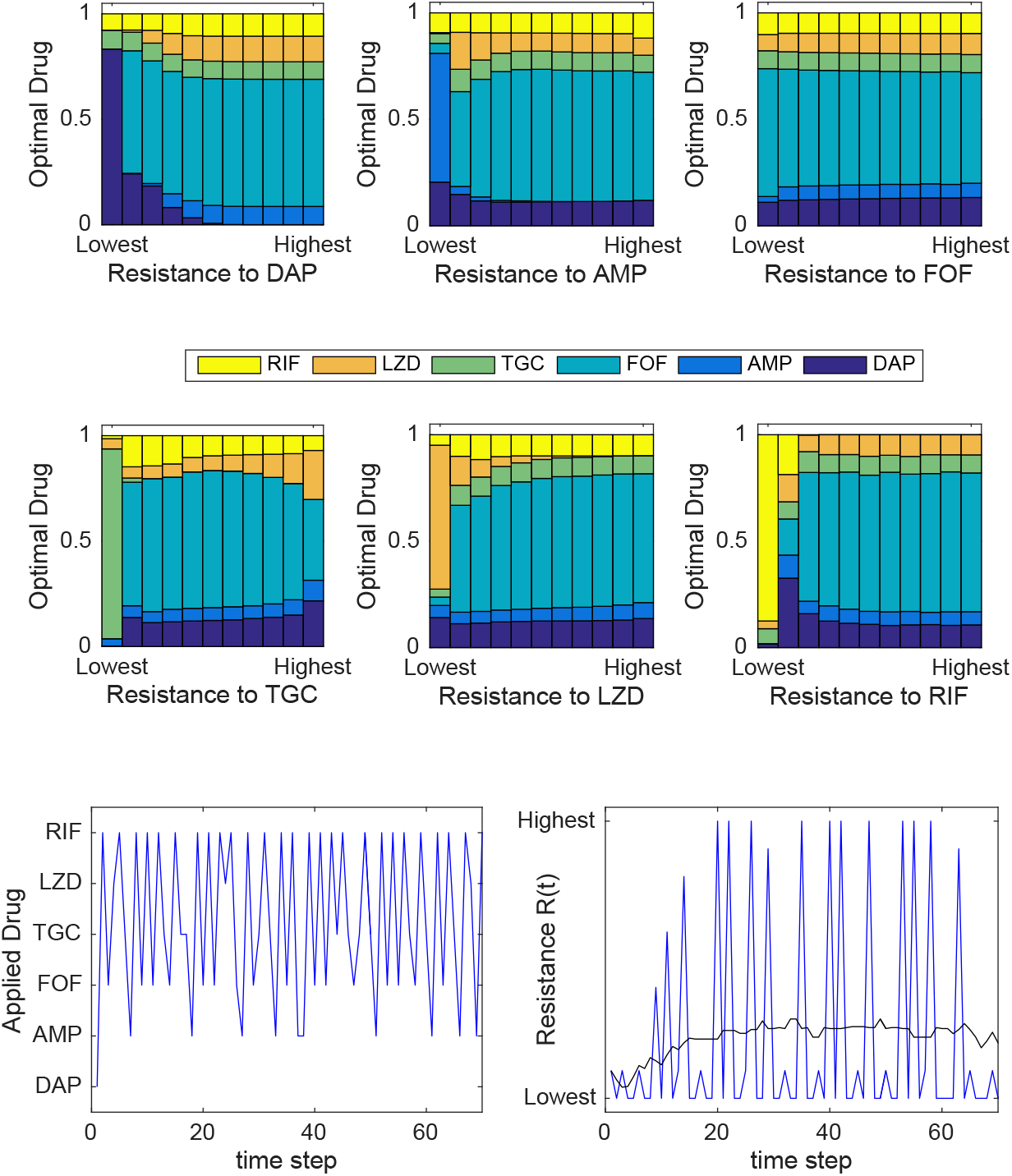
Optimal policy statistics and sample trajectories for *γ* = 0.99 The optimal policy *π*\*(*s*) is a mapping from the set of all possible resistance profiles (*S*) to the set of drugs (*A*). The policy associates each resistance profile with a unique (optimal) drug. Top panels: Frequency with which each drug is prescribed (according to the optimal policy) as a function of the level of resistance to an individual drug (horizontal axis). More specifically, for each of the six panels, the state space is partitioned into eleven distinct subsets, with each subset containing all states characterized by a given level of resistance to the particular drug in question (horizontal axis). The colored bars then show how frequently each of the six drugs is prescribed (according to the optimal policy) across all states within that subset. Bottom left panel: single simulated trajectory showing drug choice over time. Bottom right panel: single simulated trajectory of the instantaneous reward *R*, which corresponds to the resistance level to the applied drug. Blue curve is the specific trajectory; black curve is a moving average of the trajectory with a window size of 20.

**FIG S9.**
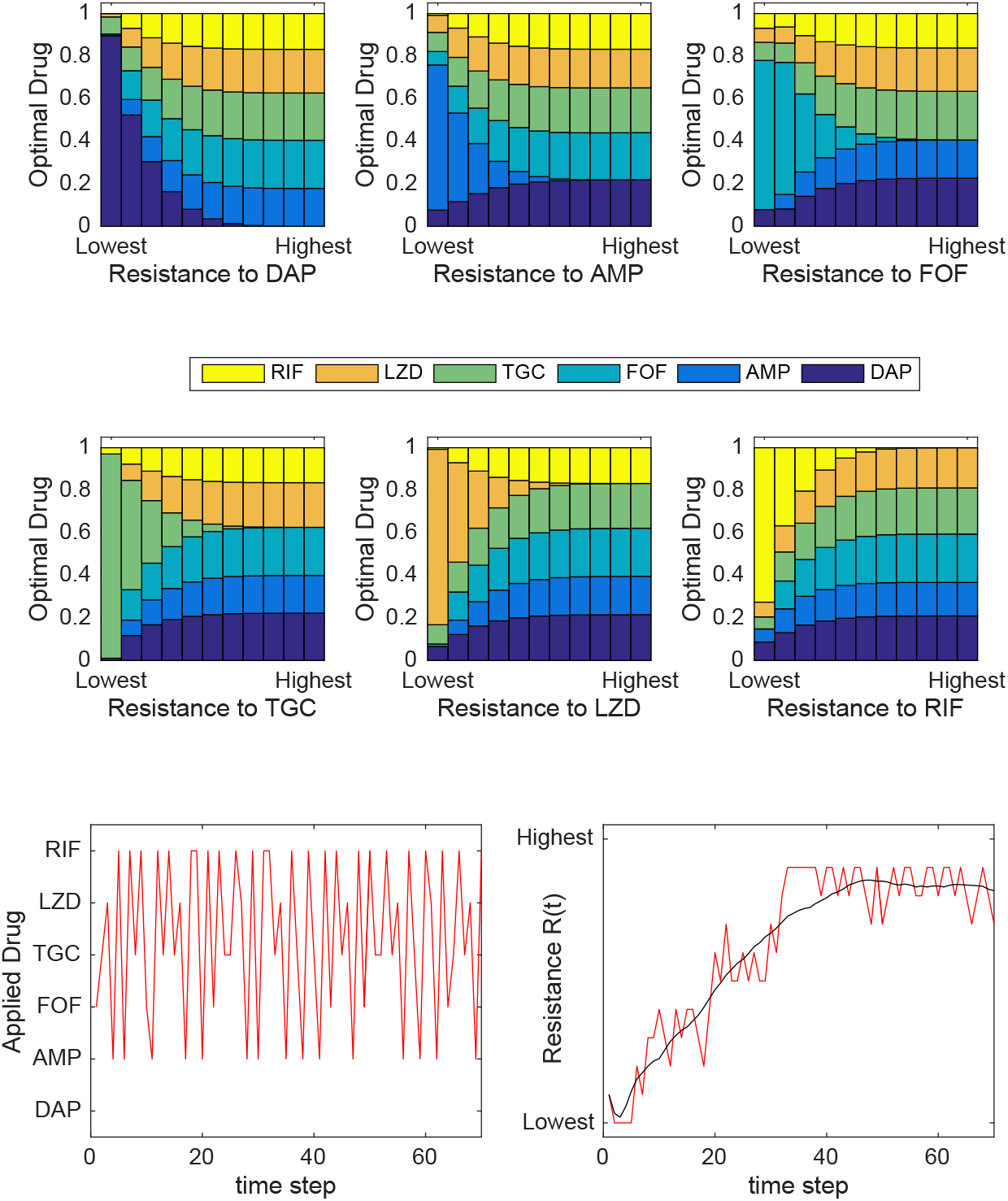
Optimal policy statistics and sample trajectories for *γ* = 0.1 Top panels: Frequency with which each drug is prescribed (according to the optimal policy) as a function of the level of resistance to an individual drug (horizontal axis). In each of the six panels, the state space is partitioned into eleven distinct subsets, with each subset containing all states with a given level of resistance to the particular drug in question. The colored bars then show how frequently each of the six drugs is prescribed across all states within that subset. Bottom left panel: single simulated trajectory showing drug choice over time. Bottom right panel: single simulated trajectory of the instantaneous reward *R*, which corresponds to the resistance level to the applied drug. Red curve is the specific trajectory; black curve is a moving average of the trajectory with a window size of 20.

**FIG S10.**
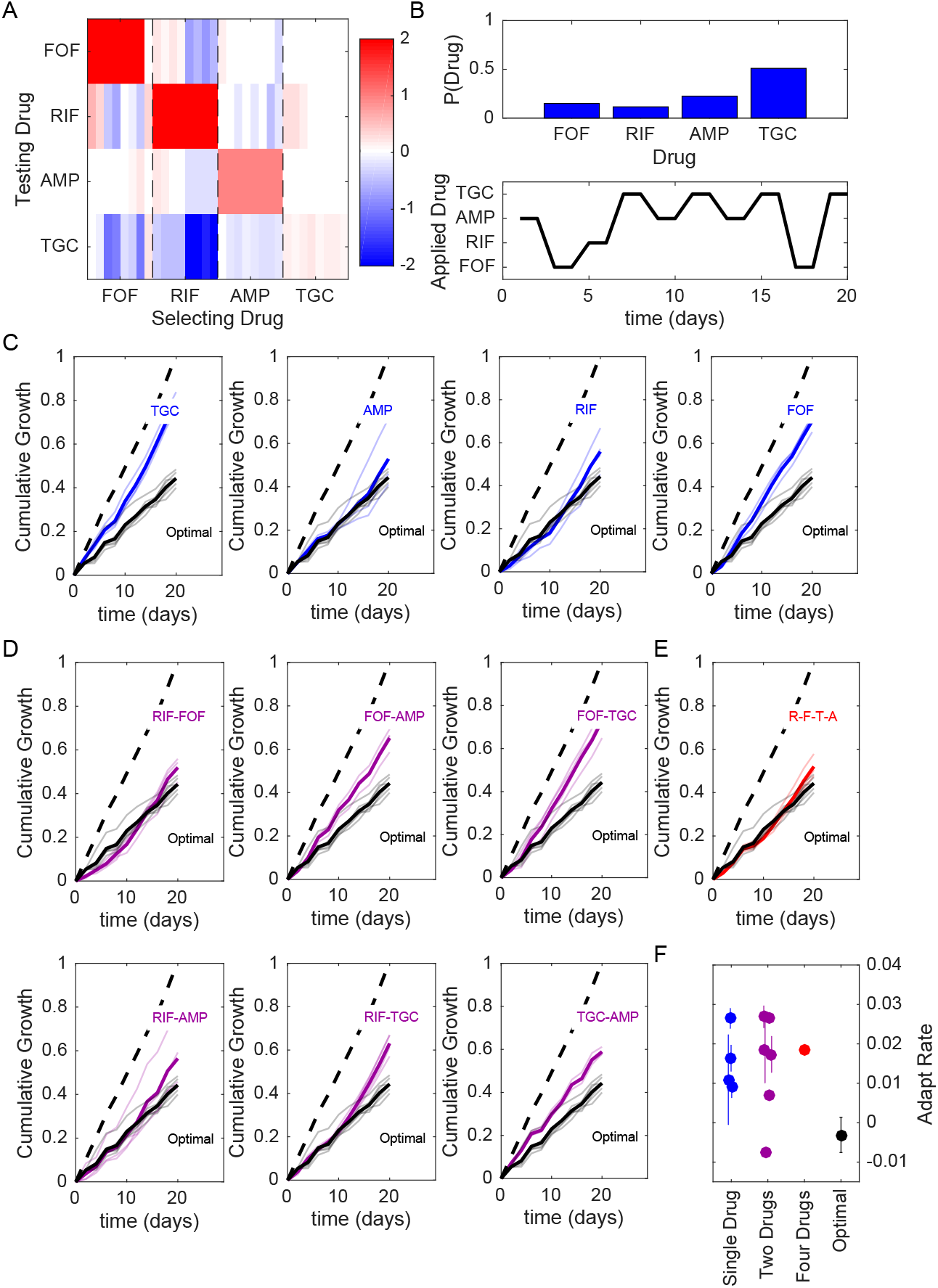
Optimized drug sequences reduce cumulative growth and adaptation rates in lab evolution experiments. A. Resistance (red) or sensitivity (blue) of each evolved mutant (horizontal axis; 4 drugs x 8 mutant per drug) to each drug (vertical axis) following 2 days of selection is quantified by the log_2_-transformed relative increase in the IC_50_ of the testing drug relative to that of wild-type (V583) cells. B. Top: distribution of applied drug at time step 20 (approximate steady state) calculated using an optimal policy with *γ* = 0.78. Bottom: sequence of applied drug from one particular realization of the stochastic process with the optimal policy (*γ* = 0.78). C-E. Cumulative population growth over time for populations exposed to single drug sequences (C, blue), two-drug sequences (D,magenta), a four drug sequence (E, red), or the optimal sequence from panel B (black curves, all panels). Transparent lines represent individual replicate experiments and each thicker dark line corresponds to a mean over replicates. Dashed line, drug-free control (normalized to a growth of 1 at the end of the experiment). F. Adaptation rate for single drug (blue), two-drug (magenta), four drug (red), and optimal sequences (black). Error bars are standard errors across replicates. Adaptation rate is defined as the slope of the best fit linear regression describing time series of daily growth (see Figure S14).

**FIG S11.**
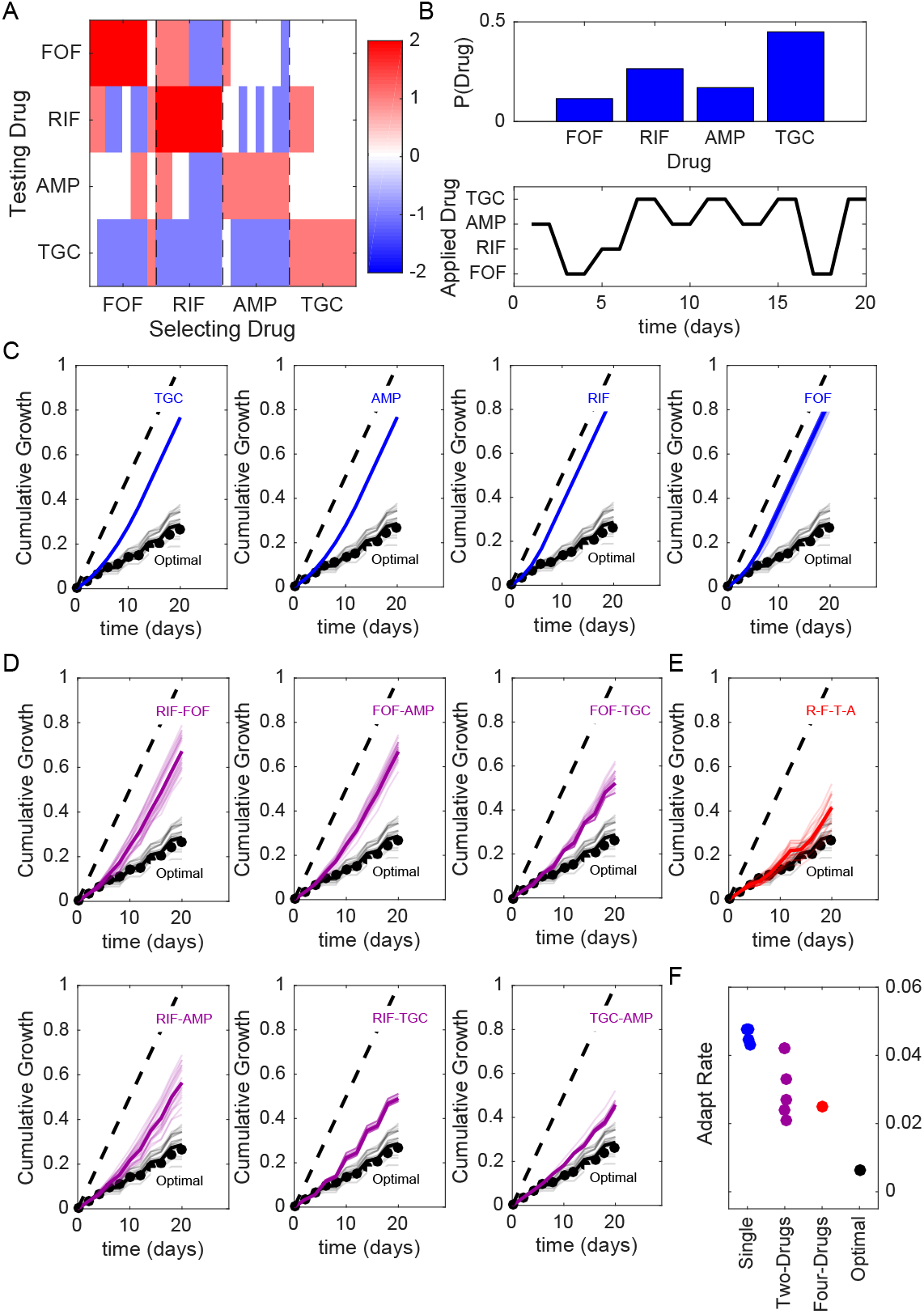
Optimized drug sequences reduce cumulative growth and adaptation rates in numerical simulations of the laboratory evolution experiments. Compare to Figure 6 (main text). A. Resistance (red) or sensitivity (blue) of each evolved mutant (horizontal axis; 4 drugs x 8 mutant per drug) to each drug (vertical axis) following 2 days of selection is quantified by the log_2_-transformed relative increase in the IC_50_ of the testing drug relative to that of wild-type (V583) cells. The profile is then discretized into 4 levels of resistance. B. Top: distribution of applied drug at time step 20 (approximate steady state) calculated using an optimal policy with *γ* = 0.9. Bottom: sequence of applied drug from one particular realization of the stochastic process with the optimal policy (*γ* = 0.9). C-E. Cumulative population growth (simulations) over time for populations exposed to single drug sequences (C, blue), two-drug sequences (D,magenta), a four drug sequence (E, red), or the optimal sequence from panel B (black curves, all panels). Black circles correspond to the true optimal (i.e. applying the MDP policy directly) and performs only slightly better, on average, than the fixed sequence in panel B. At each time step, resistance level to each drug is converted to an OD value using a linear conversion with the highest resistance level corresponding to growth of drug-free cells (OD≈ 0.6) and the lowest resistance level corresponding to OD=0. Transparent lines represent individual replicate experiments and each thicker dark line corresponds to a mean over replicates. Dashed line, drug-free control (normalized to a growth of 1 at the end of the experiment). F. Adaptation rate for single drug (blue), two-drug (magenta), four drug (red), and optimal sequences (black). Error bars are standard errors across replicates. Adaptation rate is defined as the slope of the best fit linear regression describing time series of daily growth.

**FIG S12.**
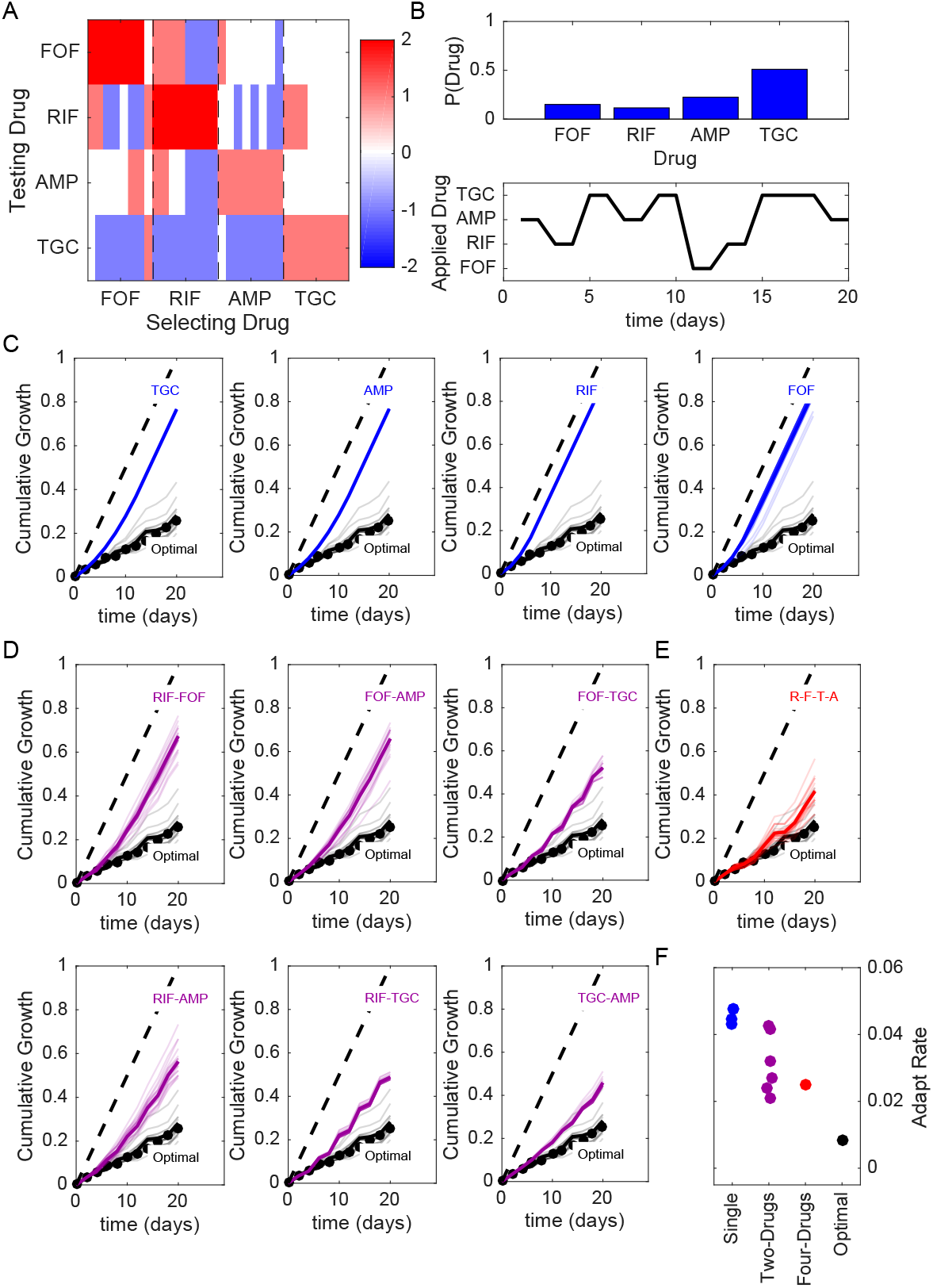
Optimized drug sequences reduce cumulative growth and adaptation rates in numerical simulations of the laboratory evolution experiments. Compare to Figure S10. A. Resistance (red) or sensitivity (blue) of each evolved mutant (horizontal axis; 4 drugs x 8 mutant per drug) to each drug (vertical axis) following 2 days of selection is quantified by the log_2_-transformed relative increase in the IC_50_ of the testing drug relative to that of wild-type (V583) cells. The profile is then discretized into 4 levels of resistance. B. Top: distribution of applied drug at time step 20 (approximate steady state) calculated using an optimal policy with *γ* = 0.78. Bottom: sequence of applied drug from one particular realization of the stochastic process with the optimal policy (*γ* = 0.78). C-E. Cumulative population growth (simulations) over time for populations exposed to single drug sequences (C, blue), two-drug sequences (D,magenta), a four drug sequence (E, red), or the optimal sequence from panel B (black curves, all panels). Black circles correspond to the true optimal (i.e. applying the MDP policy directly) and performs only slightly better, on average, than the fixed sequence in panel B. At each time step, resistance level to each drug is converted to an OD value using a linear conversion with the highest resistance level corresponding to growth of drug-free cells (OD≈ 0.6) and the lowest resistance level corresponding to OD=0. Transparent lines represent individual replicate experiments and each thicker dark line corresponds to a mean over replicates. Dashed line, drug-free control (normalized to a growth of 1 at the end of the experiment). F. Adaptation rate for single drug (blue), two-drug (magenta), four drug (red), and optimal sequences (black). Error bars are standard errors across replicates. Adaptation rate is defined as the slope of the best fit linear regression describing time series of daily growth.

**FIG S13.**
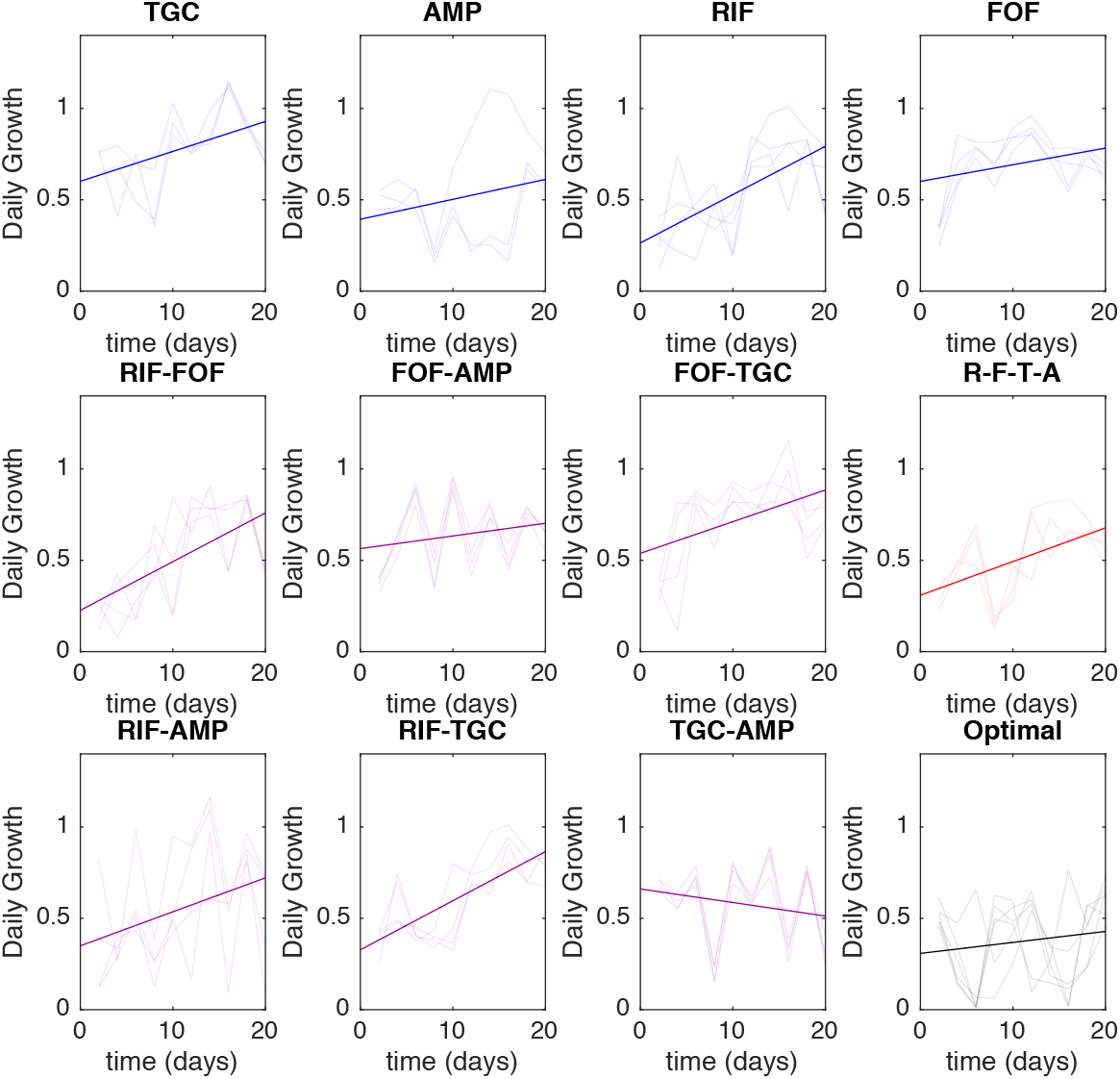
Estimated adaptation rate in lab evolution experiments based on *γ*=0.9 MDP policy. Daily growth, which is defined as the OD measured at the end of each 48-hour period (normalized to drug-free control), for populations exposed to single drug (blue), two-drug (magenta), four-drug (red), and optimal (black) drug sequences. All time series start at day 2 (i.e. following 48 hours of adaption). Transaprent curves correspond to individual replicate experiments; solid dark lines show the (average) best fit linear regression. Adaptation rate is defined as the slope of the regression line.

**FIG S14.**
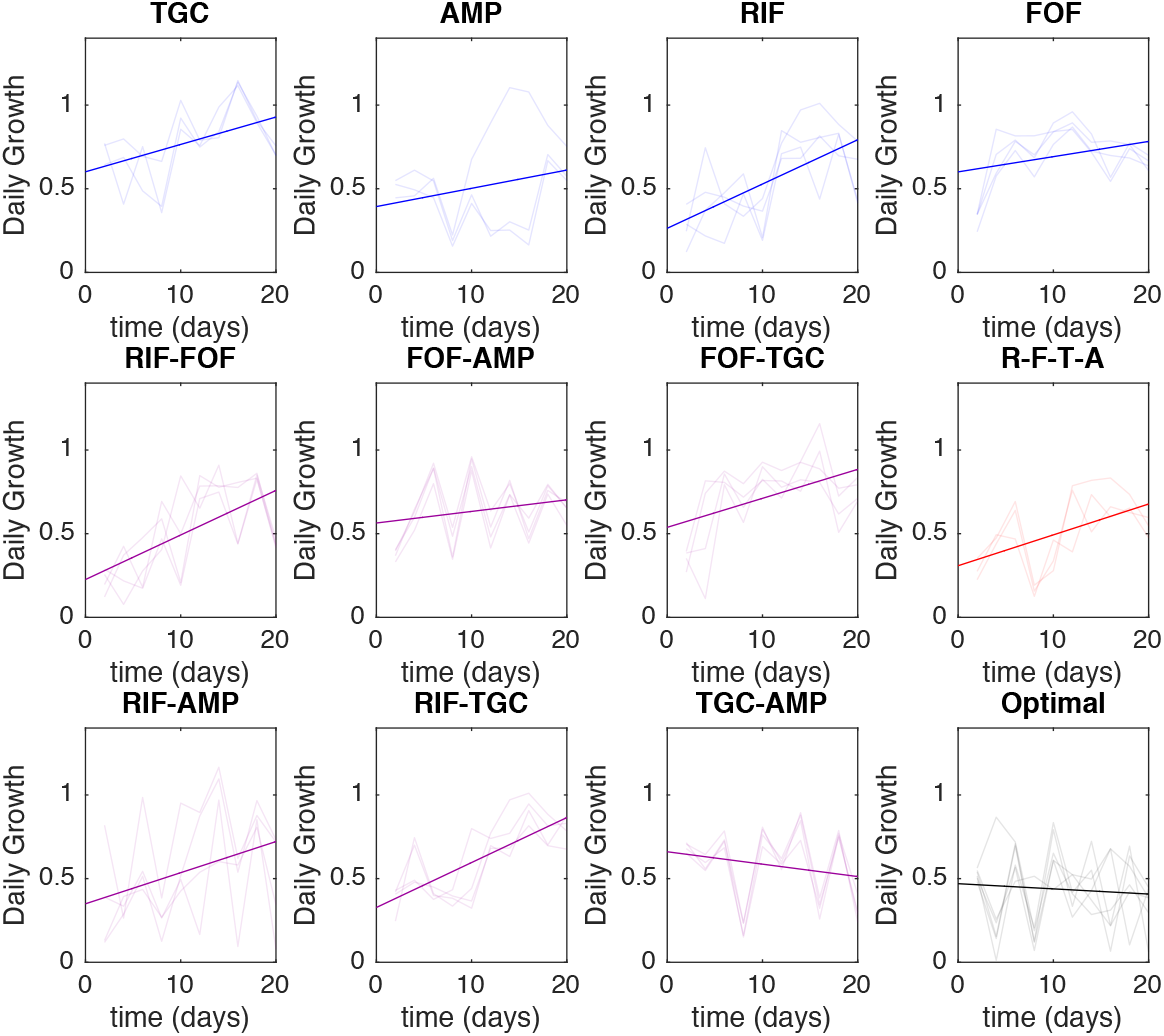
Estimated adaptation rate in lab evolution experiments based on *γ*=0.78 MDP policy. Daily growth, which is defined as the OD measured at the end of each 48-hour period (normalized to drug-free control), for populations exposed to single drug (blue), two-drug (magenta), four-drug (red), and optimal (black) drug sequences. All time series start at day 2 (i.e. following 48 hours of adaption). Transparent curves correspond to individual replicate experiments; solid dark lines show the (average) best fit linear regression. Adaptation rate is defined as the slope of the regression line.

## REFERENCES

1. Boucher HW, Talbot GH, Bradley JS, Edwards JE, Gilbert D, Rice LB, Scheld M, Spellberg B, Bartlett J. Bad bugs, no drugs; No ESAPE! An update from the Infections Diseases Society of America. Clin. Infect. Dis. 2009; 48:1–12.

2. Goldberg DE, Siliciana RF, Jr WRJ. Outwitting evolution: Fighting drug-resistant TB, malaria, and HIV. Cell 2012; 148:1271–1283.

3. Pfaller A. Antifungal drug resistance: Mechanisms, epidemiology, and consequences for treatment. Am.J. Med. 2012;125:S3–S13.

4. Raviglione M, Marais B, Floyd K, Lonnoroth K, Getahun H, Migliori GB, Harries AD, Nunn P, Lienhardt C, Graham S, Hakaya J, Weyer K, Cole S, Kaufmann SHE, Zumla A. Scaling up interentions to achieve global tuberculosis control: Progress and new developments. Lancet 2012; 379:1902–1913.

5. Borst P. Cancer drug pan-resistance: Pumps, cancer stem cells, quiescence, epithelial to mesenchymal transition, blocked cell death pathways, persister or what?. Open Biol. 2012;2.

6. Pluchino KM, Hall MD, Goldsborough AS, Callaghan R, Gotesman MM. Collateral sensitivity as a strategy against cancer multidrug resistance. Drug Resis. Updat. 2012;15:98–105.

7. Martin RB. Optimal control drug scheduling of cancer chemotherapy. J. Antimicrob. Chemother. 1992; 28:1113–1123.

8. Hansen E, Woods RJ, Read AF. How to Use a Chemotherapeutic Agent When Resistance to It Threatens the Patient. PLoS biology 2017; 15(2):e2001110.

9. Fischer A, Vázquez-García I, Mustonen V. The value of monitoring to control evolving populations. Proc. Natl. Acad. Sci. 2015; 112(4):1007–1012.

10. Feazel LM, Malhotra A, Perencevich EN, Kaboli P, Diekemea DJ, Schweizer ML. Effect of antibiotic stewardship programmes on *Clostridium difficile* incidence: a systematic review and meta-analysis. J. Antimicrob. Chemother. 2014;69:1748–1754.

11. Smith DL, Levin SA, Laxminarayan R. Strategic interactions in multi-institutional epidemics of antibiotic resistance. Proc. Natl. Acad. Sci. USA 2004;102:3153–3158.

12. Bergstrom CT, Lo M, Lipsitch M. Ecological theory suggests that antimicrobial cycling will not reduce antimicrobial resistance in hospitals. Proc. Natl. Acad. Sci. USA 2004;101:13285–13290.

13. Brown EM, Nathwani D. Antibiotic cycling or rotation: a systemic review of the evidence of efficacy. J. Antimicrob. Chemother. 2005;55:6–9.

14. Nichol D, Jeavons P, Fletcher AG, Bonomo RA, Maini PK, Paul JL, Gatenby RA, Anderson AR, Scott JG. Steering evolution with sequential therapy to prevent the emergence of bacterial antibiotic resistance. PLoS computational biology 2015;11(9):e1004493.

15. Baym M, Lieberman TD, Kelsic ED, Chait R, Gross R, Yelin I, Kishony R. Spatiotemporal microbial evolution on antibiotic landscapes. Science 2016;353(6304):1147–1151.

16. Zhang Q, Lambert G, Liao D, Kim H, Robin K, Tung Ck, Pourmand N, Austin RH. Acceleration of emergence of bacterial antibiotic resistance in connected microenvironments. Science 2011;333(6050):1764–1767.

17. DeJong MG, Wood KB. Tuning Spatial Profiles of Selection Pressure to Modulate the Evolution of Drug Resistance. Phys. Rev. Lett. 2018 Jun; 120:238102. https://link.aps.org/doi/10.1103/PhysRevLett.120.238102.

18. Yurtsev EA, Chao HX, Datta MS, Artemova T, Gore J. Bacterial cheating drives the population dynamics of cooperative antibiotic resistance plasmids. Mol. systems biology 2013;9(1):683.

19. Meredith HR, Lopatkin AJ, Anderson DJ, You L. Bacterial temporal dynamics enable optimal design of antibiotic treatment. PLoS computational biology 2015;11(4):e1004201.

20. Meredith HR, Srimani JK, Lee AJ, Lopatkin AJ, You L. Collective antibiotic tolerance: mechanisms, dynamics and intervention. Nat. chemical biology 2015; 11(3):182–188.

21. Vega NM, Gore J. Collective antibiotic resistance: mechanisms and implications. Curr. opinion microbiology 2014;21:28–34.

22. Balaban NQ, Merrin J, Chait R, Kowalik L, Leibler S. Bacterial persistence as a phenotypic switch. Science 2004;305(5690):1622–1625.

23. Tan C, Smith RP, Srimani JK, Riccione KA, Prasada S, Kuehn M, You L. The inoculum effect and band-pass bacterial response to periodic antibiotic treatment. Mol. systems biology 2012;8(1):617.

24. Karslake J, Maltas J, Brumm P, Wood KB. Population Density Modulates Drug Inhibition and Gives Rise to Potential Bistability of Treatment Outcomes for Bacterial Infections. PLoS computational biology 2016; 12(10):e1005098.

25. Torella JP, Chait R, Kishony R. Optimal drug synergy in antimicrobial treatments. PlOS Comput. Biol. 2010;6.

26. Michel J, Yeh PJ, Chait R, Jr RCM, Kishony R. Drug interactions modulate the potential for evolution of resistance. Proc. Natl. Acad. Sci. USA 2008; 105:14918–14923.

27. Hegreness M, Shoresh N, Damian D, Hartl D, Kishony R. Accelerated evolution of resistance in multi-drug environments. Proc. Natl. Acad. Sci. USA 2008;105:13977–13981.

28. Chait R, Craney A, Kishony R. Antibiotic interactions that select against resistance. Nature 2007;446(7136):668.

29. Baym M, Stone LK, Kishony R. Multidrug evolutionary strategies to reverse antibiotic resistance. Science 2016;351(6268):aad3292.

30. Wood K, Nishida S, Sontag ED, Cluzel P. Mechanism-independent method for predicting response to multidrug combinations in bacteria. Proc. Natl. Acad. Sci. 2012;109(30):12254–12259.

31. Zimmer A, Katzir I, Dekel E, Mayo AE, Alon U. Prediction of multidimensional drug dose responses based on measurements of drug pairs. Proc. Natl. Acad. Sci. 2016;113(37):10442–10447.

32. Zimmer A, Tendler A, Katzir I, Mayo A, Alon U. Prediction of drug cocktail effects when the number of measurements is limited. PLoS biology 2017;15(10):e2002518.

33. de Evgrafov MR, Gumpert H, Munck C, Thomsen TT, Sommer MOA. Collateral resistance and sensitivity modulate evolution in high-level resistance to drug combination treatment in *Staphylococcus aureus*. Mol. Biol. Evol. 2015;32:1175–1185.

34. Oz T, Guvenek A, Yildiz S, Karaboga E, Tamer YT, Mumcuyan N, Ozan VB, Senturk GH, Cokol M, Yeh P, Toprak E. Strength of selection pressure is an important parameter contributing to the complexity of antibiotic resistance evolution. Mol. Biol. Evol. 2014;31:2387–2401.

35. Lazar V, Nagy I, Spohn R, Csorgo B, Gyorkei A, Nyerges A, Horvath B, Voros A, Busa-Fekete R, Hrtyan M, Bogos B, Mehi O, Fekete G, Szap-panos B, Kegl B, Papp B, Pal C. Genome-wide analysis captures the determinants of the antibiotic cross-resistance interaction network. Nat. Commun. 2014;5.

36. Lazar V, Singh GP, Spohn R, Nagy I, Horvath B, Hrtyan M, Busa-Fekete R, Bogos B, Mehi O, Csorgo B, Posfai G, Fekete G, Szappanos B, Kegl B, Papp B, Pal C. Bacterial evolution and antibiotic hypersensitivity. Mol. Syst. Biol. 2013;9.

37. Lazar V, Martins A, Spohn R, Daruka L, Grezal G, Fekete G, Szamel M, Jangir PK, Kintses B, Csorgo B, Nyerges A, Gyorkei A, Kincses A, Der A, Walter FR, Deli MA, Urban E, Hegedus Z, Olajos G, Mehi O, Balint B, Nagy I, Martinek TA, Papp B, Pal C. Antibiotic-resistant bacteria show widespread collateral sensitivity to antimicrobial peptides. Nat. Microbiol. 2018;3:718–731.

38. Imamovic L, Ellabaan M, Machado A, Molin S, Johansen H, Sommer M. Drug-driven phenotypic convergence supports rational treatment strategies of chronic infections. Cell 2018;172:P121–134.

39. Shoval O, Sheftel H, Shinar G, Hart Y, Ramote O, Mayo A, Dekel E, Kavanagh K, Alon U. Evolutionary trade-offs, Pareto optimality, and the geometry of phenotype space. Science 2012;336(6085):1157–1160.

40. Hart Y, Sheftel H, Hausser J, Szekely P, Ben-Moshe NB, Korem Y, Tendler A, Mayo AE, Alon U. Inferring biological tasks using Pareto analysis of high-dimensional data. Nat. methods 2015;12(3):233–235.

41. Imamovic L, Sommer MOA. Use of collateral sensitivity networks to design drug cycling protocols that avoid resistance development. Sci. Transl. Med 2013;5:204ra132.

42. Kim S, Lieberman TD, Kishony R. Alternating antibiotic treatments constrain evolutionary paths to multidrug resistance. Proc. Natl. Acad. Sci. USA 2014;111:14494–14499.

43. Fuentes-Hernandez A, Plucain J, Gori F, Pena-Miller R, Reding C, Jansen G, Schulenburg H, Gudelj I, Beardmore R. Using a sequential regimen to eliminate bacteria at sublethal antibiotic dosages. PLoS biology 2015; 13(4):e1002104.

44. Roemhild R, Barbosa C, Beardmore RE, Jansen G, Schulenburg H. Temporal variation in antibiotic environments slows down resistance evolution in pathogenic Pseudomonas aeruginosa. Evol. applications 2015; 8(10):945–955.

45. Yoshida M, Reyes SG, Tsudo S, Horinouchi T, Furusawa C, Cronin L. Time-programmable dosing allows the manipulation, suppression and reversal of antibiotic drug resistance *in vitro.* Nat. Commun. 2017;8.

46. Roemhild R, Gokhale CS, Dirksen P, Blake C, Rosenstiel P, Traulsen A, Andersson DI, Schulenburg H. Cellular hysteresis as a principle to maximize the efficacy of antibiotic therapy. Proc. Natl. Acad. Sci. 2018; 115(39):9767–9772.

47. Munck C, Gumpert HK, Wallin AIN, Wang HH, Sommer MOA. Prediction of resistance development against drug components by collateral responses to component drugs. Sci. Transl. Med 2014;6:262ra156.

48. Barbosa C, Beardmore R, Schulenburg H, Jansen G. Antibiotic combination efficacy (ACE) networks for a Pseudomonas aeruginosa model. PLoS biology 2018;16(4):e2004356.

49. Barbosa C, Trebosc V, Kemmer C, Rosenstiel P, Beardmore R, Schulenburg H, Jansen G. Alternative evolutionary paths to bacterial antibiotic resistance cause distinct collateral effects. Mol. Biol. Evol. 2017; 34:2229–2244.

50. Dhawan A, Nichol D, Kinose F, Abazeed ME, Marusyk A, Haura EB, Scott JG. Collateral sensitivity networks reveal evolutionary instability and novel treatment strategies in ALK mutated non-small cell lung cancer. Sci. Reports 2017;7.

51. Nichol D, Rutter J, Bryant C, Hujer AM, Lek S, Adams MD, Jeavons P, Anderson AR, Bonomo RA, Scott JG. Antibiotic collateral sensitivity is contingent on the repeatability of evolution. Nat. communications 2019; 10(1):334.

52. Clewell DB, Gilmore MS, Ike Y, Shankar N. Enterococci: from commensals to leading causes of drug resistant infection. Massachusetts Eye and Ear Infirmary;2014.

53. Donlan RM. Biofilms and device-associated infections. Emerg. infectious diseases 2001; 7(2):277.

54. O’Driscoll T, Crank CW. Vancomycin-resistant enterococcal infections: epidemiology, clinical manifestations and optimal management. Drug Resis. Updat. 2015;8:217–230.

55. Cetinkaya Y, Falk P, Mayhall CG. Vancomycin Resistant Enterococci. Clin. Microbiol. Rev. 2000;13:686–707.

56. Huycke MM, Sahm DF, Gilmore MS. Multiple-drug resistant enterococci: the nature of the problem and an agenda for the future. Emerg. infectious diseases 1998;4(2):239.

57. Palmer KL, Daniel A, Hardy C, Silverman J, Gilmore MS. Genetic Basis for Daptomycin Resistance in Enterococci. Antimicrob. Agents Chemother. 2011;55(7):3345–3356. https://aac.asm.org/content/55/7/3345.

58. Miller C, Kong J, Tran TT, Arias CA, Saxer G, Shamoo Y. Adaptation of Enterococcus faecalis to daptomycin reveals an ordered progression to resistance. Antimicrob. agents chemotherapy 2013;p. AAC–01473.

59. Paulsen IT, Banerjei L, Myers G, Nelson K, Seshadri R, Read TD, Fouts DE, Eisen JA, Gill SR, Heidelberg J, et al. Role of mobile DNA in the evolution of vancomycin-resistant Enterococcus faecalis. Science 2003; 299(5615):2071–2074.

60. Zwietering M, Jongenburger I, Rombouts F, Van’tRiet K. Modeling of the bacterial growth curve. Appl. Environ. Microbiol. 1990;56(6):1875–1881.

61. Kelesidis T, Humphries R, Uslan DZ, Pegues DA. Daptomycin nonsus-ceptible enterococci: an emerging challenge for clinicians. Clin. Infect. Dis. 2011;52(2):228–234.

62. Tran TT, Munita JM, Arias CA. Mechanisms of drug resistance: daptomycin resistance. Annals New York Acad. Sci. 2015;1354(1):32–53.

63. Bhardwaj P, Hans A, Ruikar K, Guan Z, Palmer KL. Reduced chlorhexi-dine and daptomycin susceptibility in vancomycin-resistant Enterococcus faecium after serial chlorhexidine exposure. Antimicrob. agents chemotherapy 2018;62(1):e01235–17.

64. Bourgeois-Nicolaos N, Massias L, Couson B, Butel MJ, Andremont A, Doucet-Populaire F. Dose Dependence of Emergence of Resistance to Linezolid in Enterococcus faecalis In Vivo. The J. Infect. Dis. 2007 05; 195(10):1480–1488. https://doi.org/10.1086/513876.

65. Deatherage DE, Barrick JE. Identification of mutations in laboratory-evolved microbes from next-generation sequencing data using breseq. Methods Mol. Biol. 2014;1151:165–188.

66. Beabout K, Hammerstrom TG, Perez AM, Magalhães BF, Prater AG, Clements TP, Arias CA, Saxer G, Shamoo Y. The Ribosomal S10 Protein Is a General Target for Decreased Tigecycline Susceptibility. Antimicrob. Agents Chemother. 2015; 59(9):5561–5566. https://aac.asm.org/content/59/9/5561.

67. Criswell D, Tobiason VL, Lodmell JS, Samuels DS. Mutations Conferring Aminoglycoside and Spectinomycin Resistance in Borrelia burgdorferi. Antimicrob. Agents Chemother. 2006; 50(2):445–452. https://aac.asm.org/content/50/2/445.

68. Miller WR, Munita JM, Arias CA. Mechanisms of antibiotic resistance in enterococci. Expert. review anti-infective therapy 2014;12(10):1221–1236.

69. Li L, Wang Q, Zhang H, Yang M, Khan MI, Zhou X. Sensor histidine kinase is a β-lactam receptor and induces resistance to β-lactam antibiotics. Proc. Natl. Acad. Sci. 2016; 113(6):1648–1653. http://www.pnas.org/content/113/6/1648.

70. Kellogg SL, Kristich CJ. Convergence of PASTA Kinase and Two-Component Signaling in Response to Cell Wall Stress in Enterococcus faecalis. J. Bacteriol. 2018; 200(12). https://jb.asm.org/content/200/12/e00086-18.

71. Long KS, Vester B. Resistance to Linezolid Caused by Modifications at Its Binding Site on the Ribosome. Antimicrob. Agents Chemother. 2012; 56(2):603–612. https://aac.asm.org/content/56/2/603.

72. De Silva PM, Kumar A. Signal Transduction Proteins in Acinetobacter baumannii: Role in Antibiotic Resistance, Virulence, and Potential as Drug Targets. Front. Microbiol. 2019;10:49. https://www.frontiersin.org/article/10.3389/fmicb.2019.00049.

73. Pál C, Papp B, Lázár V. Collateral sensitivity of antibiotic-resistant microbes. Trends microbiology 2015;23(7):401–407.

74. Webber MA, Ricci V, Whitehead R, Patel M, Fookes M, Ivens A, Piddock LJV. Clinically Relevant Mutant DNA Gyrase Alters Supercoiling, Changes the Transcriptome, and Confers Multidrug Resistance. mBio 2013;4(4). https://mbio.asm.org/content/4M/e00273-13.

75. Gomez JE, Kaufmann-Malaga BB, Wivagg CN, Kim PB, Silvis MR, Renedo N, Ioerger TR, Ahmad R, Livny J. Ribosomal mutations promote the evolution of antibiotic resistance in a multidrug environment. eLife 2017;6.

76. Yeh P, Tschumi AI, Kishony R. Functional classification of drugs by properties of their pairwise interactions. Nat. Genet. 2006;38:489–494.

77. Podnecky NL, Fredheim EGA, Kloos J, Sorum V, Primicerio R, Roberts AP, Rozen DE, Samuelsen O, Johnsen PJ. Conserved collateral antibiotic susceptibility networks in diverse clinical strains of Escherichia coli. Nat. Commun. 2018;9.

78. Bellman RE, Dreyfus SE. Applied dynamic programming, vol. 2050. Princeton university press; 2015.

79. Feinberg EA, Shwartz A. Handbook of Markov decision processes: methods and applications, vol. 40. Springer Science & Business Media; 2012.

80. Bellman R. A Markovian Decision Process. J. Math. Mech. 1957; 6(5):679–684. http://www.jstor.org/stable/24900506.

81. Humphries RM, Pollett S, Sakoulas G. A current perspective on daptomycin for the clinical microbiologist. Clin. Microbiol. Rev. 2013;26:759–780.

82. Chow JW. Aminoglycoside Resistance in Enterococci. Clin. Infect. Dis. 2000 08;31(2):586–589. https://doi.org/10.1086/313949.

83. Zhao B, Sedlak JC, Srinivas R, Creixell P, Pritchard JR, Tidor B, Lauffen-burger DA, Hemann MT. Exploiting temporal collateral sensitivity in tumor clonal evolution. Cell. 2016; 165:1–13.

84. Barbosa C, Roemhild R, Rosenstiel P, Schulenburg H. Evolutionary stability of collateral sensitivity to antibiotics in the model pathogen Pseudomonas aeruginosa. bioRxiv 2019;p. 570663.

85. Jiao YJ, Baym M, Veres A, Kishony R. Population diversity jeopardizes the efficacy of antibiotic cycling. BioRxiv 2016;p. 082107.

86. Yu W, Hallinen KM, Wood KB. Interplay between antibiotic efficacy and drug-induced lysis underlies enhanced biofilm formation at subin-hibitory drug concentrations. Antimicrob. agents chemotherapy 2018; 62(1):e01603–17.

87. Martin M, Hölscher T, Dragoš A, Cooper VS, Kovács ÁT. Laboratory evolution of microbial interactions in bacterial biofilms. J. bacteriology 2016;198(19):2564–2571.

88. Steenackers HP, Parijs I, Foster KR, Vanderleyden J. Experimental evolution in biofilm populations. FEMS microbiology reviews 2016;40(3):373–397.

89. Turner CB, Marshall CW, Cooper VS. Parallel genetic adaptation across environments differing in mode of growth or resource availability. Evol. letters 2018;2(4):355–367.

90. Sahm DF, Kissinger J, Gilmore MS, Murray PR, Mulder R, Solliday J, Clarke B. In Vitro Susceptibility Studies of Vancomycin-Resistant Entero-coccus faecalis. Antimicrob. Agents Chemother. 1989 Sept;33(9):1588–1591.

