## Supplementary material for "Pervasive and diverse collateral sensitivity profiles inform optimal strategies to limit antibiotic resistance": SI_MaltasWood_Aug2019

**Summary** The Supplemental Information contains 14 figures (S1-S14) and 1 table.

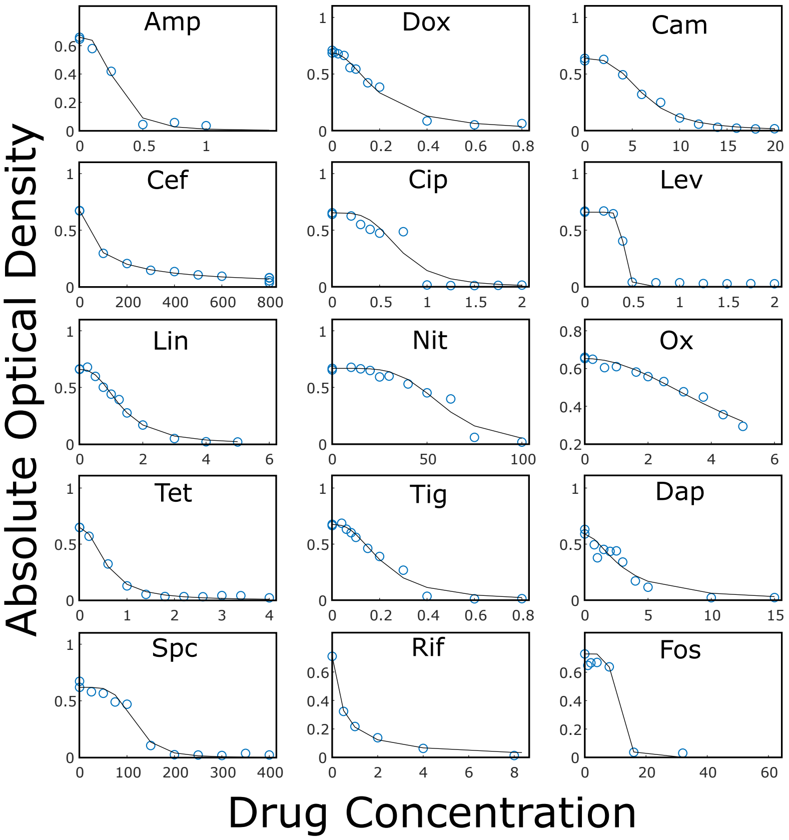


**FIG S1** **Example dose response curves for each drug Optical density (OD) of V583 cultures after 12 hours of incubation at various drug concentrations (blue circles). All drug concentrations are measured in $\mu$g/mL. Lines: fit of normalized dose response curve to Hill-like function $f(x) = (1 + (x/K)^h)^{-1}$, with $K$ the $IC_{50}$ and $h$ a Hill coefficient.**

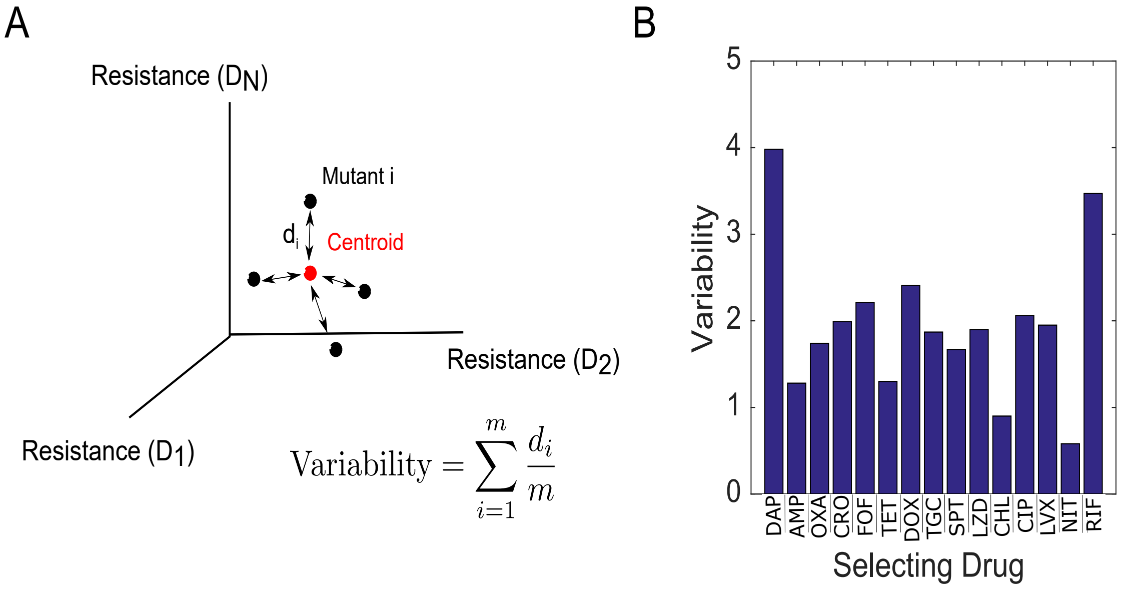


**FIG S2 Variation within replicate populations. A. Variability in collateral profiles between mutants selected by the same drug is defined by first representing each mutant's collateral profile as a vector $\bar{C}$ in 15-dimensional drug space. Dimension *i* represents the log2-scaled fold increase in $IC_{50}$ (relative to wild-type) for drug *i*. The variability for a set of mutants evolved to the same drug is then given by the average Euclidean distance $d_i$ for a mutant from the centroid. B. Variability in replicates (defined in panel A) for all 15 drugs used for selection.**

**TABLE 3** Mutant Number Table For Dendrograms

| Mutant Number | Drug Name |
|---|---|
| 1-4 | Daptomycin |
| 5-8 | Ampicillin |
| 9-12 | Oxacillin |
| 13-16 | Ceftriaxone |
| 17-20 | Fosfomycin |
| 21-24 | Tetracycline |
| 25-28 | Doxycycline |
| 29-32 | Tigecycline |
| 33-36 | Spectinomycin |
| 37-40 | Linezolid |
| 41-44 | Ciprofloxacin |
| 45-48 | Levofloxacin |
| 49-52 | Rifampicin |
| 53-56 | Chloramphenicol |
| 57-60 | Nitrofurantoin |

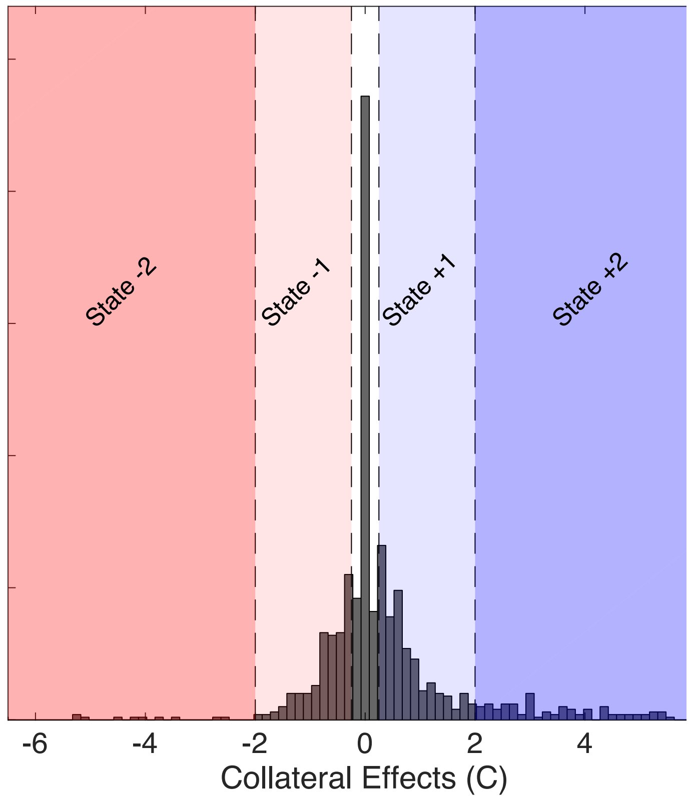


**FIG S3** **Discretization of collateral effects Histogram of collateral effects ($C > 0$ resistance, $C < 0$ sensitivity). Shaded regions indicate the five levels of discretization chosen for the MDP model ($C < -2$, red; $-2 \leq C < -0.25$, light red; $-0.25 \leq C \leq 0.25$, white; $0.25 < C \leq 2$, light blue; $C > 2$, dark blue). The discretized values range from -2 (reducing resistance by two levels) to +2 (increasing resistance by two levels).**

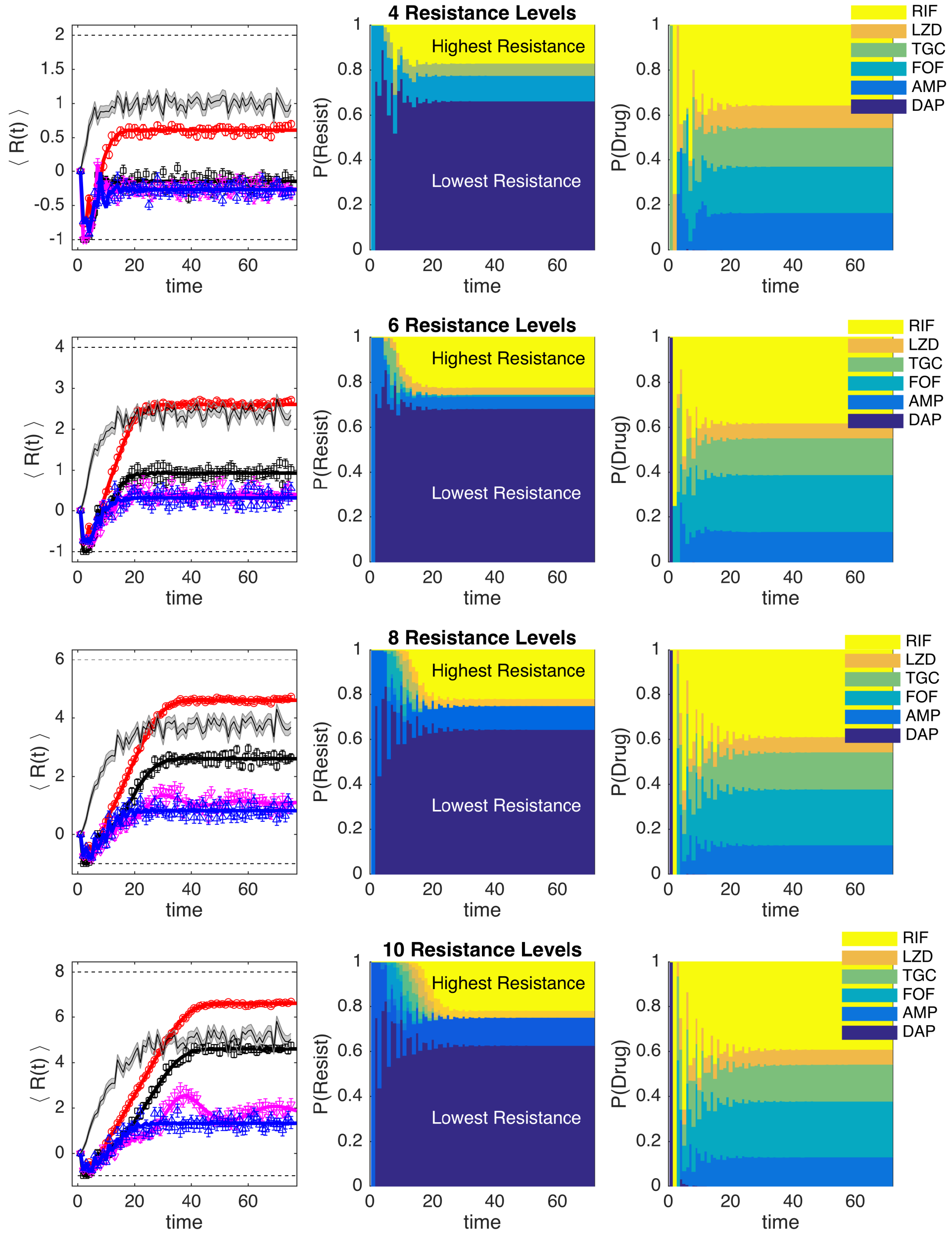


**FIG S4** **MDP models with different numbers of states show similar qualitative behavior In all panels, the MDP is solved for a selection of six drugs: daptomycin (DAP), ampicillin (AMP), fosfomycin (FOF), tigecycline (TGC), linezolid (LZD), and rifampicin (RIF). Left column: Average level of resistance ($\langle R(t) \rangle$) to the applied drug for policies with $\gamma = 0$ (red),$\gamma = 0.7$ (black),$\gamma = 0.9$ (magenta), and $\gamma = 0.99$ (blue). Resistance to each drug is characterized by 4 (top row), 6, 8, or 10 (bottom row) discrete levels. At time 0, the population starts in the second lowest resistance level (0) for all drugs. Symbols (circles, triangles, squares) are the mean of $10^3$ independent simulations of the MDP, with error bars ± SEM. Solid lines are numerical calculations using exact Markov chain calculations (see Methods). Black shaded line, randomly cycled drugs. Middle column: The probability P(Resist) of the population exhibiting a particular level of resistance to the applied drug when the optimal policy ($\gamma = 0.99$) is used. Right column: The time-dependent probability P(Drug) of choosing each of the six drugs when the optimal policy ($\gamma = 0.99$) is used.**

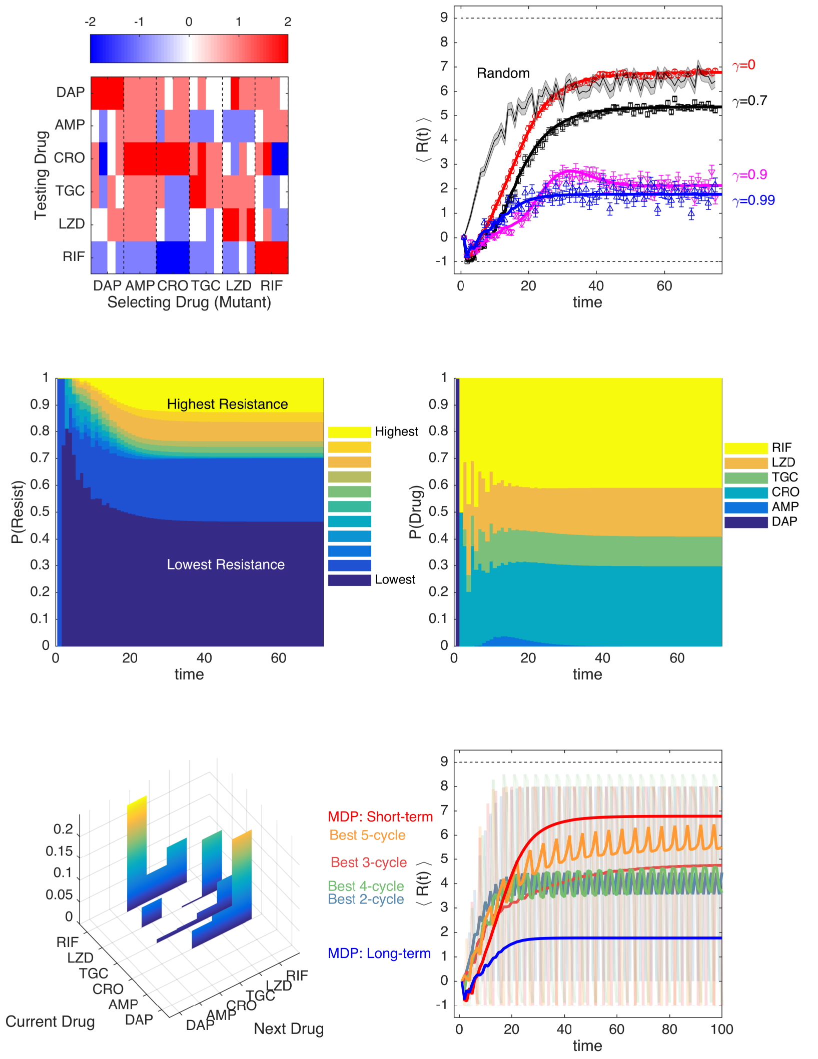


**FIG S5** **Optimal drug sequences constrain resistance on long timescales and outperform simple collateral sensitivity cycles A. Average of discretized collateral sensitivity or resistance $C_d \in \{-2, -1, 0, 1, 2\}$ for a selection of six drugs: daptomycin (DAP), ampicillin (AMP), ceftriaxone (CRO), tigecycline (TGC), linezolid (LZD), and rifampicin (RIF). For each selecting drug, the heat map shows the average value of $C_d$ from $n_r = 4$ independently evolved populations. See Fig 1 for original (non-discretized) data. B. Average level of resistance ($\langle R(t) \rangle$) to the applied drug for policies with $\gamma = 0$ (red),$\gamma = 0.7$ (black), $\gamma = 0.9$ (magenta), and $\gamma = 0.99$ (blue). Resistance to each drug is characterized by 11 discrete levels ranging from -1 (least resistant) to 9 (most resistant). At time 0, the population starts in the second lowest resistance level (0) for all drugs. Symbols (circles, triangles, squares) are the mean of $10^3$ independent simulations of the MDP, with error bars ± SEM. Solid lines are numerical calculations using exact Markov chain calculations (see Methods). Black shaded line, randomly cycled drugs. C. The probability P(Resist) of the population exhibiting a particular level of resistance to the applied drug when the optimal policy ($\gamma = 0.99$) is used. D. The time-dependent probability P(Drug) of choosing each of the six drugs when the optimal policy ($\gamma = 0.99$) is used. E. Steady state joint probability distribution P(current drug, next drug) for consecutive time steps when the optimal policy ($\gamma = 0.99$) is used. F. Average level of resistance ($\langle R(t) \rangle$) to the applied drug for collateral sensitivity cycles of 2 (dark green, CRO-RIF), 3 (pink, RIF-CRO-TGC), 4 (light green, TGC-LZD-AMP-RIF), and 5 (orange, AMP-RIF-CRO-TGC-LZD) drugs are compared with MDP policies with $\gamma = 0$ (short-term, red) and $\gamma = 0.99$ (long-term, blue). For visualizing the results of the collateral sensitivity cycles, which give rise to periodic behavior with large amplitude, the curves show a moving time average (window size 10 steps), but the smoothed curves are shown transparently in the background.**

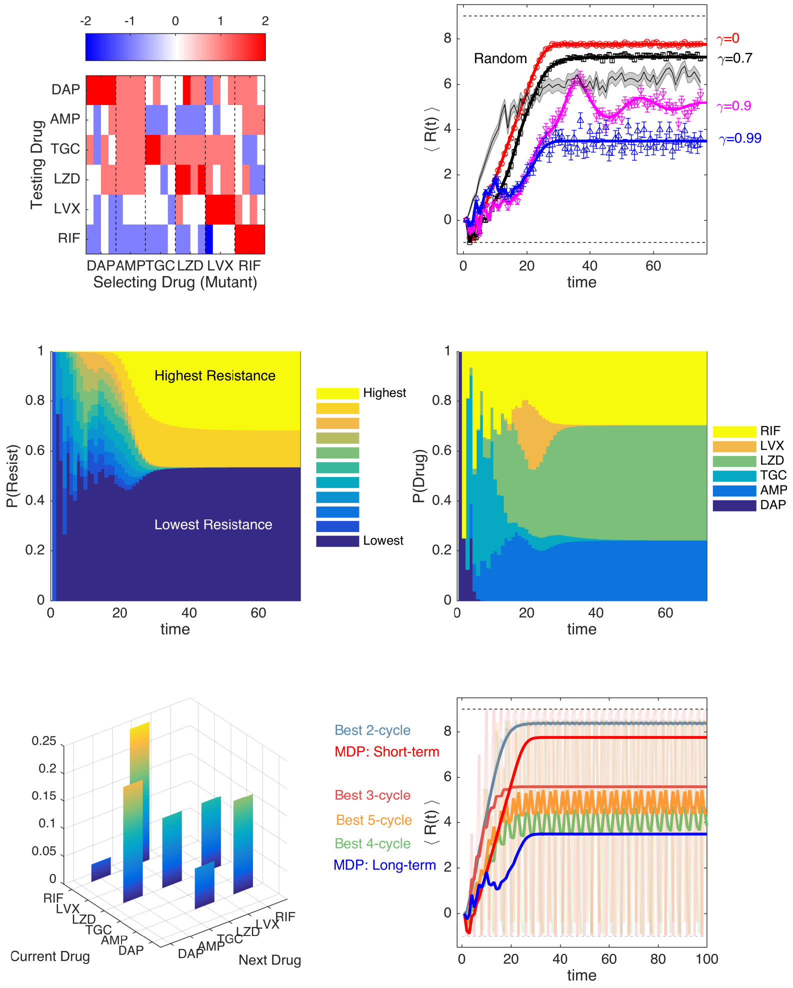


**FIG S6** **Optimal drug sequences constrain resistance on long timescales and outperform simple collateral sensitivity cycles A. Average of discretized collateral sensitivity or resistance $C_d \in \{-2,-1,0,1,2\}$ for a selection of six drugs: daptomycin (DAP), ampicillin (AMP), tigecycline (TGC), linezolid (LZD), levofloxacin (LVX), and rifampicin (RIF). For each selecting drug, the heat map shows the average value of $C_d$ from $n_r = 4$ independently evolved populations. See Fig 1 for original (non-discretized) data. B. Average level of resistance ($\langle R(t) \rangle$) to the applied drug for policies with $\gamma = 0$ (red),$\gamma = 0.7$ (black), $\gamma = 0.9$ (magenta), and $\gamma = 0.99$ (blue). Resistance to each drug is characterized by 11 discrete levels ranging from -1 (least resistant) to 9 (most resistant). At time 0, the population starts in the second lowest resistance level (0) for all drugs. Symbols (circles, triangles, squares) are the mean of $10^3$ independent simulations of the MDP, with error bars ± SEM. Solid lines are numerical calculations using exact Markov chain calculations (see Methods). Black shaded line, randomly cycled drugs. C. The probability P(Resist) of the population exhibiting a particular level of resistance to the applied drug when the optimal policy ($\gamma = 0.99$) is used. D. The time-dependent probability P(Drug) of choosing each of the six drugs when the optimal policy ($\gamma = 0.99$) is used. E. Steady state joint probability distribution P(current drug, next drug) for consecutive time steps when the optimal policy ($\gamma = 0.99$) is used. F. Average level of resistance ($\langle R(t) \rangle$) to the applied drug for collateral sensitivity cycles of 2 (dark green, TGC-RIF), 3 (pink, LZD-AMP-LVX), 4 (light green, RIF-TGC-LZD-AMP), and 5 (orange, AMP-LVX-RIF-TGC-LZD) drugs are compared with MDP policies with $\gamma = 0$ (short-term, red) and $\gamma = 0.99$ (long-term, blue). For visualizing the results of the collateral sensitivity cycles, which give rise to periodic behavior with large amplitude, the curves show a moving time average (window size 10 steps), but the smoothed curves are shown transparently in the background.**

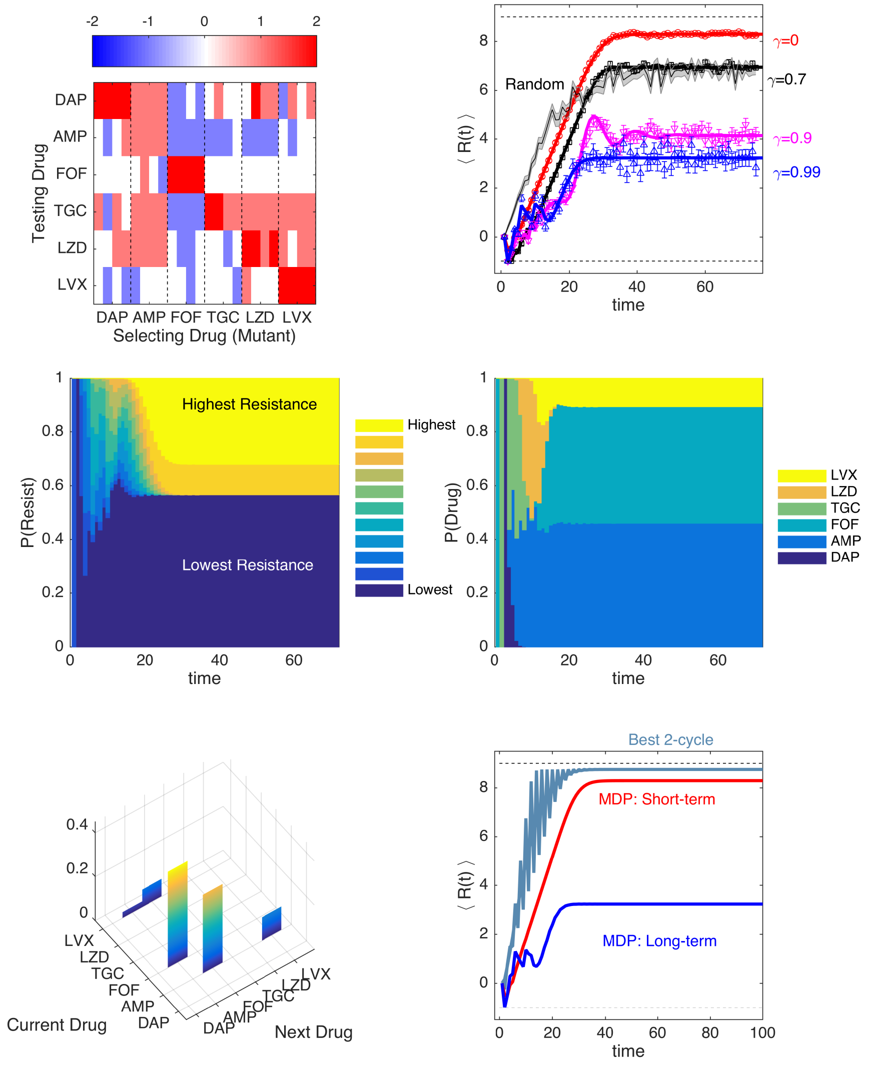


**FIG S7** **Optimal drug sequences constrain resistance on long timescales and outperform simple collateral sensitivity cycles A. Average of discretized collateral sensitivity or resistance $C_d \in \{-2,-1,0,1,2\}$ for a selection of six drugs: daptomycin (DAP), ampicillin (AMP), tigecycline (TGC), linezolid (LZD), levofloxacin (LVX), and rifampicin (RIF). For each selecting drug, the heat map shows the average value of $C_d$ from $n_r = 4$ independently evolved populations. See Fig 1 for original (non-discretized) data. B. Average level of resistance ($\langle R(t) \rangle$) to the applied drug for policies with $\gamma = 0$ (red),$\gamma = 0.7$ (black), $\gamma = 0.9$ (magenta), and $\gamma = 0.99$ (blue). Resistance to each drug is characterized by 11 discrete levels ranging from -1 (least resistant) to 9 (most resistant). At time 0, the population starts in the second lowest resistance level (0) for all drugs. Symbols (circles, triangles, squares) are the mean of $10^3$ independent simulations of the MDP, with error bars ± SEM. Solid lines are numerical calculations using exact Markov chain calculations (see Methods). Black shaded line, randomly cycled drugs. C. The probability P(Resist) of the population exhibiting a particular level of resistance to the applied drug when the optimal policy ($\gamma = 0.99$) is used. D. The time-dependent probability P(Drug) of choosing each of the six drugs when the optimal policy ($\gamma = 0.99$) is used. E. Steady state joint probability distribution P(current drug, next drug) for consecutive time steps when the optimal policy ($\gamma = 0.99$) is used. F. Average level of resistance ($\langle R(t) \rangle$) to the applied drug for collateral sensitivity cycles of 2 (dark green, AMP-LVX) drugs are compared with MDP policies with $\gamma = 0$ (short-term, red) and $\gamma = 0.99$ (long-term, blue). For visualizing the results of the collateral sensitivity cycles, which give rise to periodic behavior with large amplitude, the curves show a moving time average (window size 10 steps), but the smoothed curves are shown transparently in the background.**

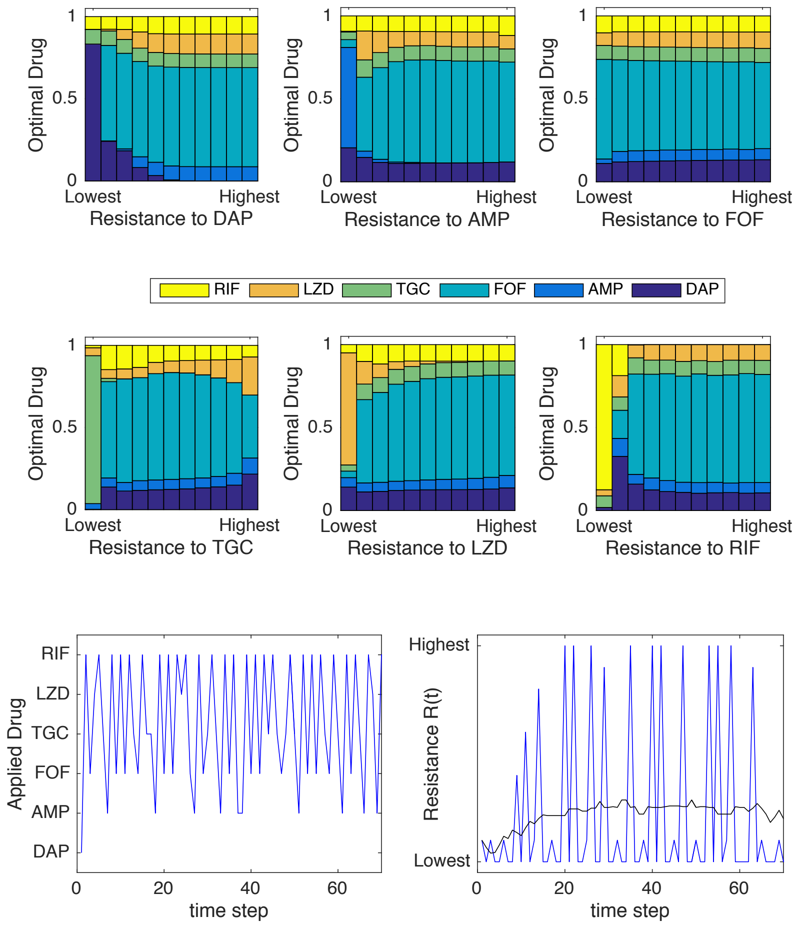


**FIG S8 Optimal policy statistics and sample trajectories for** $\gamma = 0.99$ **The optimal policy** $\pi^*(s)$ **is a mapping from the set of all possible resistance profiles (**$S$**) to the set of drugs (**$A$**). The policy associates each resistance profile with a unique (optimal) drug. Top panels: Frequency with which each drug is prescribed (according to the optimal policy) as a function of the level of resistance to an individual drug (horizontal axis). More specifically, for each of the six panels, the state space is partitioned into eleven distinct subsets, with each subset containing all states characterized by a given level of resistance to the particular drug in question (horizontal axis). The colored bars then show how frequently each of the six drugs is prescribed (according to the optimal policy) across all states within that subset. Bottom left panel: single simulated trajectory showing drug choice over time. Bottom right panel: single simulated trajectory of the instantaneous reward** $R$**, which corresponds to the resistance level to the applied drug. Blue curve is the specific trajectory; black curve is a moving average of the trajectory with a window size of 20.**

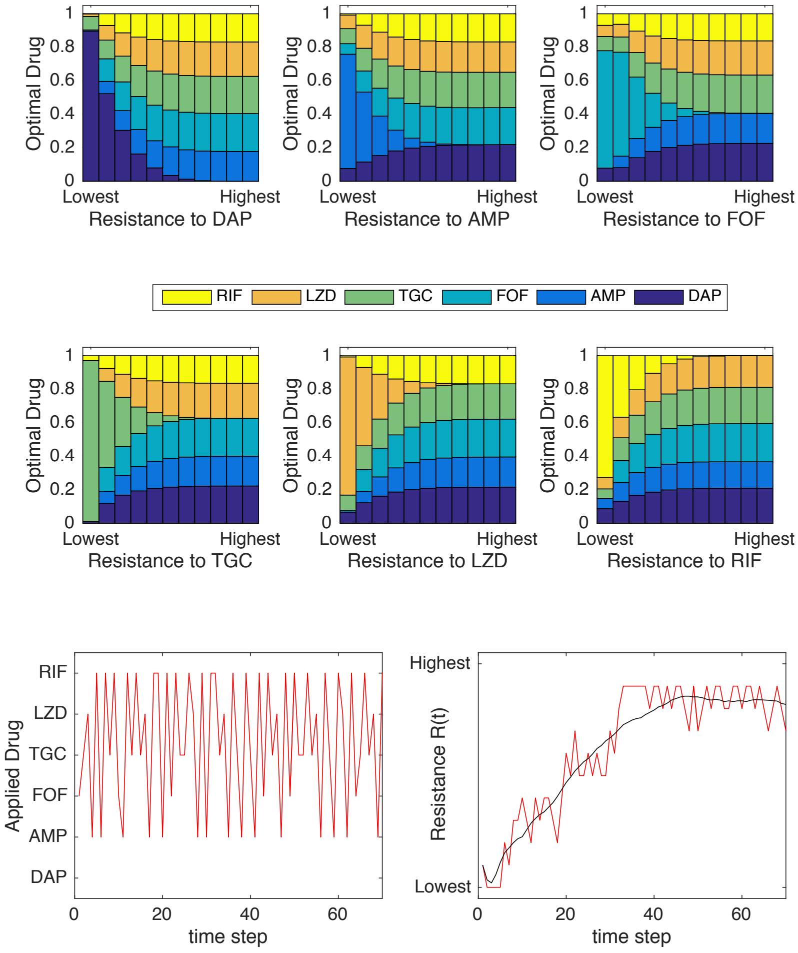


**FIG S9** **Optimal policy statistics and sample trajectories for** $\gamma = 0.1$ **Top panels: Frequency with which each drug is prescribed (according to the optimal policy) as a function of the level of resistance to an individual drug (horizontal axis). In each of the six panels, the state space is partitioned into eleven distinct subsets, with each subset containing all states with a given level of resistance to the particular drug in question. The colored bars then show how frequently each of the six drugs is prescribed across all states within that subset. Bottom left panel: single simulated trajectory showing drug choice over time. Bottom right panel: single simulated trajectory of the instantaneous reward** $R$**, which corresponds to the resistance level to the applied drug. Red curve is the specific trajectory; black curve is a moving average of the trajectory with a window size of 20.**

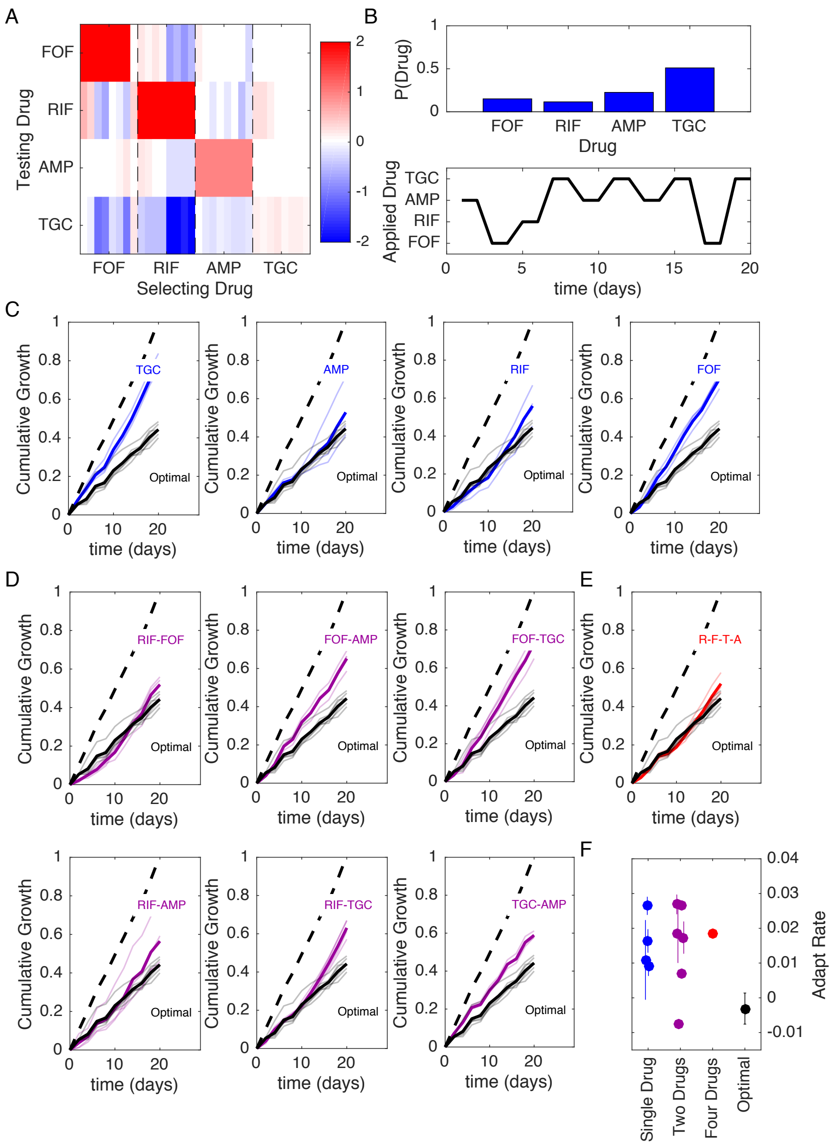


**FIG S10** **Optimized drug sequences reduce cumulative growth and adaptation rates in lab evolution experiments. A. Resistance (red) or sensitivity (blue) of each evolved mutant (horizontal axis; 4 drugs x 8 mutant per drug) to each drug (vertical axis) following 2 days of selection is quantified by the $\log_2$-transformed relative increase in the $IC_{50}$ of the testing drug relative to that of wild-type (V583) cells. B. Top: distribution of applied drug at time step 20 (approximate steady state) calculated using an optimal policy with $\gamma = 0.78$. Bottom: sequence of applied drug from one particular realization of the stochastic process with the optimal policy ($\gamma = 0.78$). C-E. Cumulative population growth over time for populations exposed to single drug sequences (C, blue), two-drug sequences (D,magenta), a four drug sequence (E, red), or the optimal sequence from panel B (black curves, all panels). Transparent lines represent individual replicate experiments and each thicker dark line corresponds to a mean over replicates. Dashed line, drug-free control (normalized to a growth of 1 at the end of the experiment). F. Adaptation rate for single drug (blue), two-drug (magenta), four drug (red), and optimal sequences (black). Error bars are standard errors across replicates. Adaptation rate is defined as the slope of the best fit linear regression describing time series of daily growth (see Figure S14).**

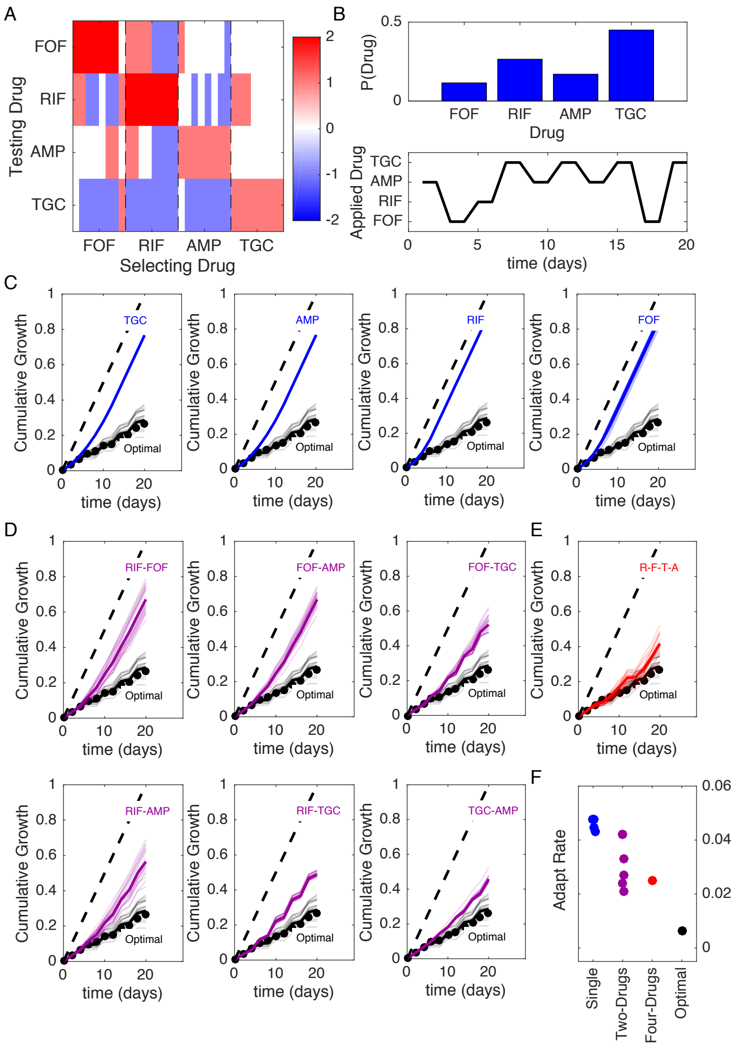

**FIG S11** **Optimized drug sequences reduce cumulative growth and adaptation rates in numerical simulations of the laboratory evolution experiments. Compare to Figure 6 (main text). A. Resistance (red) or sensitivity (blue) of each evolved mutant (horizontal axis; 4 drugs x 8 mutant per drug) to each drug (vertical axis) following 2 days of selection is quantified by the $\log_2$-transformed relative increase in the $IC_{50}$ of the testing drug relative to that of wild-type (V583) cells. The profile is then discretized into 4 levels of resistance. B. Top: distribution of applied drug at time step 20 (approximate steady state) calculated using an optimal policy with $\gamma = 0.9$. Bottom: sequence of applied drug from one particular realization of the stochastic process with the optimal policy ($\gamma = 0.9$). C-E. Cumulative population growth (simulations) over time for populations exposed to single drug sequences (C, blue), two-drug sequences (D,magenta), a four drug sequence (E, red), or the optimal sequence from panel B (black curves, all panels). Black circles correspond to the true optimal (i.e. applying the MDP policy directly) and performs only slightly better, on average, than the fixed sequence in panel B. At each time step, resistance level to each drug is converted to an OD value using a linear conversion with the highest resistance level corresponding to growth of drug-free cells (OD$\approx 0.6$) and the lowest resistance level corresponding to OD=0. Transparent lines represent individual replicate experiments and each thicker dark line corresponds to a mean over replicates. Dashed line, drug-free control (normalized to a growth of 1 at the end of the experiment). F. Adaptation rate for single drug (blue), two-drug (magenta), four drug (red), and optimal sequences (black). Error bars are standard errors across replicates. Adaptation rate is defined as the slope of the best fit linear regression describing time series of daily growth.**

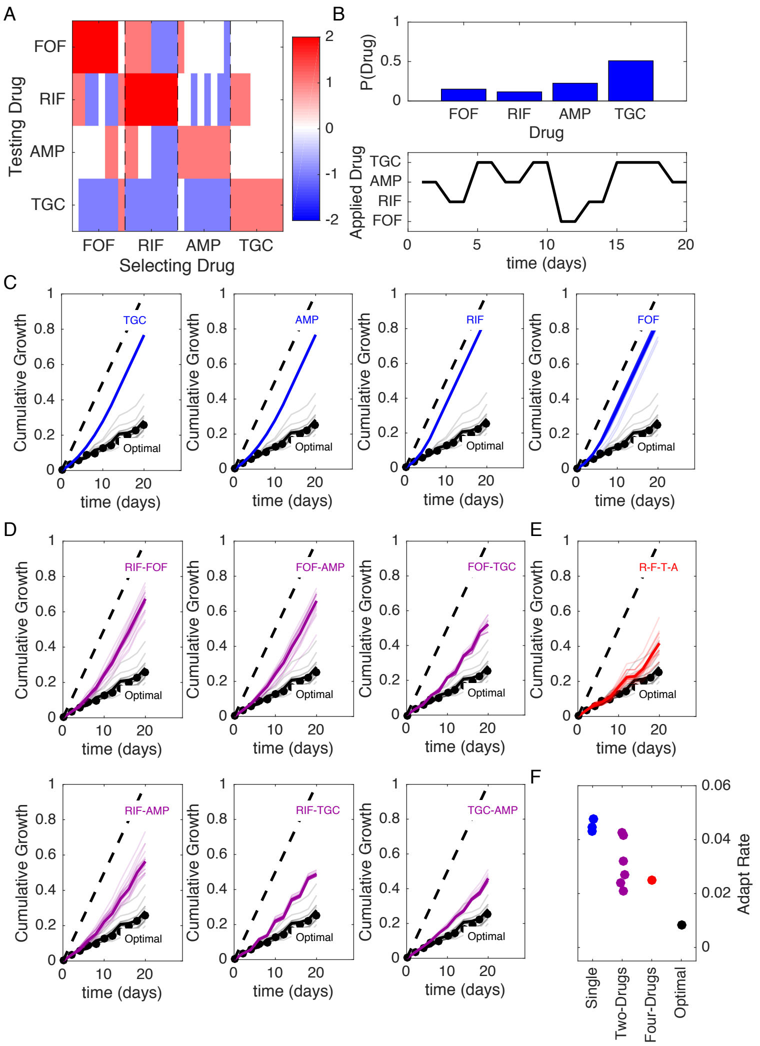


**FIG S12** **Optimized drug sequences reduce cumulative growth and adaptation rates in numerical simulations of the laboratory evolution experiments. Compare to Figure S10. A. Resistance (red) or sensitivity (blue) of each evolved mutant (horizontal axis; 4 drugs x 8 mutant per drug) to each drug (vertical axis) following 2 days of selection is quantified by the $\log_2$-transformed relative increase in the $IC_{50}$ of the testing drug relative to that of wild-type (V583) cells. The profile is then discretized into 4 levels of resistance. B. Top: distribution of applied drug at time step 20 (approximate steady state) calculated using an optimal policy with $\gamma = 0.78$. Bottom: sequence of applied drug from one particular realization of the stochastic process with the optimal policy ($\gamma = 0.78$). C-E. Cumulative population growth (simulations) over time for populations exposed to single drug sequences (C, blue), two-drug sequences (D,magenta), a four drug sequence (E, red), or the optimal sequence from panel B (black curves, all panels). Black circles correspond to the true optimal (i.e. applying the MDP policy directly) and performs only slightly better, on average, than the fixed sequence in panel B. At each time step, resistance level to each drug is converted to an OD value using a linear conversion with the highest resistance level corresponding to growth of drug-free cells (OD$\approx 0.6$) and the lowest resistance level corresponding to OD=0. Transparent lines represent individual replicate experiments and each thicker dark line corresponds to a mean over replicates. Dashed line, drug-free control (normalized to a growth of 1 at the end of the experiment). F. Adaptation rate for single drug (blue), two-drug (magenta), four drug (red), and optimal sequences (black). Error bars are standard errors across replicates. Adaptation rate is defined as the slope of the best fit linear regression describing time series of daily growth.**

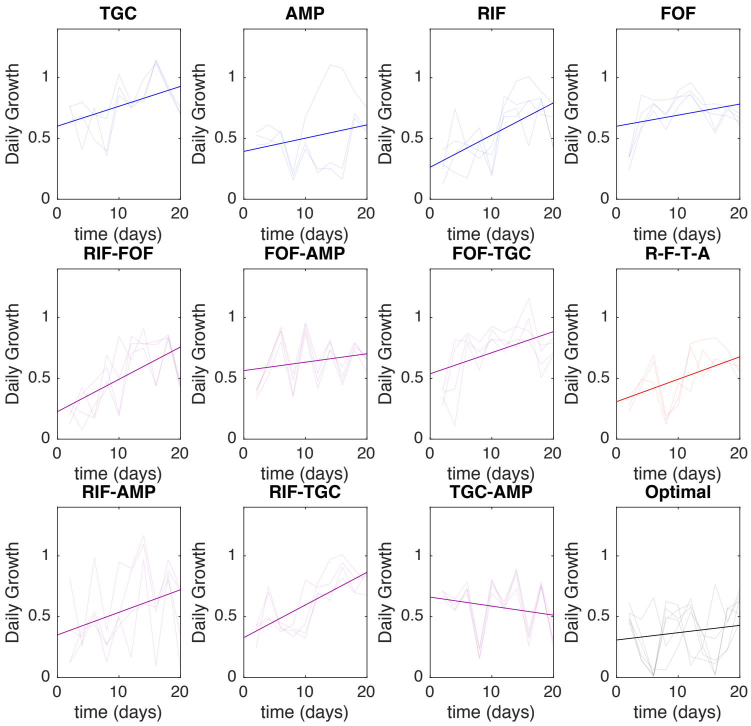


**FIG S13** **Estimated adaptation rate in lab evolution experiments based on γ=0.9 MDP policy. Daily growth, which is defined as the OD measured at the end of each 48-hour period (normalized to drug-free control), for populations exposed to single drug (blue), two-drug (magenta), four-drug (red), and optimal (black) drug sequences. All time series start at day 2 (i.e. following 48 hours of adaption). Transaprent curves correspond to individual replicate experiments; solid dark lines show the (average) best fit linear regression. Adaptation rate is defined as the slope of the regression line.**

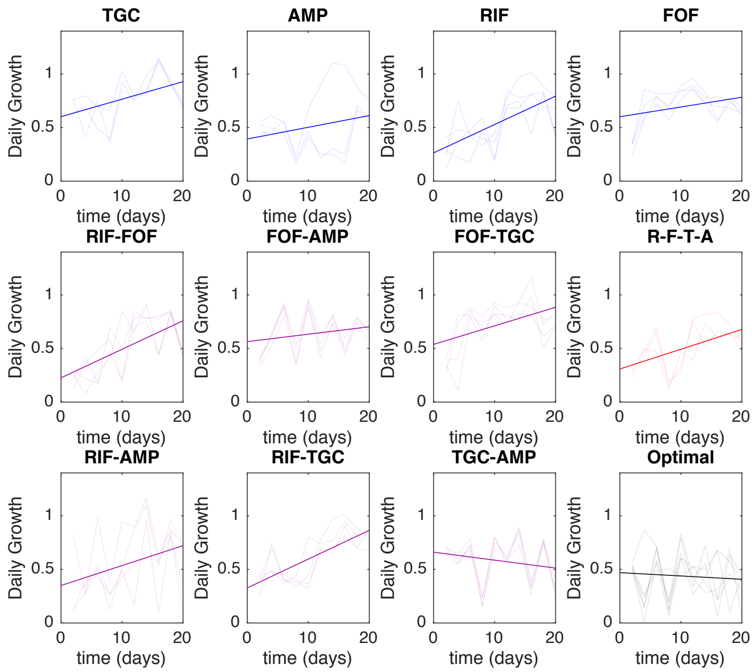


**FIG S14** **Estimated adaptation rate in lab evolution experiments based on γ=0.78 MDP policy. Daily growth, which is defined as the OD measured at the end of each 48-hour period (normalized to drug-free control), for populations exposed to single drug (blue), two-drug (magenta), four-drug (red), and optimal (black) drug sequences. All time series start at day 2 (i.e. following 48 hours of adaption). Transparent curves correspond to individual replicate experiments; solid dark lines show the (average) best fit linear regression. Adaptation rate is defined as the slope of the regression line.**
